# An empirical model of aminoacylation kinetics for *E. coli* class I and II aminoacyl tRNA synthetases

**DOI:** 10.1101/2024.09.26.615143

**Authors:** Eric Charles Dykeman

## Abstract

Efficient functioning of the prokaryotic translational system depends on a steady supply of aminoacylated tRNAs to be delivered to translating ribosomes via ternary complex. As such, tRNA synthetases play a crucial role in maintaining efficient and accurate translation in the cell, as they are responsible for aminoacylating the correct amino acid to its corresponding tRNA. Moreover, the kinetic rate at which they perform this reaction will dictate the overall rate of supply of aminoacylated tRNAs to the ribosome and will have consequences for the average translational speed of ribosomes in the cell. In this work, I develop an empirical kinetic model for the 20 aminoacyl tRNA synthetase enzymes in *E. coli* enabling the study of the effects of tRNA charging dynamics on translational efficiency. The model is parametrised based on *in vitro* experimental measurements of substrate *K*_*m*_ and *k*_*cat*_ values for both pyrophosphate exchange and aminoacylation. The model also reproduces the burst kinetics observed in class I enzymes and the transfer rates measured in single turnover experiments. Stochastic simulation of *in vivo* translation shows the kinetic model is able to support the tRNA charging demand resulting from translation in exponentially growing *E. coli* cells at a variety of different doubling times. This work provides a basis for the theoretical study of the amino acid starvation and the stringent response, as well as the complex behaviour of tRNA charging and translational dynamics in response to cellular stresses.

**Author summary:** Elucidating the complex interplay between tRNA charging by aminoacyl tRNA synthetases and the overall ribosomal demand for tRNAs will have important consequences for understanding the effects of amino acid starvation and the stringent response. Here I introduce an empirical kinetic model of the 20 *E. coli* tRNA synthetases and examine tRNA charging dynamics during exponential growth. The results show that the model is in good agreement with a variety of experimental observations, such as tRNA charging fractions, average translational speed of the ribosome, and measured total cellular tRNA abundances.

## Introduction

Understanding the functioning of the translational machinery in bacteria has the potential to impact a wide variety of fields in microbiology, such as understanding the effects of antibiotics which target ribosomes or aminoacyl tRNA synthetases, or in the development of efficient cell free protein synthesis systems. A key aspect of the protein translational apparatus of all cells is the aminoacylation of tRNAs, where amino acids are covalently linked to tRNAs for subsequent delivery to translating ribosomes. This action, sometimes referred to as tRNA charging, is performed via one of the 20 aminoacyl tRNA synthetases which are responsible for ensuring accuracy in the translation of the genetic code by covalently linking the correct amino acid to the correct tRNA. However, in order to theoretically and computationally study how tRNA synthetases affect the translational process, the creation of a model of tRNA charging by tRNA synthetases is required.

Since roughly the mid 1960’s, there has been tremendous work on elucidating the kinetic mechanism of tRNA charging by aminoacyl tRNA synthetases (AARS). Pioneering experimental work on AARSs started with the examination of their pyrophosphate exchange kinetics [1]. Using radio labelled pyrophosphate, researchers monitored the conversion of radioactive pyrophosphate into radioactive ATP by AARSs which could be isolated using thin layer chromotography. Later work began to look at both pyrophosphate exchange kinetics and the overall aminoacylation reaction by radio labelling the amino acid and monitoring the production of radioactive aa-tRNAs [2]. These experiments started to provide data on the kinetic behaviour of different AARSs, specifically the *k*_*cat*_ and *K*_*m*_ values of the three different substrates (amino acid, ATP, and tRNA) along with details of the overall steady-state kinetics. Subsequent kinetic experiments examined the behaviour of several AARSs under single-turnover conditions (i.e. where enzyme is in excess over tRNA), allowing estimations of the kinetic rates of amino acid transfer to the tRNA (*k*_*tran*_) and of the overall chemistry step (*k*_*chem*_), which is a composite rate that includes the formation of amino acid adenylate complex along with the transfer step and formation of aminoacylated tRNA [3].

This work, together with structural data of AARSs in complex with various substrates, has allowed a fairly descriptive picture to emerge of the kinetic steps which take place on these enzymes and a classification into two main classes based on their observed kinetic properties [4, 5]. Class I AARSs are mostly monomeric and usually display burst kinetics of aminoacyl tRNA (aa-tRNA) formation, while Class II AARSs are dimers and display no burst kinetics. Here burst kinetics describes a property of the enzyme in which there is an initial burst of aa-tRNA production at a rate greater then the *k*_*cat*_ of the enzyme followed by the steady state aa-tRNA production rate. Supplementary Table A summarises the classes for each of the AARS enzymes in *E. Coli*.

While experimental measurements of AARS enzymes have had sufficient time to develop over the last 50-60 years, there has been limited attempts to computationally model, or reproduce empirically, the experimental observations of AARS kinetics. Airas [6, 7] used experimental observations of tRNA charging kinetics for class I argRS and ileRS to estimate a set of best fit kinetic parameters for a theoretical model. However, it is unclear if these models reproduce burst kinetics for these enzymes. On the other hand, Santra and Bagchi [8] provide ODE models of tRNA charging kinetics for both class I and class II AARS enzymes where their kinetic models are able to demonstrate burst and non-burst kinetics. In this work, the authors consider product release as the rate limiting step in a model of Class I cysRS kinetics, which reproduced the burst kinetics reported in Zhang *et al*. [9]. For class II enzymes, they introduce an amino acid activation pathway in the presence of tRNA which was much slower than the activation in the absence of tRNA [8]. This resulted in steady state kinetics that displayed no pre-steady state burst. However, despite the success of their model in reproducing burst kinetics, it is unclear if their models would accurately reproduce experimentally observed *K*_*m*_ measurements for the substrates.

More recently, Choi and Covert have attempted to develop a Michaelis–Menten model for all 20 *E. coli* aminoacyl tRNA synthetases [10]. Their model is developed based on a review of the experimental literature of *in vitro k*_*cat*_ and *K*_*m*_ measurements, which have been subsequently optimised to support the translation speed required to double the *E. coli* proteome in a model of the *E. coli* cell cycle [11]. While such a model should potentially reproduce experimentally observed *K*_*m*_ measurements for the substrates, it ignores subtle features of the AARS enzymes such as burst kinetics.

The goal of this work is to develop an empirical kinetic model, similar to the models of Airas and/or Santra and Bagchi, which details the individual kinetic steps of aminoacylation reaction whilst also reproducing burst kinetics and experimentally observed *K*_*m*_ measurements for the substrates. The development of an empirical model of AARS kinetics can be useful for several reasons. First, in a bacterial cell such as *E. coli*, several important questions remain regarding tRNA charging and turnover by AARSs during ribosome elongation on messenger RNA (mRNA). One such question relates to the effect on tRNA charging during amino acid starvation and the activation of the stringent response. Work by Elf *et al*. examined a simple model of tRNA charging competition between different tRNA isoacceptors after amino acid starvation [12, 13]. However, a more detailed model that also accounts for *K*_*m*_ values of amino acid can also look at, for example, regimes just at the threshold of starvation, where there may also be subtle effects on the charging levels of tRNAs [13].

The paper is outlined as follows. First, I create an empirical kinetic model for all 20 aminoacyl tRNA synthetases of *E. coli* which recapitulate their observed kinetic properties *in vitro*. Here, the observed kinetic properties are their *k*_*cat*_, *K*_*m*_, and single turnover rates *k*_*tran*_ and *k*_*chem*_. Second, using these models combined with proteomics data for *E. coli* growing under controlled chemostat conditions in the exponential phase, I demonstrate that the observed *in vitro* kinetic rates (*k*_*cat*_ and *K*_*m*_) are roughly in line with the *k*_*cat*_ values that would be required to supply charged tRNAs to elongating ribosomes *in vivo*, with a few enzymes requiring a small adjustment (by a factor of 2 on average) to their *k*_*cat*_. This is in contrast to the recent model of Choi and Covert [10] where the Authors report that the majority of enzymes require, on average, a 7-fold increase in their *k*_*cat*_ values to support observed *in vivo* charging rates. Finally, I discuss some of the issues in creating a Michaelis–Menten model of tRNA synthetases, and compare my results with the recent model reported by Choi and Covert [10].

## Methods

### Aminoacyl tRNA synthetase kinetic model construction

Development of an empirical model of AARS kinetics requires two main steps; first a description of the individual kinetic steps involved in the charging of tRNAs with amino acids, and second, a method for fitting the kinetic rates such that the experimentally observed kinetic properties of the enzyme are reproduced by the model. Amino acid charging of tRNAs by aminoacyl tRNA synthetases occurs via a two step mechanism consisting of amino acid activation (Eq. 1) and amino acid transfer (Eq. 2), as described below. It is critically important to note that both of these steps are catalytic events and hence each have their own *k*_*cat*_ and *K*_*m*_ values for the reactions. In the first step, amino acid and ATP bind to the enzyme, forming an amino acid adenylate (AMP-aa), followed by release of pyrophosphate (activation). The amino acid adenylate reaction can occur with or without tRNA present on the enzyme, except in the three class I enzymes argRS, gluRS and glnRS, which specifically require tRNA to be present for activation to occur. In the second step, the amino acid is transferred from the adenylate to the tRNA to form the charged aa-tRNA (transfer). While the basic reaction scheme of the two-step process is described by Eqs. 1 and 2, this does not form a complete kinetic picture of the individual reactions (binding of ATP, dissociation of pyrophosphate *etc*.) which occur, nor the kinetic order of these events.

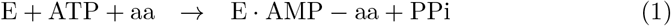

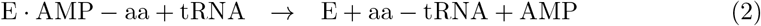

In the next sections I give details on the individual kinetic steps that I consider in my model for both monomeric class I, dimeric class I, and dimeric class II AARS enzymes. In addition, I discuss the mathematical procedures used to calculate observed kinetic properties of the enzyme, (*i*.*e. k*_*cat*_, *K*_*m*_, *k*_*chem*_, *k*_*tran*_), and describe the general steps used to identify kinetic parameters for the model. Additional discussion of more technical information regarding the search of the multi-dimensional kinetic parameter space can be found in supplementary S1 Text.

### Kinetic reaction scheme for monomeric class I tRNA synthetases

For the monomeric class I aminoacyl tRNA synthetases (cysRS, argRS, valRS, ileRS, leuRS, glnRS, and gluRS), I use the reaction scheme depicted in Figure 1a to model the full aminoacylation reaction. I also model the class I dimers tyrRS and trpRS using this scheme since tyrRS was shown to have half of sites activity [14, 15] while trpRS was shown to bind tRNA asymmetrically across the dimer interface [16]. The labelled states *S*_*i*_ with *i*∈ [0, 15] represent different states of the AARS enzyme, with *S*_0_ representing the AARS enzyme free of all substrates. Specific details on each state of the enzyme, along with the labelled kinetic rates, are given in the Supplementary Figure C. All of the labelled kinetic rates in the diagram have both a forward and a backward rate (e.g. *k*_2*f*_ and *k*_2*b*_), and I label dissociation constants for substrate binding reactions as *K*_2*d*_ = *k*_2*b*_*/k*_2*f*_ . Only the general kinetic binding and chemical reaction steps are considered, and any conformational changes of the enzyme itself are ignored in the kinetic reaction scheme. Any editing reactions where the enzyme checks for mis-charged tRNAs are also not considered in this kinetic model.

**Fig 1.**
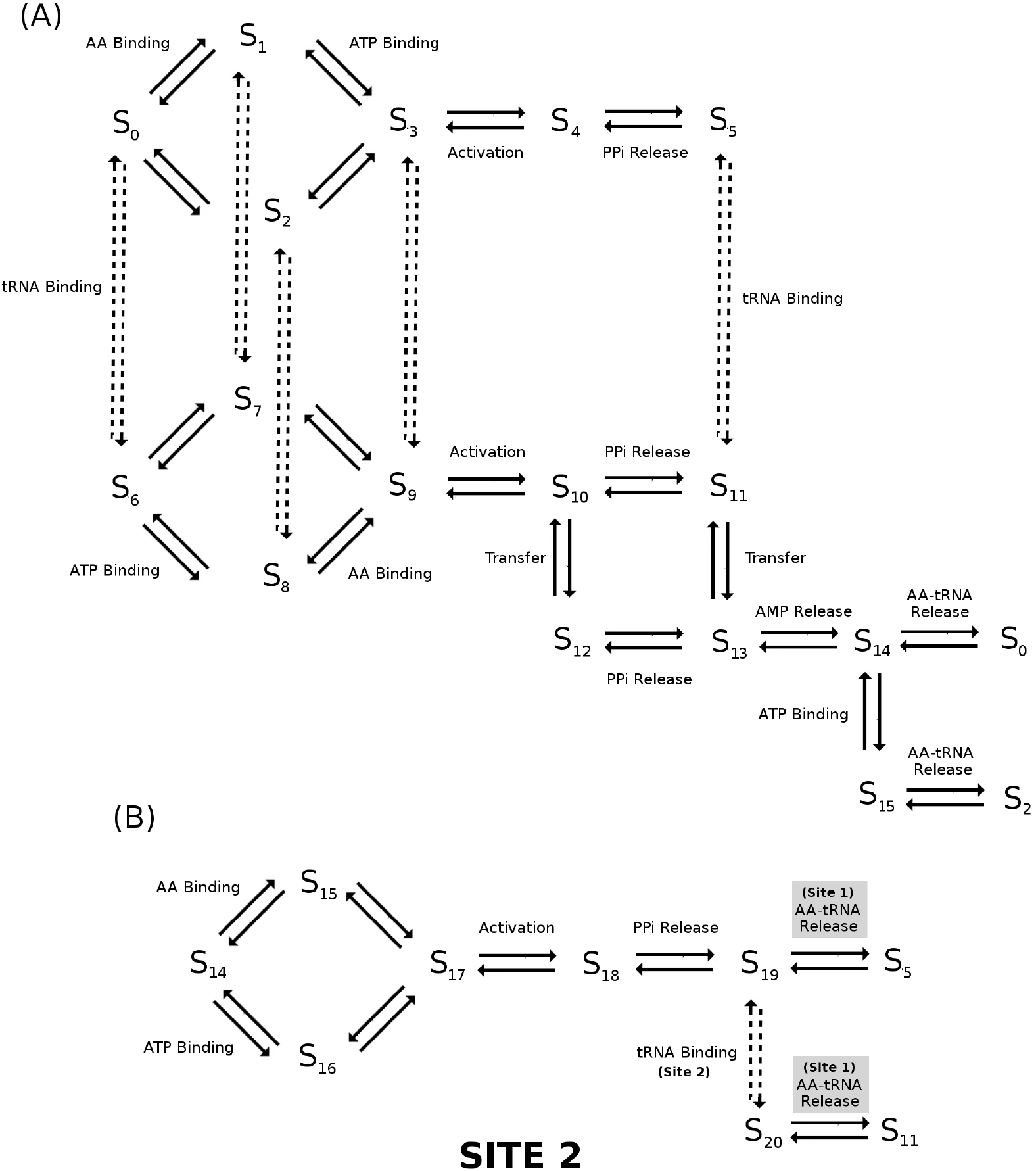
Kinetic reaction scheme for class I and class II tRNA synthetases. (A) Detailed map of the reaction pathway for monomeric class I AARS enzymes. State *S*_0_ corresponds to the enzyme free of all substrates. The two main catalytic events, activation of the amino acid to form amino acid adenylate plus pyrophosphate (activation), and transfer of the amino acid from the adenylate to the tRNA (transfer) are labelled. (B) The reaction pathway for class I dimer metRS and class II AARS enzymes follows the kinetic flip-flop mechanism reported in Guth *et al*. [17]. Here, the same kinetic steps in site 1 of the dimer occur as in the class I enzymes followed by a subsequent activation event in site 2 while charged aa-tRNA in site 1 remains bound. After activation in site 2, aa-tRNA is released from site 1.

While the reaction scheme may seem overly complicated, all of these steps are required in order for the model to reproduce *K*_*m*_ values across both pyrophosphate exchange and aminoacylation experiments. Previous work on a detailed experimental analysis of isoleucine tRNA synthetase kinetics from *S. aureus* by Pope and colleagues [18] has shown that in the presence of tRNA, amino acid, and ATP, all three substrates bind to the enzyme in a random order before undergoing adenylate formation and aa-tRNA formation. As such, I have allowed the random binding of these substrates to occur and have also allowed adenylate to form with or without tRNA present (except for argRS, gluRS, and glnRS). It is interesting to note that if the kinetic steps of the model are simplified to consider an ordered binding of tRNA, amino acid, and ATP to the enzyme, then the predicted *K*_*m*_ for tRNA in the aminoacylation reaction is often drastically lower (usually by a factor of 10 or more) than what is measured experimentally. However, if a random binding of the three substrates is considered, as shown in Figure 1A, then the predicted *K*_*m*_ for tRNA can be easily parameterised to be within the range of *K*_*m*_ values measured experimentally. Additionally, when tRNA is bound to the enzyme, I have also allowed for pyrophosphate release to occur either before or after transfer of the amino acid to the tRNA (path *S*_10_ → *S*_11_ → *S*_13_ vs. path *S*_10_ → *S*_12_ → *S*_13_). Without this dual pathway, *K*_*m*_s for ATP and amino acid in the aminoacylation reaction are often drastically lower than what is measured experimentally. This same problem was also noticed by Pope and colleagues when constructing a kinetic model of ileRS from *S. aureus* [18] which caused them to propose a similar mechanism of amino acid transfer prior to pyrophosphate release. Moreover, several other experimentalists have postulated that pyrophosphate may stay bound during the amino acid transfer step in some AARSs [19]. Finally, the reaction path *S*_14_ →*S*_15_ →*S*_0_ in which ATP binds to the enzyme prior to the release of the aa-tRNA product ensures that the AARS enzyme is not kinetically inhibited in the presence of cellular concentrations of AMP (estimated 125-250 *µ* M [20]) and/or pyrophosphate (estimated 250-500 *µ* M [21, 22]).

### Kinetic reaction scheme for dimeric class II tRNA synthetases

For the dimeric class II AARS enzymes (serRS, thrRS, proRS, hisRS, aspRS, asnRS, lysRS, alaRS, glyRS and pheRS), as well as the dimeric class I metRS, I use the reaction scheme depicted in Figures 1A and B to model the aminoacylation reaction. Here, I assume that the dimeric enzymes follow the flip-flop kinetic model as described in Guth et al. [17] for hisRS. The flip-flop kinetic model assumes amino acid adenylate is formed and transferred to a tRNA at site 1 of the enzyme following the kinetic steps of a class I enzyme (Figure 1A). Afterwards, the charged tRNA remains bound to site 1 while an additional amino acid adenylate is formed at site 2 as shown in Figure 1B. Subsequently, the charged tRNA is released from site 1 while an uncharged tRNA is recruited to site 2. I allow the charged tRNA at site 1 to be released after activation of the amino acid at site 2, regardless of whether tRNA is bound at site 2.

### Identification of kinetic parameters for the model

For the reaction scheme of class I AARS in Figure 1A, there are 48 kinetic parameters that must be chosen, with the goal that the resulting kinetics predicted by the model (specifically the *K*_*m*_ and *k*_*cat*_ for the enzymes substrates) match experimental observations. At first, the prospect of identifying a set of 48 parameters which will reproduce experimental values of *K*_*m*_ and *k*_*cat*_ seems impossible, even with experimental input. However, the complexity of fitting the parameters can be drastically reduced if one performs the fitting in three separate stages and allows some parameters, such as the rate of formation of the adenylate in the presence or absence or tRNA, to be identical with each other. Moreover, tRNA, amino acid, and ATP have usually had their dissociation constants (*K*_*d*_) to AARS enzymes measured, which allows for further complexity reductions.

The specific idea that I propose to use to identify the kinetic parameters has two parts. The first is to exploit the fact that experimentalists perform three separate assays to elucidate the kinetic properties of AARSs; (1) the pyrophosphate exchange assay, (2) the single turnover kinetics assay, and (3) the aminoacylation assay. While the aminoacylation assay examines the entire process (and thus depends on all 48 kinetic parameters), the pyrophosphate exchange and single turnover kinetics only examine a subsection of the reaction scheme and thus only depend on a subset of the kinetic parameters. Figure 2A illustrates the kinetic scheme for the pyrophosphate exchange (12 kinetic parameters), while 2B shows the scheme for the single turnover assay measuring the transfer rate *k*_*tran*_ (8 kinetic parameters). Thus, by fitting first the single turnover reaction, followed by the pyrophosphate exchange, one can slowly constrain parameters resulting in a smaller number to be varied in the final fitting of the aminoacylation assay. The second part relies on the observation that only a few of the kinetic parameters have an effect on the *K*_*m*_ or *k*_*cat*_ of the enzyme. For example, while the pyrophosphate exchange assay depends on 12 kinetic parameters, it can be shown through systematic variation of pairs of parameters that only 4 have a major effect on *K*_*m*_ or *k*_*cat*_ (see supplementary Figure B and discussion in S1 Text). While this implies that one can have a free choice for the other parameters, this is not how the fitting is done. Instead, the remaining parameters that have little or no effect on *K*_*m*_ or *k*_*cat*_ are chosen based on experimental measurements.

**Fig 2.**
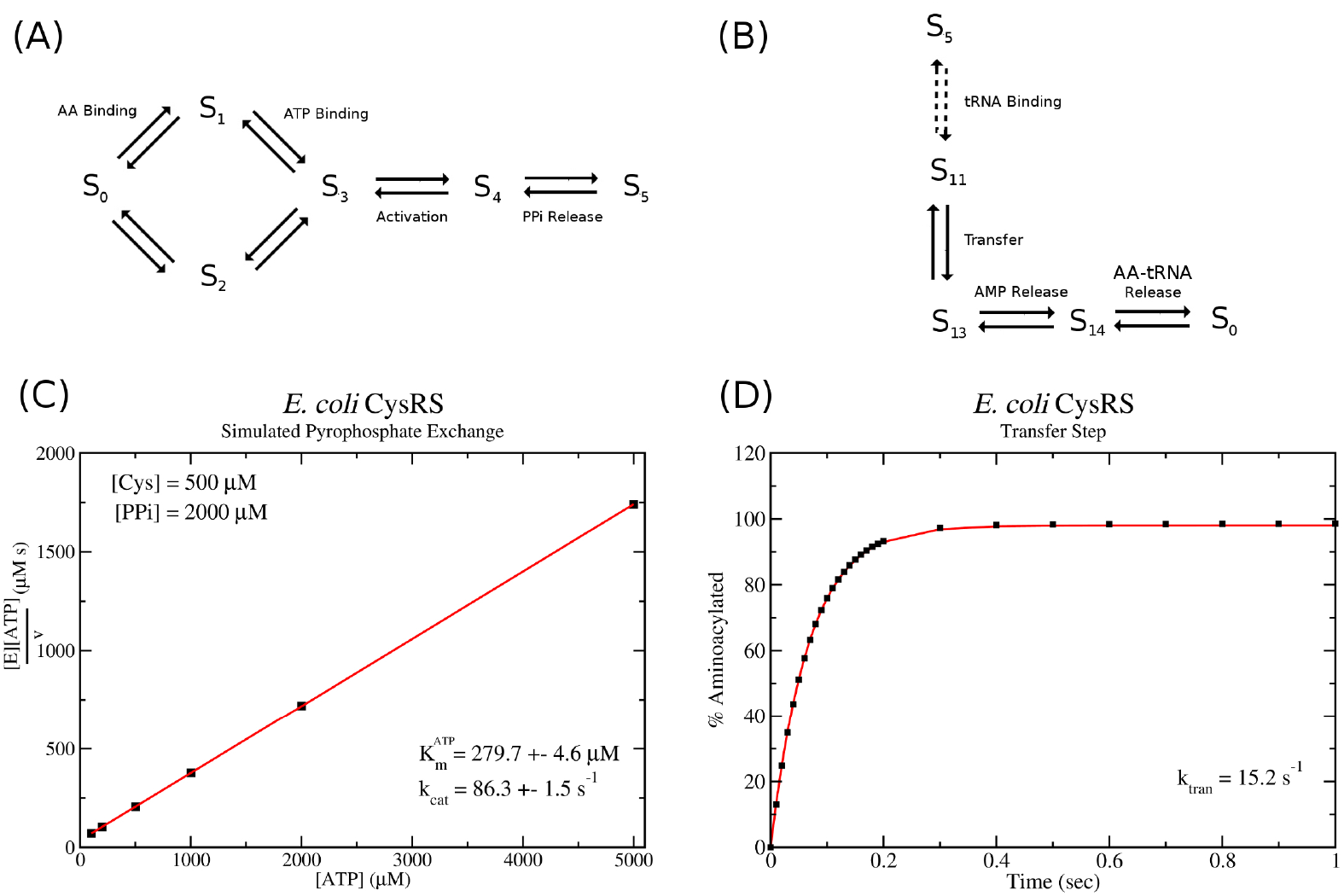
Computer simulation of pyrophosphate exchange and single turnover kinetics. (A) Reaction scheme used to simulate pyrophosphate exchange kinetics. (B) Reaction scheme used to simulate single turnover kinetics involving the transfer step. (C) Example Woolf-Hanes plot and the resulting values of 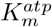 and *k*_*cat*_ from a computer simulation of pyrophosphate exchange for *E. coli* cysRS. (D) Example plot from a computer simulation of single turnover kinetics for *E. coli* cysRS. Black squares represent data points from the computer simulation while the red curve gives the best fit exponential *y*(*t*) = *A*(1 − exp(−*k*_*tran*_*t*)) to the data.

Parameters are varied following the algorithm in S1 Text until the kinetic properties of the enzyme predicted by the model (*k*_*cat*_, *K*_*m*_ for each substrate, and *k*_*trans*_, *k*_*chem*_) match target values. These target values are typically the average of experimental data measurements, with outliers removed. The result is a set of parameters that reproduce the observed *K*_*m*_ and *k*_*cat*_ values for the enzyme, as well as any other kinetic behaviours, such as burst kinetics for the class I enzymes.

### Calculation of *k*_*cat*_ and *K*_*m*_ in the kinetic model

As discussed above, there are three different experimental assays which examine different aspects of AARS kinetics. The first, pyrophosphate exchange, involves introducing radio labelled pyrophosphate to a mixture of ATP, amino acid, and enzyme with the reaction occurring according to the scheme shown in Figure 2A. ATP is then isolated by thin layer chromatography and its specific activity measured. Thus, this assay indirectly measures the rate of activation of the amino acid by measurement of the reverse process, *i*.*e*. the conversion of radioactive polyphosphate and adenylate back to radioactive ATP and amino acid. The assay measures *k*_*cat*_ and *K*_*m*_ for both ATP and amino acid by fixing the concentration of one at saturating conditions then varying the concentration of the other. Cole and Schimmel [23] showed that the concentration of radioactive ATP (denoted ATP*) can be written as

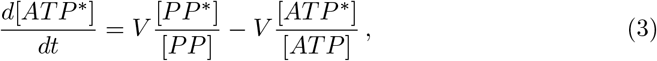

where *V* is the speed of the pyrophosphate exchange reaction. At *t* = 0 the amount of radioactive ATP is zero, and one obtains the relation for the initial speed of the conversion reaction as

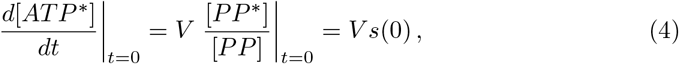

where *s*(*t*) is the specific activity of the pyrophosphate. Thus, one can directly simulate using either a stochastic or ODE based model the reaction scheme for pyrophosphate exchange using initial conditions matching experimental protocol. These initial conditions are usually [*PP* ] = 2 mM, [*ATP* ] = 5 mM, [*AA*] = 0.5 − 1 mM, with initial enzyme concentration [*E*] = 1 − 10 nM with a specific activity of *s* = 0.02. Varying either the initial ATP, PP, or amino acid concentrations while keeping the remaining substrates fixed, I simulate the pyrophosphate exchange scheme (Figure 2A) using the stochastic Gillespie algorithm. For each concentration of the varying substrate, the reaction kinetics are simulated 10 times, and the average and standard deviation of the initial reaction velocity *V* is computed. I then plot the velocity and concentration data using a Woolf-Hanes plot, which plots substrate concentration over the velocity versus substrate concentration. Linear regression is used to produce the best fit line (*y* = *mx* + *b*) and the slope of this line is *m* = 1*/k*_*cat*_ while the line intercept is *b* = *K*_*m*_*/k*_*cat*_. Figure 2C shows an example of a *K*_*m*_ and *k*_*cat*_ calculation resulting from a computer simulation of pyrophosphate exchange for *E. coli* cysRS. The procedure is automated such that a parameter search can be employed by calling a function that returns *k*_*cat*_ and *K*_*m*_ and their standard error given initial substrate concentrations and a set of kinetic parameters. A near identical procedure can be done for the aminoacylation reaction schemes (Figure 1) where the production rate of aa-tRNA is monitored. Typical initial concentrations for the aminoacylation reactions are [*ATP* ] = 5 mM, [*AA*] = 0.5 − 1 mM, [*tRNA*] = 10 *µ*M, and [*E*] = 1 − 10 nM.

For the single turnover kinetics assay, the computer simulation and experiment measurements are slightly different. Here, the assay first pre-forms enzyme bound with adenylate (state *S*_5_ in Figures 1 and 2B) and then isolates this enzyme complex. The adenylate contains a radio-labelled amino acid which will subsequently be transferred to a tRNA. The assay then proceeds with measuring this transfer rate under single turnover conditions, i.e. where concentration of the enzyme bound with radioactive adenylate is roughly 10x greater then the concentration of tRNA. Charged tRNAs are isolated at varying time points using thin layer chromatography and the data are fit to the exponential function,

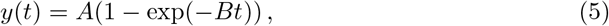

with *B* = *k*_*tran*_. In order to compare the model with this experiment, I simulate the tRNA charging reaction using the reaction scheme in Figure 2B and typical concentrations used in experiments ([*tRNA*] = 5 *µ*M, and [*E* : *Ad*] = 50 *µ*M). The formation of aa-tRNA as a function of time is output, and the results are fit to Eq. 5. A similar single turnover assay which mixes ATP, amino acid, tRNA and enzyme simultaneously with enzyme 10x tRNA concentration can be used to measure the overall chemistry step *k*_*chem*_. The inverse of *k*_*chem*_ approximates the average time required for the enzyme to bind all substrates, activate an amino acid, and transfer the amino acid to tRNA. Most single turnover assays observe that *k*_*chem*_ ≈ *k*_*tran*_ suggesting that binding of substrate and activation of the amino acid is fast compared with the final transfer step. Thus, *k*_*chem*_ can be used to confirm that the kinetic rates for substrate binding and amino acid activation in the model are sufficiently fast.

### Computational simulations of *in vivo* translation and tRNA charging

Previously I developed a stochastic model of *in vivo* prokaroytic translation which takes into account protein synthesis on the full transcriptome present in an *E. coli* cell [24, 25]. For example, in a typical *E. coli* cell which has a doubling time of *τ* = 60 min, the model accounts for the protein synthesis occurring by roughly 15000 ribosomes on a transcriptome of roughly 2 million nucleotides of mRNA. In addition, the model also accounts for competition between mRNAs for available ribosomes as well as the competition for the A-site on the ribosome between different tRNA isoacceptors. Finally, since the translational model [24] takes into account each of the known individual kinetic steps in the translation process, the resulting elongation rate of the ribosomes emerges as a result of the codon bias in the mRNA and the concentration of tRNAs in ternary complex. Thus my model is capable of predicting the resulting change to the ribosome elongation rate that would result from any alteration to tRNA charging rates. The base code used in this work can be downloaded from Github at github.com/edykeman/ribofold and kinetic parameters used for the ribosome elongation steps can be found in [24]. Predictions of tRNA charging kinetics by AARS enzymes are incorporated into the stochastic ribosome model from [24] using the kinetic scheme in Figure 1. The individual kinetic parameters used in the model can be found in Supplementary S1 Text, Tables B, C, and D. The translational model can also be run using a simplified Michaelis–Menten model for aminoacylation kinetics. Parameters used for the *k*_*cat*_ and *K*_*m*_ values are listed in Table I, with the optimized tRNA numbers for use with the Michaelis–Menten model given in Table J. Software used in the simulations can be downloaded from http://www-users.york.ac.uk/~ecd502/ or from Github at edykeman/ribosome-aars.

### Construction of *E. coli* transcriptome

To construct the transcriptome for the stochastic model of translation, I have used mRNA-seq measurements from Li *et al*. [26] which give experimental measurements of the overall average numbers of individual mRNAs (in RPKM units) in the transcriptome. This information can be used to back construct a snapshot of the average transcriptome that would be observed in exponentially growing *E. coli* cells. To do this, I normalise the RPKM values (*n*_*R*_(*i*)) for each mRNA type *i*, i.e.

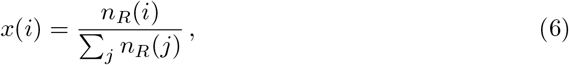

where the sum is over all mRNAs in the RNA-seq data. Then, I calculate the total number of mRNAs of each type via

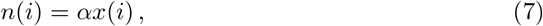

where *α* is a constant. The value *α* is chosen such that the total nucleotide content of the transcriptome approximately reproduces what has been estimated by Bremer and Dennis [27] for *E. coli* cells growing exponentially at growth rates of *µ* = 0.41, 0.69, 1.04, and 1.73 h^*−*1^, *i*.*e*. doubling times *τ* = 100, 60, 40, and 21 min. As a check, I calculate the codon bias in the predicted transcriptome and compare with the codon usage that was measured by a separate experiment reported in Dong *et al*. [28]. Supplementary S2 Spreadsheet Tab 3 shows that there is excellent agreement between the codon usage in the transcriptome versus what was measured in Dong *et al*.

### Construction of violin plots

Consider a set of *n* measurements *y*_*i*_ with *i* ∈ [1, *n*], each with an associated measurement error *σ*_*i*_. Using this data set, I construct a violin plot *p*(*y*) with *p* ∈ [0, 1] via

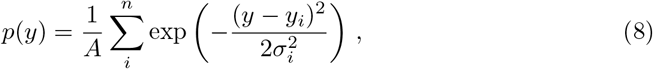

where the constant *A* is chosen such that

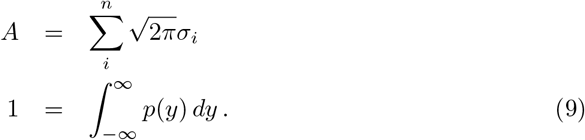

The function *p*(*y*) can thus be considered as a probability density, where input of a particular value *y* gives an estimate of the probability that an experiment would measure this value. In the case of the proteomics data points, where no error value *σ* is usually given, I use a default 20% error value for the measurement.

## Results

### Empirical kinetic model of *E. coli* tRNA synthetases

I begin by discussing the kinetic models that have been constructed using the procedure in Methods for class I cysRS and class II hisRS enzymes from *E. Coli*. Cystine tRNA synthetase is a monomeric class I enzyme which displays burst kinetics, while hisRS is a class II enzyme which displays no burst kinetics in the pre-steady state. The kinetic properties of both enzymes (*k*_*cat*_, *K*_*m*_ and *k*_*chem*_, *k*_*tran*_ from single turnover assays) have been extensively measured *in vitro* by several different groups.

Table 1 gives the *in vitro* measurements of *k*_*cat*_ and *K*_*m*_ values for *E. coli* cysRS. Using this data, consensus *K*_*m*_ and *k*_*cat*_ values for the pyrophosphate exchange and aminoacylation assays can be estimated and, using the procedure outlined in Methods, a set of kinetic parameters which reasonably reproduce these values can be determined. Specifically for pyrophosphate exchange, the kinetic model gives values of 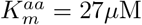, 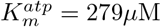, and *k*_*cat*_ = 85 s^-1^ Similarly for aminoacylation, the kinetic model gives values of 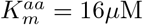, 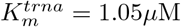,and *k*_*cat*_ = 2.96 s^*−*1^. Comparing with experiment, one can see that these values are within the range of those that have been measured experimentally (Figure 3A). Moreover, the identified parameters and kinetic model for cysRS reproduces the observed burst kinetics (Figure 3B) as well as the observed *k*_*tran*_ and *k*_*chem*_ values in simulations of single turnover kinetics (Figures 3C and 3D).

**Table 1.**
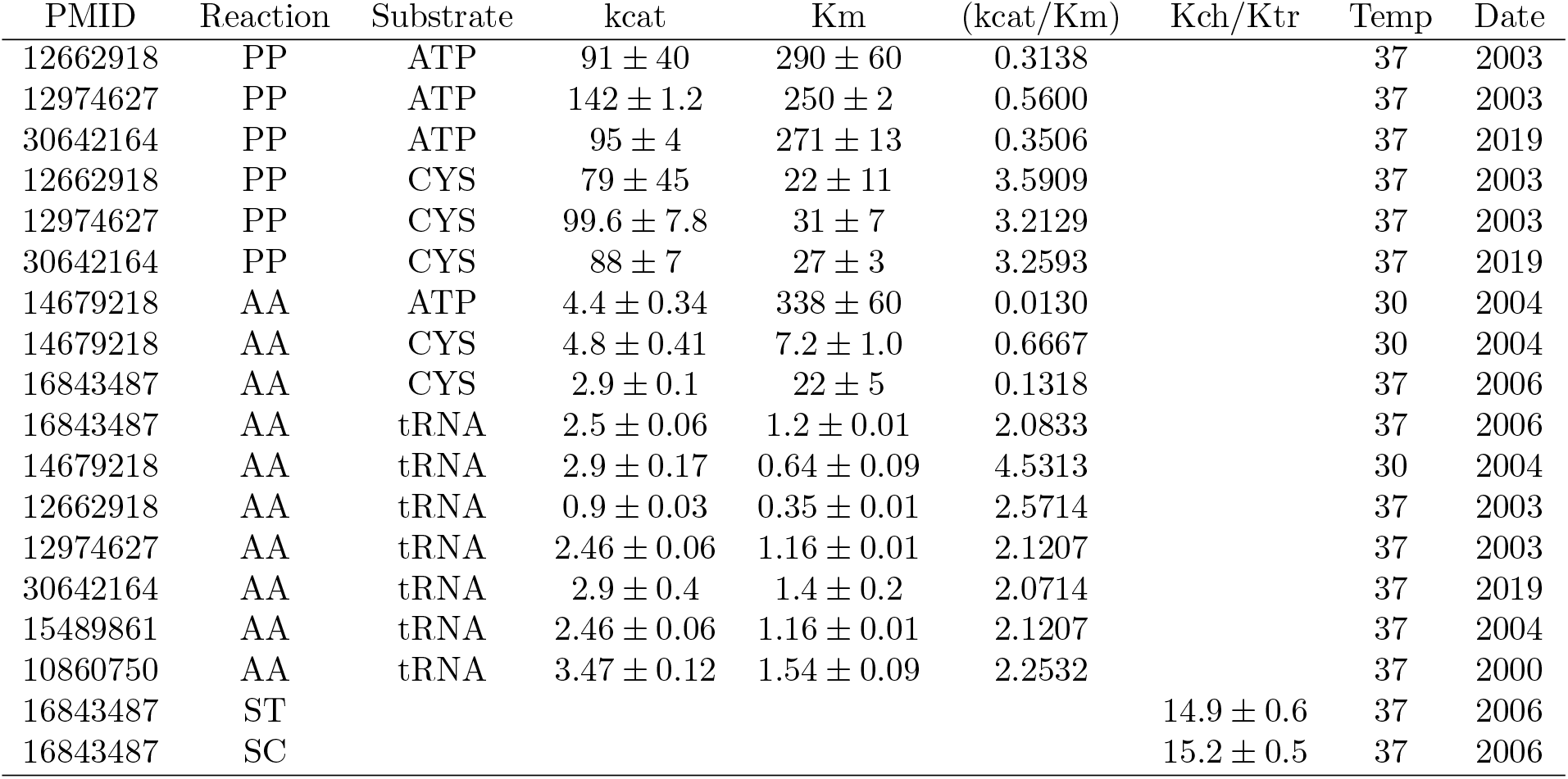
In vitro measurements of *E. coli* cysRS *K*_*m*_ and *k*_*cat*_ values. The experimental measurements for three separate kinetic assays are reported for the Class I cysRS aminoacyl-tRNA synthetase enzyme. Abbreviations are, PP = pyrophosphate exchange, AA = aminoacylation, ST = single turnover transfer, and SC = single turnover overall chemistry. Units for *k*_*cat*_ are in *s*^*−*1^ and *K*_*m*_ are in *µ*M.

**Fig 3.**
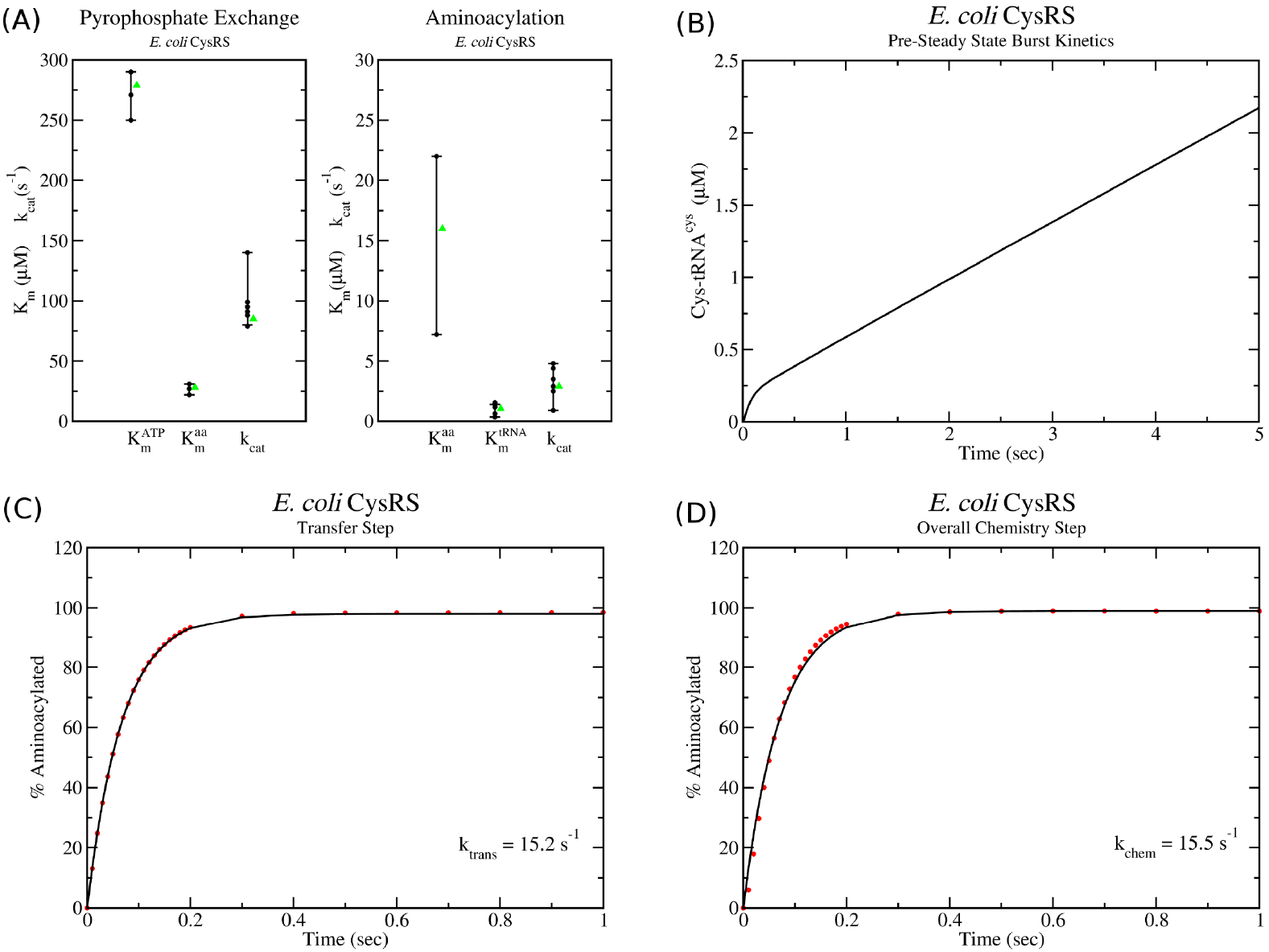
Kinetic model of class I cysRS tRNA charging. (A) Range of *k*_*cat*_ and *K*_*m*_ values determined by *in vitro* pyrophosphate and aminoacylation assays (black dots) are compared with the values of the kinetic model (green triangles). (B) Pre-steady state kinetics simulation with starting concentrations of [*cys*] = 500*µ*M, [*ATP* ] = 5 mM and [*tRNA*] = 10*µ*M and [*E*] = 0.25*µ*M show an initial burst of cys-tRNA charging followed by a slower steady state recapitulating what was observed experimentally in Ref. [9]. (C) Single turnover simulation of the transfer rate (*k*_*tran*_) and (D) of the overall chemistry step (*k*_*chem*_) show excellent agreement with experimentally observed values in Ref. [9].

As with cysRS, Table 2 gives the *in vitro* measurements of *k*_*cat*_ and *K*_*m*_ values for *E. coli* hisRS. In this case, my fitting procedure (see Methods and Supplementary S1 Text) has identified kinetic parameters for the kinetic model which, for pyrophosphate exchange, give values of 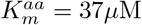, 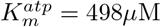, and *k*_*cat*_ = 130 s^*−*1^. Similarly for aminoacylation, the kinetic model gives values of 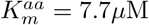, 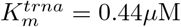, and *k*_*cat*_ = 7.2 s^*−*1^. As with cysRS, one can see that these values are within the range of those that have been measured experimentally for hisRS (Figure 4A). In addition, the identified parameters and kinetic model for hisRS results in no burst of charged his-tRNA^*his*^ (Figure 4B) as well as the observed *k*_*tran*_ and *k*_*chem*_ values in simulations of single turnover kinetics (Figures 4C and 4D).

**Table 2.**
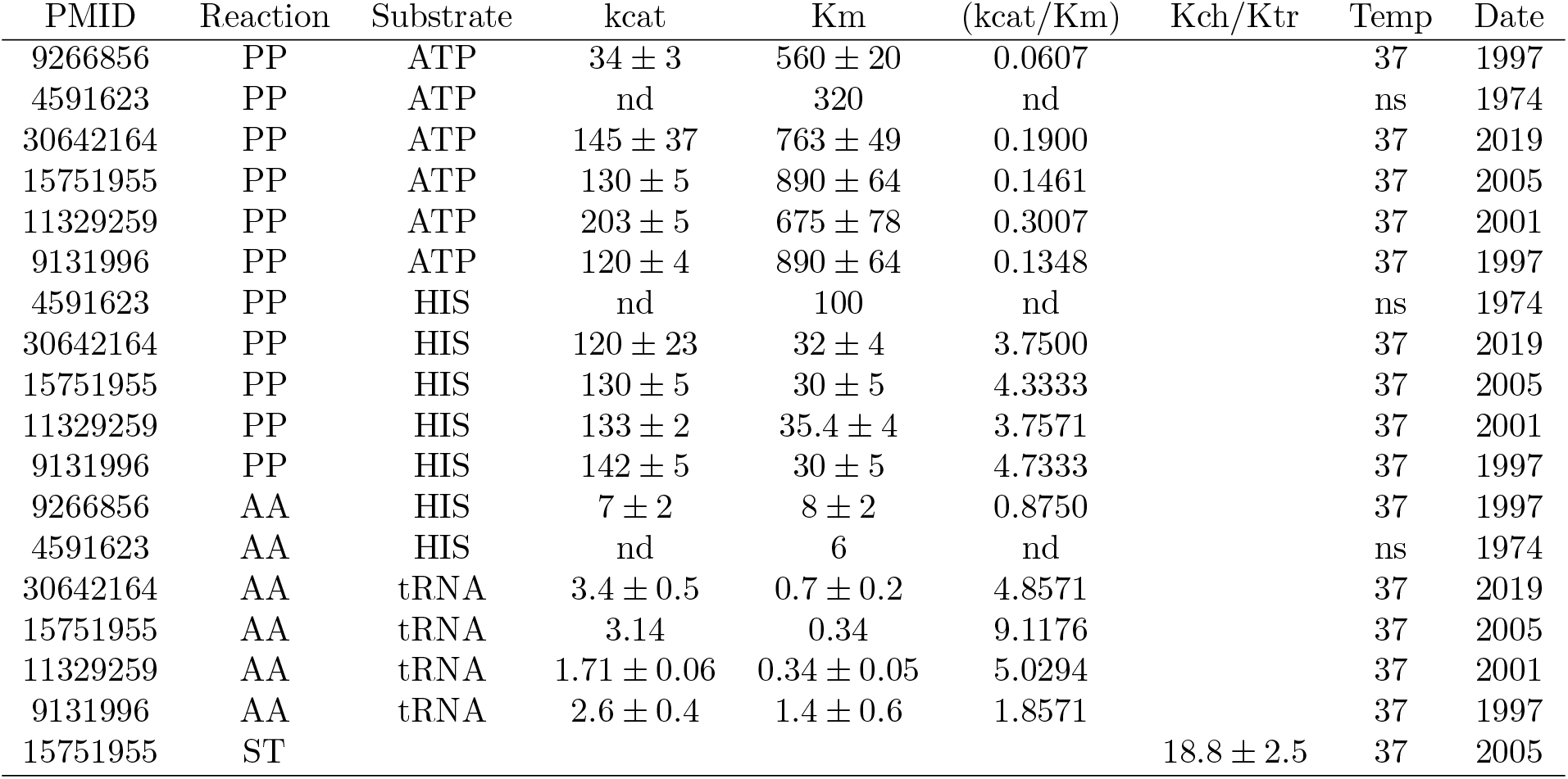
In vitro measurements of *E. coli* hisRS *K*_*m*_ and *k*_*cat*_ values. The experimental measurements for three separate kinetic assays are reported for the Class II hisRS aminoacyl-tRNA synthetase enzyme. Abbreviations are, PP = pyrophosphate exchange, AA = aminoacylation, ST = single turnover transfer, and SC = single turnover overall chemistry. Units for *k*_*cat*_ are in *s*^*−*1^ and *K*_*m*_ are in *µ*M. nd/ns = not determined/stated.

**Fig 4.**
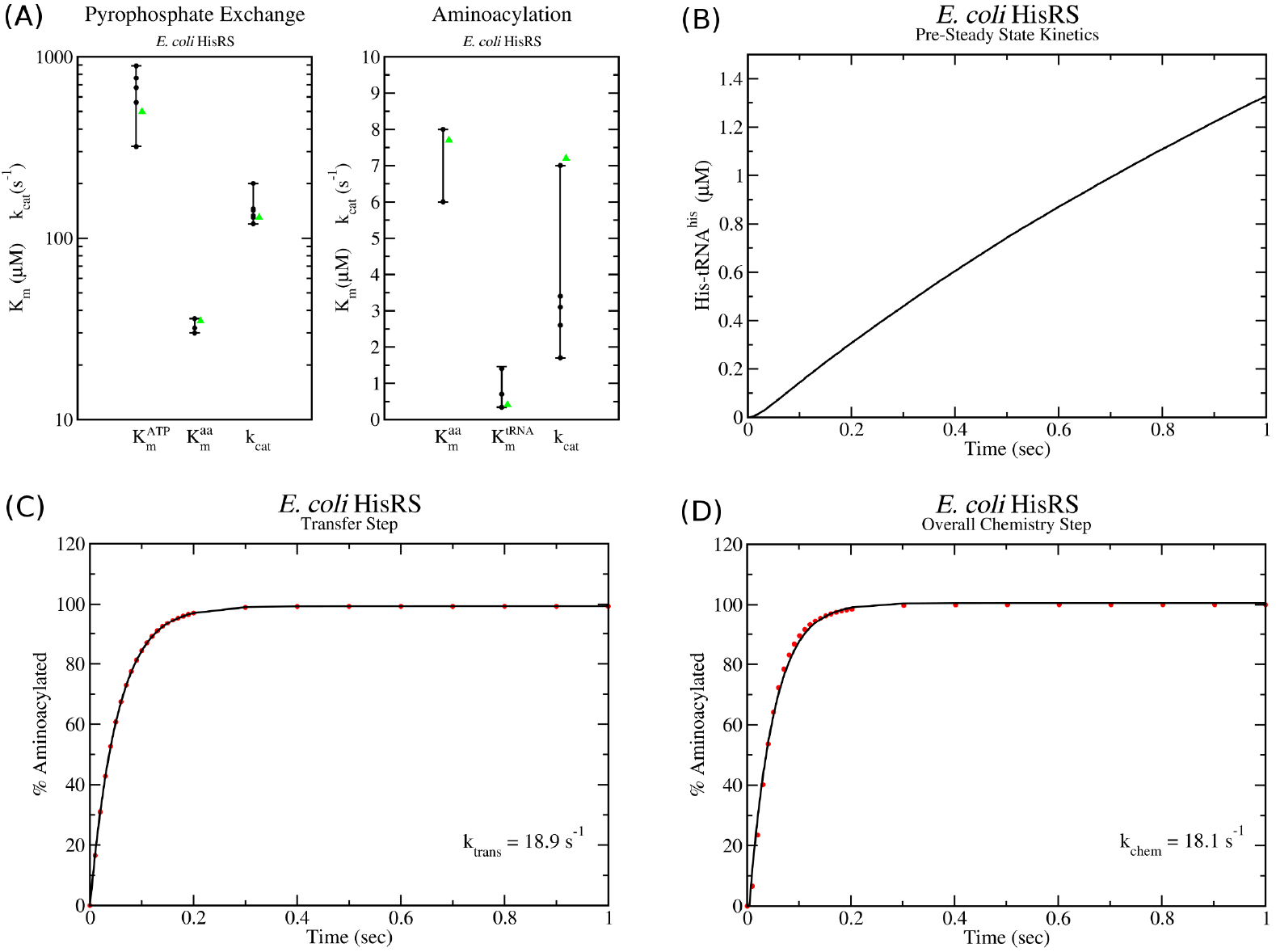
Kinetic model of class II hisRS tRNA charging. (A) Range of *k*_*cat*_ and *K*_*m*_ values determined by *in vitro* pyrophosphate and aminoacylation assays (black dots) are compared with the values of the kinetic model (green triangles). (B) Pre-steady state kinetics simulation with starting concentrations of [*his*] = 500*µ*M, [*ATP* ] = 5 mM and [*tRNA*] = 10*µ*M and [*E*] = 0.25*µ*M show no burst of his-tRNA charging as observed experimentally in [17]. (C) Single turnover simulation of the transfer rate (*k*_*tran*_) and (D) of the overall chemistry step (*k*_*chem*_) show excellent agreement with experimentally observed values in Ref. [17].

As with cysRS and hisRS, I have also parametrised the remaining 18 tRNA synthetases from *E. coli* by identifying a set of kinetic parameters that reproduce experimental measurements for *k*_*cat*_ and *K*_*m*_. Kinetic rates for each of the enzymes can be found in the supplementary information (S1 Text, Tables B, C, and D), while information on the experimental *k*_*cat*_ and *K*_*m*_ measurements and the models fit to them can be found in Supplementary S2 spreadsheet.

### Optimization and validation of kinetic AARS models

#### Comparison of *in vivo* AARS activity with *in vitro k*_*cat*_ measurements

An important and critical question now arises regarding the *k*_*cat*_ and *K*_*m*_ values that have been measured experimentally *in vitro* for the aminoacyl tRNA synthetases. Specifically, are these *in vitro* measured *k*_*cat*_ and *K*_*m*_ values sufficient to support the tRNA charging rates observed *in vivo* in a typical exponentially growing *E. coli* cell. This question is complicated to answer for a number of reasons. First, bacterial cells undergoing exponential growth will have a cell cycle in which the number of proteins and ribosomes in the cell is increasing up until the point of cell division, after which the cellular contents are partitioned between the two daughter cells. Second, given a sample of cells growing at an average rate *µ* h^*−*1^ in a medium, the distribution of cells in the medium will be at different stages of the cell cycle, hence the cells will have different volumes as well as different numbers of ribosomes and proteins present. Finally, although the *average* growth rate of cells in the medium is *µ*, it is not clear that *all* cells in the medium will have this growth rate. Estimation of the average tRNA turnover by an AARS enzyme per cell is directly dependent on the growth rate (which determines tRNA/amino acid usage) and the number of AARS enzymes in the cell. Use of single cell proteomics measurements [29], which have been noted to contain a high amount of variation in protein numbers between cells, may present difficulties in determining the relation between the number of AARS proteins and the cellular growth rate.

Works by Bremer and Dennis as well as other colleagues in the field have shown that the macro-molecular composition of an *E. coli* cell growing exponentially in culture can be straightforwardly described by simple mathematical relations for the average cell in the medium [27]. Here I take a similar point of view and use, for instance, the amino acid usage rate that would occur *on average* in a exponentially growing culture of *E. coli* cells at an average growth rate of *µ* h^*−*1^. Similarly, I calculate the average number of AARS enzymes that would be present and thus obtain the average tRNA turnover per enzyme. In order to determine the *in vivo* AARS activity from this point of view, one would require estimates of; (1) the average usage rate per second of each tRNA isoacceptor or alternatively the total rate of incorporation of each amino acid (*a*_*i*_, *i* ∈ [1, 20]) into protein by ribosomes in the cell, (2) the average number of aminoacyl tRNA synthetases (*n*_*i*_) in the cell, (3) the average number of tRNAs in the cell, and (4) the average volume of the cell. The amino acid usage and number of each AARS in the cell can be used to estimate the enzyme activity (*r*_*i*_), *i*.*e*. the rate of tRNA turnover, for a single enzyme *in vivo*,

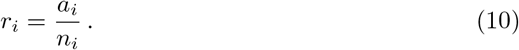

The volume and number of tRNAs on the other hand can be used to determine if the resulting concentration of free uncharged tRNAs is sufficient to achieve a velocity (*v*_*i*_) from the enzyme that is greater than or equal to the tRNA charging rate, *v*_*i*_ ≥ *r*_*i*_.

To estimate the number of each tRNA synthetase *n*_*i*_, I have used 12 measurements of recent proteomics data across five different experimental groups [26, 30–33]. These data points measure average protein abundances in exponentially growing *E. coli* across a variety of growth rates (*µ* = 0.41 to *µ* = 1.98 h^*−*1^) where the elongation rates of ribosomes are estimated to vary between 12-22 aa/sec [27, 34]. It is important to note that the amino acid and tRNA isoacceptor usages are directly related to ribosome elongation rates, which are in turn dependent on the relative concentration of tRNA isoacceptors in ternary complex. Moreover, it has been observed that slower growing bacteria can undergo an immediate increase in protein synthesis following nutrient a up-shift [35] (see discussion in S1 Text, sections 3 and 4). Thus, I assume that the AARS numbers present in the cell are able to support the maximal ribosome elongation rate possible following a nutrient up-shift.

Figure 5 shows the 12 proteomics data points from [26, 30–33] for cysRS and hisRS turnover numbers, which shows a striking amount of variation between experimental groups. The corresponding violin plots have been computed using Eq. 8, and the peak can thus be thought of as the most likely value to be observed in an experiment based on the data points and their errors. This analysis provides the ability to pin-point a ‘consensus” value for the turnover rate as the peak in the violin plot. The red line corresponds to the turnover value that has been used to calculate the number of enzymes via Eq. 10 and for the subsequent optimization of *k*_*cat*_ values in the next section. Similar plots for the remaining 18 AARS enzymes can be found in Supplementary Figures F and G.

**Fig 5.**
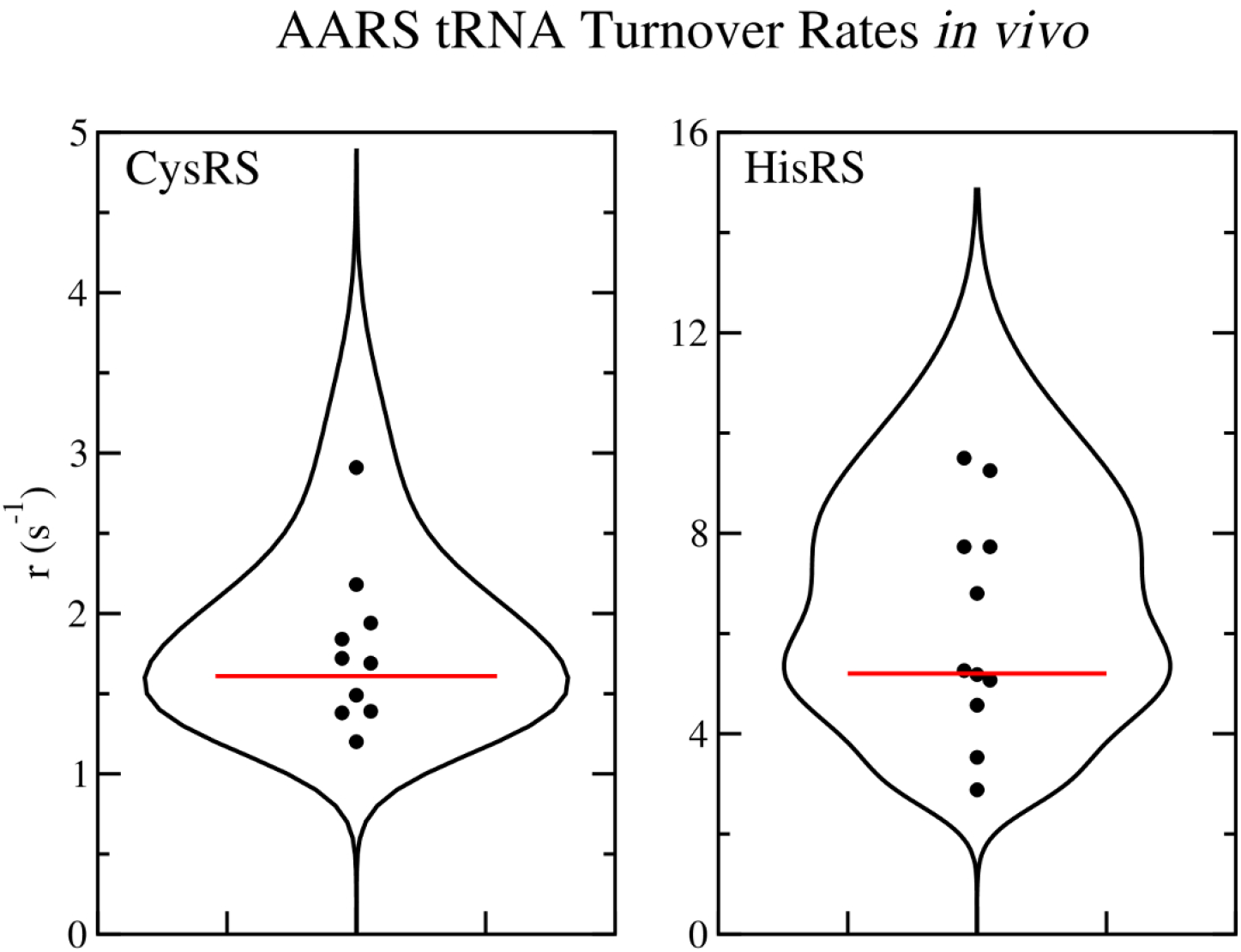
Average tRNA turnover rates *in vivo* for CysRS and HisRS. (A) Violin plot of the average tRNA turnover rates for (A) cysRS and (B) hisRS computed using the 12 proteomics measurements. Red line corresponds to the consensus turnover rate used for computing average enzyme numbers for use in translation simulations.

Finally, Figure 6 shows a comparison of the *in vivo* activity of AARS enzymes (*i*.*e*. their calculated turnover rates) with experimental *in vitro* measurements of the their *k*_*cat*_ values. Black bars indicate the range of *in vivo* activity based on the 12 proteomics data measurements while red bars indicate the range of *k*_*cat*_ measurements. In general, there is some level of overlap between the range of *in vivo* turnover rates and *in vitro k*_*cat*_ measurements for 14 of the enzymes, while 6 (specifically valRS, ileRS, serRS, proRS, thrRS, and alaRS) have measured *k*_*cat*_ ranges below their expected *in vivo* turnover rates based on the proteomics data. This suggests that some of the experimental *k*_*cat*_ measurements will require a shift upwards in order for the enzyme to support the observed *in vivo* turnover rates.

**Fig 6.**
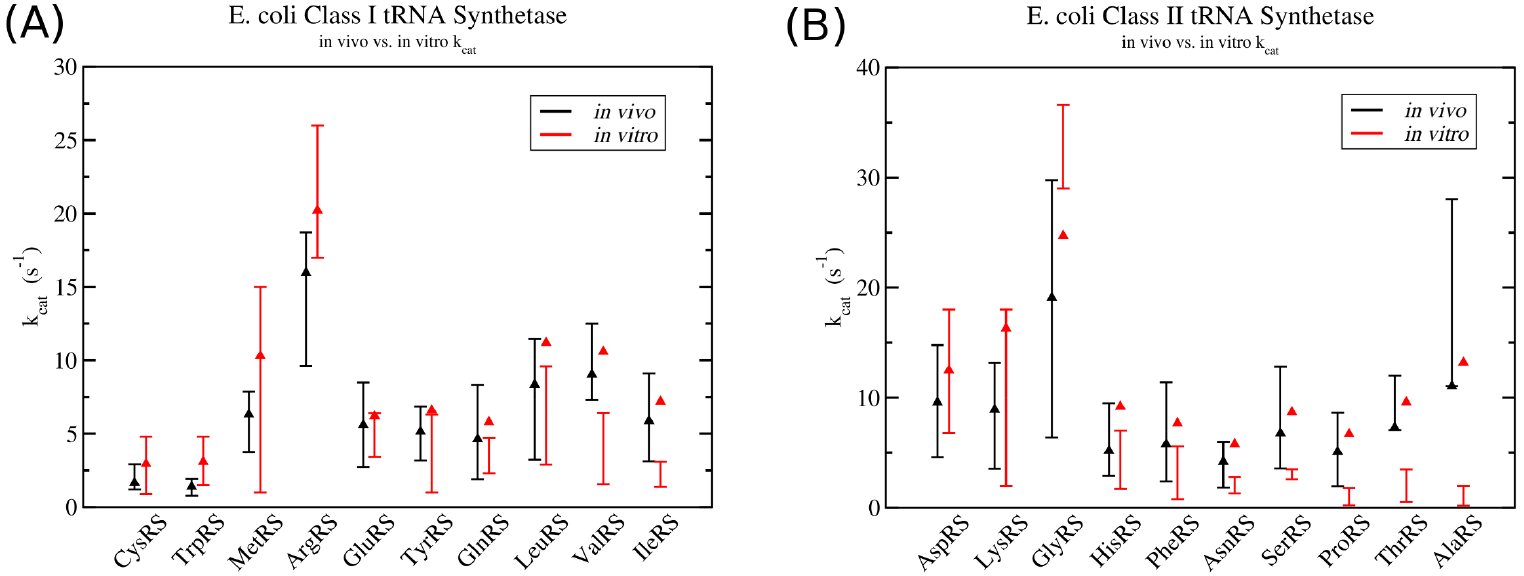
Comparison between *in vivo* activity verses *in vitro k*_*cat*_ measurements in class I and II aminoacyl tRNA synthetases. Black lines with error bars indicate the range of *in vivo* activity for each AARS enzyme determined from proteomics data, while black triangles give the turnover rate used to calculate the number or enzymes for computational simulations. Red lines with error bars correspond to the range of experimental *in vitro k*_*cat*_ measurements. Red triangles represent the optimized *k*_*cat*_ values which are able to support maximum ribosome elongation rate that would be possible following a nutrient up-shift. Data for (A) class I tRNA synthetases and (B) class II tRNA synthetases.

#### Optimized *k*_*cat*_, *K*_*m*_, and tRNA numbers are in reasonable agreement with *in vitro* measurements

We now come to the general problem of reconciling differences between *in vitro* measured aminoacylation *k*_*cat*_ values and the observed turnover rates of the enzymes *in vivo*. We must have the situation that the *k*_*cat*_ values for the AARS enzymes are strictly greater than the *in vivo* turnover rates (*k*_*cat*_ *> r*). Based on the data in Figure 6, it is clear that several of the AARS enzymes will require an adjustment to their *k*_*cat*_ in order for the enzyme to support the observed *in vivo* turnover rate. To make the adjustments, I follow the procedure outlined in Supplementary S1 Text, Section 7. Essentially, this procedure attempts to minimize the discrepancy between computational simulations of *in vivo* translation and tRNA aminoacylation and several experimental observations. These observations are; the average numbers of each tRNA in the cell, proteomics data on the average number of AARS enzymes, and the percentages of charged tRNA in the cell. For full discussion of the optimization procedure, see Supplementary S1 Text, Section 7.

The optimized *k*_*cat*_ values for all 20 AARS enzymes are shown as red triangles in Figure 6, with the optimised turnover rates shown as black triangles. Optimized turnover rates can be converted to enzyme numbers using Eq. 10. Figure 6, along with Figures F and G in S1 Text, reveal that all of the optimized turnover numbers are within the range of experimental proteomics measurements. Moreover, the optimized *k*_*cat*_ values in Figure 6 show that 8 of the AARS enzymes (specifically cysRS, trpRS, metRS, argRS, gluRS, tyrRS, aspRS, and lysRS) have optimized *k*_*cat*_ values within the range of experimental *in vitro* measurements. The remaining 12 AARS enzymes have optimized *k*_*cat*_ values which are above the range of *in vitro* measurements, deviating on average by a factor of 2.08 from the highest measured value. Specific deviation factors are; (valRS - 1.65), (ileRS - 2.32), (leuRS - 1.16), (glnRS - 1.23), (serRS - 2.48), (thrRS - 2.74), (proRS - 3.72), (hisRS - 1.31), (asnRS - 2.07), (alaRS - 6.6), and (pheRS - 1.37). Interestingly, the class I tRNA synthetases have the smallest deviations, while the class II synthetases have the largest, with alaRS and proRS deviating by more than a factor of 3.

The optimized *K*_*m*_ values for both amino acid and tRNA as substrate are shown in Figure 7. Experimental *K*_*m*_ measurements are depicted as violin plots (black lines), with the peak in the violin plot being the most likely value to be observed in an experiment based on the data points and their errors. All of the optimized *K*_*m*_ values, shown as green triangles, fit within the ranges of the experimental *in vitro* measurements. Supplementary Table I summarises the kinetic parameters for each of the enzymes.

**Fig 7.**
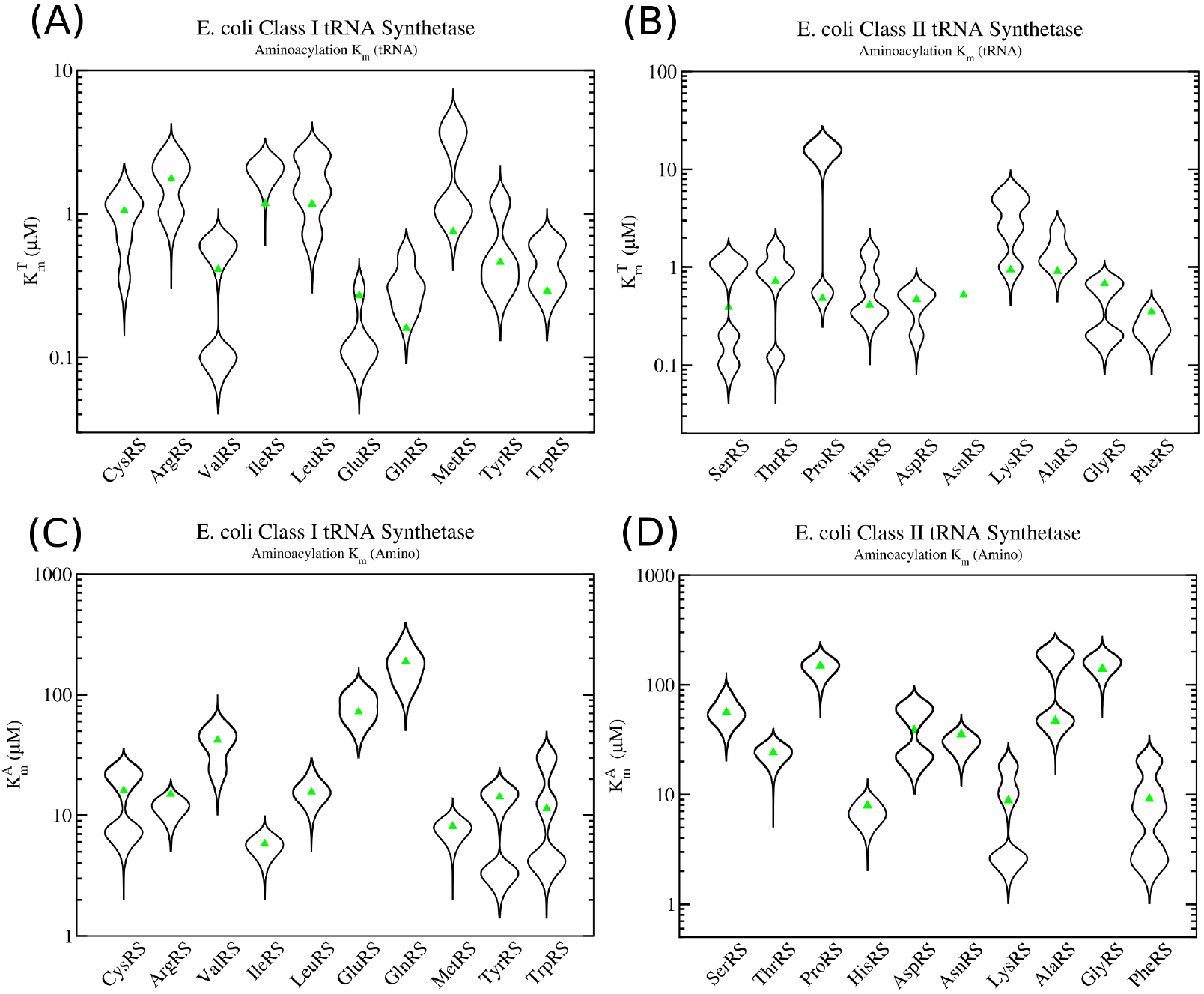
Experimental *in vitro K*_*m*_ measurements in class I and class II aminoacyl-tRNA synthetases. Violin plots of the experimental *in vitro K*_*m*_ measurements are given as black lines. Green triangles represent the optimized *K*_*m*_ values used in translation simulations. (A) 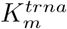 values for tRNA as substrate in the aminoacylation reaction for class I enzymes and (B) class II enzymes. (C) 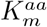 values for amino acid as substrate in the aminoacylation reaction for class I enzymes and (D) class II enzymes.

Finally, Supplementary Table G compares the total number of each tRNA isoacceptor used in the model with experimental measurements from Dong *et al*. [28] for *E. coli* cells growing at *µ* = 0.41 h^*−*1^ (doubling time of *τ* = 100 min). The average absolute error from the values in Dong *et al*. at this growth rate is 0.3, with the majority of the error coming from only a few tRNAs, specifically metM, his, gln1, gln2, arg3, lys, asn, pro3, and phe. However, when examining the model fit to different growth rates (see Supplementary Spreadsheet S2, Tab 8), one sees that there is sometimes better agreement between the model and the measurements. Taking the minimal absolute error for each tRNA over all growth rates one finds that the error is only 0.18 on average, with the largest errors coming from tRNAs lys, asn, pro3, and phe. Supplementary Table J gives the optimal tRNA numbers for growth rates of *µ* = 0.41, 0.69, 0.98 and 1.73 which are used in the full model.

#### Simulations of *in vivo* tRNA charging kinetics

With optimized *k*_*cat*_ and *K*_*m*_ values for the AARS enzymes identified, it now remains to validate that the tRNA charging kinetics of the enzymes can support the translation and tRNA turnover rates observed *in vivo*. I have Incorporated the kinetic reaction scheme for class I and II enzymes (Figure 1) into my stochastic model of stranslation [24, 25] using the kinetic parameters in Supplementary Tables B, C, and D. The values of *k*_*cat*_ and *K*_*m*_ that result from these parameters are shown in Supplementary Table I. The stochastic model of *in vivo* translational kinetics of *E. coli* in Refs. [24, 25] is capable of simulating up to 60k ribosomes on 10k mRNAs and takes into account all known biochemical steps in the translation process. A few examples of features included in the model are; Ef-Tu recharging by Ef-Ts, competition between tRNAs for the A-site, stalling between ribosomes on the same mRNA, and initiation, recycling, and termination events on the mRNAs.

Figure 8 shows the charging kinetics of tRNA^*cys*^ and tRNA^*his*^ from a simulation of translation *in vivo* at a growth rate of *µ* = 0.69 h^*−*1^ with a doubling time of *τ* = 60 min. At this growth rate, there are roughly 15000 ribosomes and the total nucleotide content in the cell from the mRNA transcriptome is roughly 2 × 10^6^ nucleotides [27]. To insure that the AARS kinetics can support a nutrient up-shift, I simulate translation post nutrient up-shift by using experimental estimates for amino acid concentrations that would be found for a minimal media supplemented with amino acids (see Table H). The simulation of translational dynamics takes part in two stages. The first stage is a constrained initiation of the system, where tRNAs are allowed to be rapidly charged, followed by a slow equilibration of the AARS enzyme dynamics with the translational dynamics (pre-steady state). This constrained initiation is biologically artificial and is to ensure that the charging demand of the 20 AARS enzymes do not exceed their maximal capacity during initiation of the simulation. After 40 seconds, the simulation is allowed to enter unconstrained dynamics (steady state). As can be seen in Figure 8, the dynamics of cysRS and hisRS with optimized *k*_*cat*_ values of 2.9 and 9.2 s^*−*1^, respectively, are sufficient to support ribosome translation at the maximal average elongation rate of 18.6 aa/sec, which occurs at this growth rate post nutrient up-shift. The remaining 18 AARS enzymes, with their *k*_*cat*_ values given in Table I, have also been verified to support translational dynamics at these elongation speeds. Finally, all 20 AARS enzymes have been found to support a maximal average elongation speed of 19.1 aa/sec at the highest growth rate used in this study of *µ* = 1.73 h^*−*1^, and are stable for over 30 min of simulation time.

**Fig 8.**
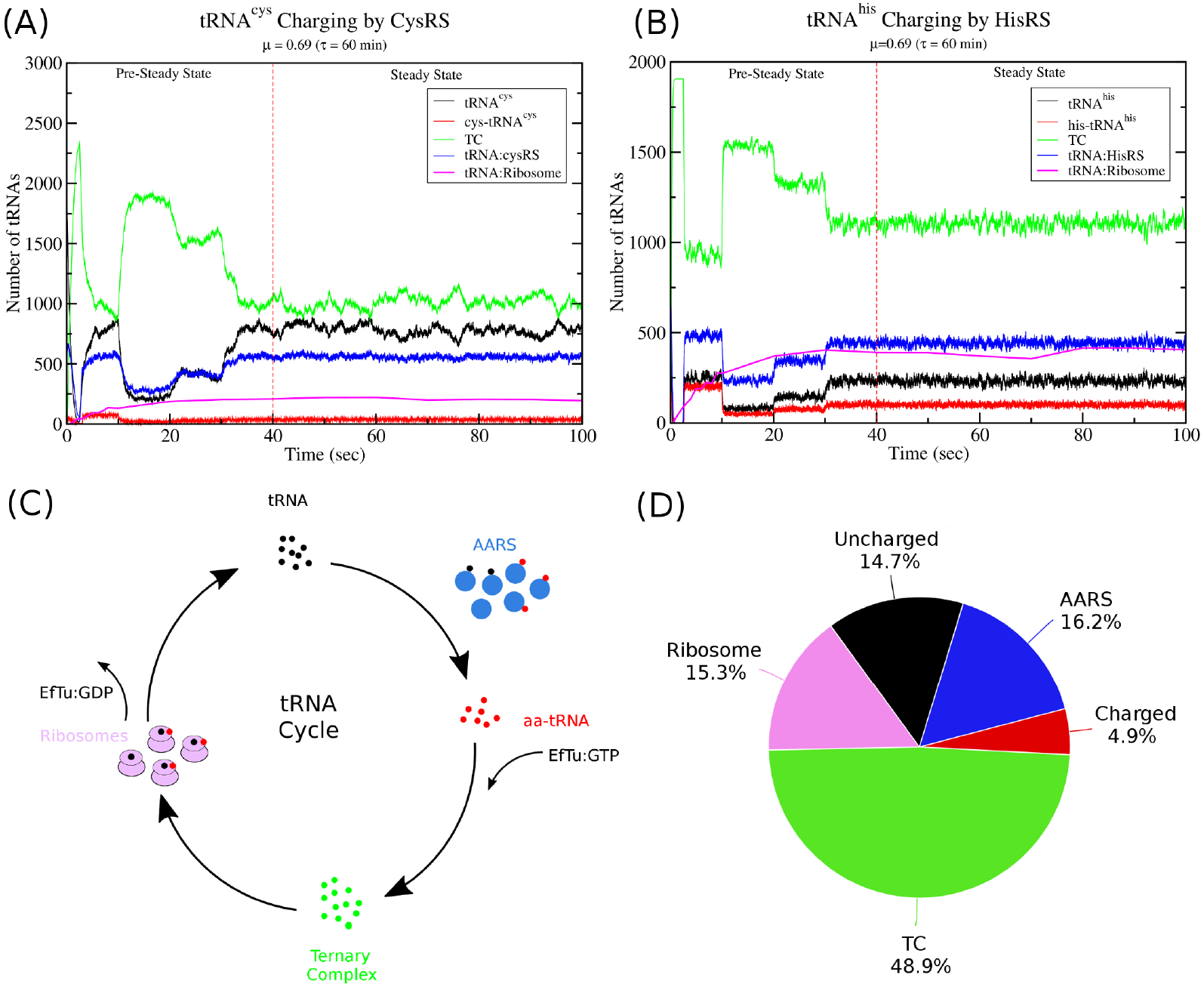
Computational Simulations of tRNA Charging and Translation *in vivo*. (A) The kinetics of tRNA^*cys*^ charging by cysRS during translation. (B) The kinetics of tRNA^*his*^ charging by hisRS during translation. (C) The movement of tRNA during the translation cycle. Uncharged tRNAs (black) are charged by AARS enzymes (blue) and released as aa-tRNA (red). The aa-tRNA then binds to EfTu:GTP to form ternary complex (green) which is recruited to the ribosome. Ribosomes (purple) incorporate the amino acid into the peptide chain releasing EfTu:GDP and uncharged tRNAs. (D) Partitioning of the *total* amount of tRNA in the cell. Percentage of total tRNA as free uncharged tRNAs (black), free charged tRNAs (red), free tRNAs in ternary complex (green), and tRNA bound to either ribosomes (purple) or AARS enzymes (blue) is shown as a pie chart at a growth rate of *µ* = 0.69 (*τ* = 60 min).

Interestingly, as illustrated in Figure 8C and 8D, a substantial portion of the tRNA remains bound to AARS enzymes in the steady state. When averaged over all the tRNAs, roughly 16.2% of the total tRNA in the cell is bound to AARS enzymes either as charged or uncharged tRNAs. This, as we will see in the next section, has important implications for the use of the Michaelis–Menten equation when modelling AARS kinetics.

#### Effect of intracellular amino acid concentration on translational speed and tRNA charging

It should be noted that I have parametrised the AARS enzyme *k*_*cat*_ and *K*_*m*_ values to ensure that they can support the fastest translational speeds that would occur after a nutrient up-shift in exponentially growing *E. coli* cells. For instance, I calculate the average translational speed of ribosomes in cells with a doubling time of *τ* = 100 min post nutrient up-shift is roughly 18.0 aa/sec for all mRNAs and 16.2 aa/sec for the LacZ mRNA. These values are quite similar to the average translational speeds of 19.1 and 17.3 aa/sec that I calculate over all mRNAs or the LacZ mRNA at one of the fastest growth rates (doubling time of 24 min), where nutrients are presumed to be plentiful. However, in minimal media with no supplemented amino acids, the average translational speed of the ribosome over all mRNAs have been experimentally estimated to be roughly 15 aa/sec by Bremer and Dennis at a doubling time of 100 min [27]. Dai *et al*. on the other hand measure the average translational speed of ribosomes on the LacZ mRNA and find values of roughly 12 aa/sec at a cell doubling time of 100 min [34].

To test if my model is capable of recapitulating these lower average ribosome speeds in nutrient poor conditions, I have calculated the average translational speed of ribosomes at the growth rate of *µ* = 0.41 h − 1 in minimal media *without* supplemented amino acids. Since *E. Coli* is capable of synthesising all 20 amino acids, the lack of supplied amino acids must presumably result in lower intracellular concentrations of at least some of the 20 amino acids. Experimental measurements of intracellular amino acid concentrations by Bennett *et al*. [20], as well as Avcilar-Kucukgoze *et al*. [36], suggest that the amino acids tryptophan, tyrosine, phenylalanine, and serine may be close to the 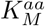 values for their enzymes in minimal media conditions. Using the intracellular amino acid concentrations shown in Supplementary Table H, I simulate translation at the growth rate of *µ* = 0.41 h − 1 and find that the average translational speed of ribosomes are 13.3 aa/sec averaged over all mRNAs and 12.5 aa/sec averaged over only the LacZ mRNA. Both of these theoretical calculations are in good agreement with measurements from Dai *et al*. [34].

Finally, I compare the fraction of charged tRNAs predicted by the model and compare with the observations from several experiments [36–39]. For tRNAs with more then one isoacceptor, the averages for each are combined and weighted by the number of each tRNA isoacceptor in the model. As can be seen in Figure 9, there is very good agreement with the experimental observations, with only tRNA^*ile*^, tRNA^*his*^, tRNA^*asp*^, and tRNA^*asn*^ being outside the range of experimental measurements. Moreover, the results show that the majority of tRNAs are charged between 70 and 90 %, even at this lower growth rate. A comparison with the results of Avcilar-Kucukgoze *et al*. [36] by individual isoacceptor is shown in Supplementary Figure I.

**Fig 9.**
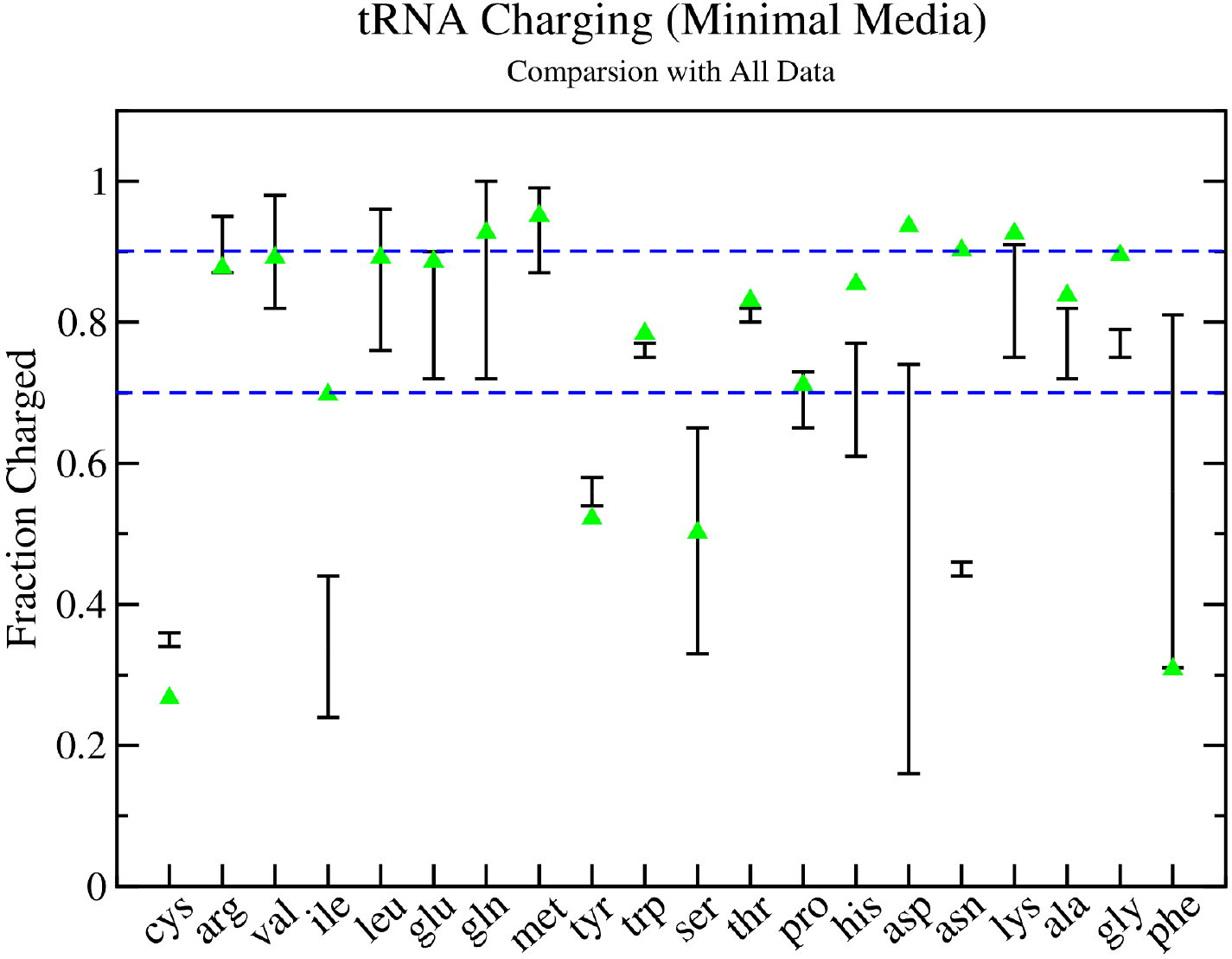
Theoretical and Experimental Estimates of tRNA Charging Fractions in Minimal Media. The total fraction of tRNAs charged in exponentially growing *E. coli* cells. Black lines with error bars indicate the range of experimental measurement from several different experiments [36–39], while green triangles give the results of the stochastic translational model with aminoacyl tRNA synthetase kinetics following the reaction scheme in Figure 1 at a growth rate of *µ* = 0.41 h^*−*1^.

#### Development of a Michaelis–Menten Model of AARS Kinetics

Simulating the full kinetic model of AARS kinetics, as depicted in Figure 1, will usually have substantial computational costs. Because of this, it is often useful to simplify the kinetic model in order to accelerate the computation speed. One of the most popular and highly used models for enzyme kinetics is the Michaelis–Menten model, which allows for the rate product formation to be estimated from the equation

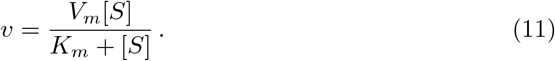

The rate of product formation *v* depends on the concentration of free substrate [*S*] and the maximal velocity (or maximal rate of product formation) of the enzyme *V*_*m*_. The quantity *V*_*m*_ in Eq. 11 is related to the total amount of enzyme [*E*_0_] and the enzymes *k*_*cat*_ via *V*_*m*_ = *k*_*cat*_[*E*_0_]. The *K*_*m*_ is the Michaelis–Menten constant which can be thought of in simple terms as the concentration of substrate at which the enzyme works at half of *V*_*m*_. Typically one models the formation of product and loss of substrate using the ODEs

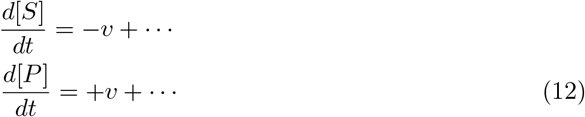

where the dots denote that there may be additional terms in the ODEs which, for example, model binding of substrate and product to other proteins.

The Michaelis–Menten (MM) model is one of the most popular models for enzyme kinetics due to its simplicity and has been previously used to examine AARS kinetics in a model of *in vivo E. coli* metabolism [10]. However, there are at least two issues when using the MM equation (Eq. 11) to model AARS enzyme kinetics. First, there are several substrates (ATP, amino acid, and tRNA) and each of these will impact on the speed of product formation. One potential fix is to say that ATP is saturating, and thus model the velocity using the modified equation [10]

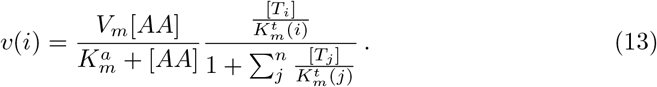

This equation accounts for charging of *n* different tRNA isoacceptors by a single AARS enzyme. For instance, glnRS has two tRNA isoacceptors that it charges (tRNA^*gln*1^ and tRNA^*gln*2^), and *v*(*i*) is the rate of charging for tRNA isoacceptor *i* with free concentration [*T*_*i*_]. The second, and more critical issue, is that there are several assumptions about the enzyme and substrate that must be true in order for the MM equation to be accurate. The MM equation is typically derived assuming that; (1) the total amount of enzyme is much less than the free concentration of substrate and, (2) the enzyme is present as either free enzyme or a complex of enzyme and substrate. The first condition is clearly violated here, as proteomics data [26, 30–33] and measurements of total tRNA concentrations [28] show that the number of AARS enzymes is often similar to the total tRNA numbers. For example, Refs. [26, 31] both report around 800 cysRS enzymes at a growth rate of *µ* = 0.69 h^*−*1^, while Dong *et al*. measure around 3200 tRNA^*cys*^. Most of this tRNA will be in ternary complex or charged, and as can be seen from the simulation of the full cysRS kinetic model in the presence of translation in Figure 8A, around 750 tRNA^*cys*^ are uncharged, similar to the number of cysRS enzymes. Similarly, the second condition is also violated as a substantial amount of aa-tRNA (the product) remains bound to the enzyme during steady state catalysis (cf. Figure 8).

In one sense these are minor issues as I have found that, in general, the AARS enzyme kinetics modelled by the reaction scheme in Figure 1 clearly can be approximated by the MM equation. However potential problems arise when optimising *k*_*cat*_ and 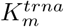 values for a MM model. For example, if the MM model does not correctly account for the amount of tRNA that is bound to the enzyme, then this will result in an over estimation of the quantity of uncharged tRNAs, or tRNAs in ternary complex, *etc*. as one tries to reconcile the model with the total tRNA measurements from Dong *et al*. [28]. For example, 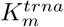 values may need to be increased in order to compensate for the extra uncharged tRNA. There are two approaches that can be used to fit *k*_*cat*_, *K*_*m*_, and overall tRNA numbers for an MM model; (1) use the tRNA numbers from Dong *et al*. and adjust *k*_*cat*_ and *K*_*m*_ appropriately, or (2) use the *K*_*cat*_ and *K*_*m*_ values that have been optimised for the full kinetic reaction scheme in Figure 1 (see Supplementary Table I for values) and adjust the tRNA numbers. Since the MM model does not properly account for the amount of charged tRNA product that remains bound to the enzyme, I have taken the latter approach here and adjusted the tRNA numbers to account for this issue. The optimization procedure can be found in Supplementary S1 Text, Section 7.

The final optimised tRNA numbers for various *E. coli* growth rates are given in Supplementary Table J. This table gives both the optimised tRNA numbers for the MM model as well as for the full kinetic reaction scheme shown in Figure 1. As can be seen, there is a reduction in the number of tRNAs for the MM model compared to the full kinetic model. This illustrates that the full kinetics of AARS enzymes, as well as the status and location of the tRNAs (*i*.*e*. are they bound to AARS, bound to ribosome, or in free ternary complex), need to be accounted for in order to correctly compare with the experimental tRNA numbers from Dong *et al*. [28]. Details of the amount of tRNA that is bound to ribosome and is present in ternary complex used for the fitting of the full model can be found in Supplementary Spreadsheet S2, Tab 8.

## Discussion

I have developed an empirical kinetic model for all 20 tRNA synthetase enzymes from *E. coli*, where the individual binding and catalytic events that take place in the aminoacylation process are explicitly considered. Model predictions for each of the 20 kinetic models have been validated and shown to support steady state translational kinetics in an average cell undergoing exponential growth with doubling times of *τ* = 100, 60, 40, and 24 min. The resulting optimised *k*_*cat*_ values are in reasonable agreement with *in vitro k*_*cat*_ measurements, with 8 out of 20 AARS enzymes having optimized values within the ranged observed in experiments. The remaining 12 AARS have optimised *k*_*cat*_ which deviate from experiment by only a factor of 2 on average. Likewise, the optimized Michaelis–Menten constants for amino acid and tRNA (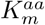 and 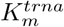) are also within experimental ranges for all 20 AARS enzymes (cf. Figure 7). Despite these results, it is still important to note that the kinetic parameters and overall reaction scheme for the aminoacylation model depicted in Figure 1 should not be considered as definitive, but instead should be thought of as a starting point for refinements as further experimental data are obtained in future. However, several important conclusions can be drawn based on these empirical kinetic models.

One of the important conclusions from the analysis of this work is that the total tRNA in the cell is partitioned as free uncharged/charged tRNA as well as bound tRNA, and that a substantial fraction (approximately 15% of the total tRNA) remains bound to AARS enzymes during steady state translation. Dong *et al*. [28] have previously noted that while the amount of tRNA in ternary complex required for optimal elongation rate should be proportional to the square root of the codon frequency (see Section 4 in S1 Text), the total tRNA abundances measured roughly scale with the square of the codon frequency. By taking into account the number of tRNAs bound to the ribosome, Dong *et al*. showed that the resulting total tRNA abundances more closely fit to the square of the codon frequency. However, they assumed that tRNA was only either in free ternary complex or bound to ribosomes.

Based on the kinetic model in Figure 1, I have shown that the fraction of charged and uncharged tRNA that is bound to AARS enzymes will likely also represent a substantial portion of the total tRNA. This has important consequences for the fitting of parameters for Michaelis–Menten models of AARS kinetics and their subsequent validation with experimental observations as discussed in the results.

A second conclusion is that most of the *in vitro* experimental measurements of *k*_*cat*_ and *K*_*m*_ values are fairly close to what are needed to support the estimated tRNA aminoacylation rates in the cell. In this work, I have found that 8 out of the 20 AARS enzymes (cysRS, trpRS, metRS, argRS, gluRS, tyrRS, aspRS, and lysRS) have *in vitro* measured *k*_*cat*_ values which are within the range of expected turnover rates determined by proteomics measurements (cf. Figure 6). The remaining 12 AARS enzymes have optimised *k*_*cat*_ values which deviate on average by a factor of 2 from experimental *in vitro* measurements. This is in contrast to recent work by Choi and Convert [10] which parameterised a Michaelis–Menten model of AARS kinetics for all 20 AARS enzymes and found that their optimised values deviated from *in vitro k*_*cat*_ measurements on average by roughly a factor of 7 [10]. There are several reasons for the discrepancy with my work here.

First, Choi and Covert do not seem to distinguish between the *k*_*cat*_ for aminoacylation versus that for pyrophosphate exchange. For example, the *k*_*cat*_ for pyrophosphate exchange for cysRS (see Table 1) tends to be measured around 90 s^*−*1^, about 30 times faster than the *k*_*cat*_ measurements for aminoacylation (around 2.9 s^*−*1^). Instead, their fitting procedure appears to consider both values when optimising *k*_*cat*_. It is important to note that the pyrophosphate exchange *k*_*cat*_ is measuring the exchange of ATP and pyrophosphate on an adenylate bound enzyme. Thus, it is a proxy for the rate of the first catalytic event, *i*.*e*. activation of the amino acid. In contrast, the aminoacylation *k*_*cat*_ measures the entire process of amino acid activation plus transfer of the amino acid to the tRNA and is the only one that has a relation to the measured *in vivo* turnover rates. Their higher reported aminoacylation *k*_*cat*_ for cysRS (69.44 s^*−*1^ [10]) is potentially due to their fitting algorithm being allowed to search up to 90 *s*^*−*1^. As a result, they report a factor of ≈ 3 deviation from an experimental average of 23.1 ± 35.9 s^*−*1^, where this average includes the pyrophosphate exchange *k*_*cat*_ measurements. Comparing to only experimental aminoacylation *k*_*cat*_ values in Table 1, the value 69.44 deviates from the average of 2.96 s^*−*1^ by over a factor of 23.

Second, there seems to be a substantial difference between AARS numbers that I have calculated here versus what Choi and Covert have calculated. To calculate AARS numbers, Choi and Covert use their Parca *E. coli* model [40] to estimate protein numbers in the cell. They then compare the average protein numbers with those from Schmidt *et al*. [32] and with single cell proteomics data from Taniguchi *et al*. [29]. Next, they estimate amino acid usage in the cell by calculating the number of proteins to be doubled, and use the codon composition of the protein’s mRNA and doubling time of the cell to determine the codon reading rate for each codon. In contrast, I have used averaged proteomics data from 12 different measurements to estimate consensus AARS numbers and have used my ribosome translational model which simulates translation on the full transcriptome in the cell. My transcriptome was constructed using mRNA-seq measurements from Li *et al*. [26] (see Methods) to estimate the number of each mRNA present in the cell. Remarkably, we are both in very good agreement on the amino acid usage / codon reading rates. Choi and Covert report for *E. coli* with ≈ 26000 ribosomes in minimal media supplemented with amino acids a usage rate for the amino acid cystine of 4.478 *µ*M s^*−*1^ (or 2696.6 s^*−*1^ in the average cell volume of 1 *µ*m^3^ at this growth rate). I find a very similar result using a transcriptome based on mRNA-seq data from Li *et al*. [26] of 4.508 *µ*M s^*−*1^ (or 2714.8 s^*−*1^). Despite the good agreement on amino acid usage rates, we substantially differ on AARS numbers. For example, I have calculated an optimal value of 1610 enzymes for cysRS, similar to the 1409 measured by Mori *et al*. [31] for bacteria with a growth rate of approximately *µ* = 1.04 h^*−*1^. In contrast, Choi and Covert estimate 606 on average at a very similar growth rate [10]. As a result, my average turnover rate for cysRS is 1.68 s^*−*1^ versus 4.44 s^*−*1^ for Choi and Covert, roughly 3 times lower. There is a similar trend over the remaining AARS enzymes (see Supplementary S1 Text, Table K). Choi and Covert report that, since their simulation of average protein numbers from the Parca model are within a factor of 10 from Schmidt *et al*. [32] and had a coefficient of determination of *R*^2^ = 0.63, there was satisfactory agreement with the measured proteome values of Schmidt *et al*. [10]. However, as Figure 5 shows, if we perform a meta analysis by including additional proteomics measurements from several different groups, we see that there is substantial variation amongst the measurements. Although this suggests that relying on a single proteomics data point may be potentially problematic, a consensus value does seem to emerge when weighted over multiple measurements from different groups.

Finally, Choi and Covert report optimised values of *k*_*cat*_ = 69.44 s^*−*1^ and 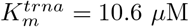 for cysRS [10]. This is in contrast to the optimised values that I find here of *k*_*cat*_ = 2.9 s^*−*1^ and 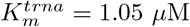 Given Choi and Covert’s estimated turnover rate of *r* = 4.44 s^*−*1^ for cysRS, one can solve for the amount of free uncharged tRNA in their model using the MM equation (Eq. 11) and find [*T*_*u*_] = 0.724 *µ*M. Thus in a quasi-steady state, their translational model with a MM model of aminoacylation should have each cysRS enzyme averaging a turnover of 4.44 tRNAs per second, with a free concentration of uncharged tRNA^*cys*^ of 0.724 *µ*M in the cell. But in the Michaelis–Menten model, *k*_*cat*_ and 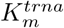 satisfy the relation

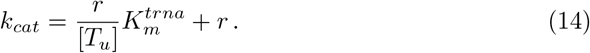

This relation implies that, given the enzyme turnover rate *r* and the amount of free uncharged tRNA [*T*_*u*_], a re-scaling of *k*_*cat*_ and 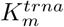 is possible. This re-scaling procedure does not seem to have been applied in their fitting procedure, or it was applied to values that incorrectly included the pyrophosphate exchange *k*_*cat*_ values, which tend to be an order of magnitude higher. For example, applying this re-scaling to Choi and Covert’s cysRS parameters, one can obtain new values of *k*_*cat*_ = 8.36 s^*−*1^ and 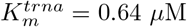, which are much closer to the two experimental *in vitro* measurements of *k*_*cat*_ = 4.8 s^*−*1^ and 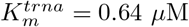 reported in Table 1.

In summary, the models I have developed here provide a basis for examining tRNA charging that occurs *in vivo* by aminoacyl tRNA synthetases in *E. coli* cells. The analysis has revealed that for 8 out of the 20 AARS enzymes, the *in vitro* measurements of *k*_*cat*_ and 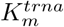 are in line with the enzyme turnover rates that would be expected *in vivo* based on average proteomics measurements, with the remaining 12 requiring a small adjustment by a factor of 2 on average. However, a few class II enzymes, in particular, serRS, thrRS, proRS, and alaRS, deviate more substantially. More experimental data will be needed to elucidate the origin of this discrepancy, and refine the kinetic models appropriately in these cases. Regardless, the models I report on here should hopefully be an important tool for the community to theoretically investigate, for example, the effects of amino acid supply on the cellular translational machinery in bacteria.

## Supporting information

**S1 Text. Supplementary Text**. https://doi.org/10.1371/journal.pcbi.xxxxxx.xxx

**S2 Spreadsheet. Supplementary Spreadsheet**.

https://doi.org/10.1371/journal.pcbi.xxxxxx.xxx

**Fig. A Kinetic reaction scheme for the pyrophosphate exchange reaction**. State *S*_0_ represents the AARS enzyme, while *S*_1_ and *S*_2_ are the amino acid and ATP bound enzymes, respectively. State *S*_3_ is the enzyme with both amino acid and ATP bound, *S*_4_ is the adenylate and pyrophosphate bound state, while state *S*_5_ is the adenylate bound enzyme state.

**Fig. B Effect of parameter variation on pyrophosphate exchange kinetics**. (A-C) Effect on Michaelis–Menten constant *K*_*m*_ and pyrophosphate *k*_*cat*_ as the amino acid and ATP dissociation constant is varied. (D-F) Effect on the Michaelis–Menten constant *K*_*m*_ and pyrophosphate *k*_*cat*_ as the amino acid activation rates *k*_5*f*_ and *k*_5*b*_ are varied. (G-I) Effect on the Michaelis–Menten constant *K*_*m*_ and pyrophosphate *k*_*cat*_ as the pyrophosphate release rates *k*_6*f*_ and *k*_6*b*_ are varied.

**Fig. C Kinetic reaction scheme for Class I aminoacyl tRNA synthetases**. Reaction scheme diagram labelling the individual kinetic reactions in the aminoacylation reaction for the class I aminoacyl tRNA synthetase enzymes. Individual kinetic rates for each reaction can be found in Supplementary Table B.

**Fig. D Kinetic reaction scheme for Class II aminoacyl tRNA synthetases**. Reaction scheme diagram labelling the individual kinetic reactions in the aminoacylation reaction for the class II aminoacyl tRNA synthetase enzymes. Individual kinetic rates for each reaction can be found in Supplementary Tables C and D.

**Fig. E The dependence of the translation elongation rate on total ternary complex concentration**. Stochastic simulations of translation *in vivo* are used to calculate the average elongation rate of ribosomes in *E. coli* for different cellular volumes verses the percentage of the total amount of tRNA in ternary complex. The fractions of each tRNA isoacceptor in ternary complex set to the optimal values listed in Table G (see section 4 for further details).

**Fig. F Estimated average tRNA turnover rates for class I aminoacyl tRNA synthetases**. Violin plots for the AARS tRNA turnover rates are shown for each of the class I enzymes, with cysRS shown in Figure 5 of the main text. Each of the individual proteomics data points used to construct the violin plot are shown (black dots) along with the value used for optimization (red line).

**Fig. G Estimated average tRNA turnover rates for class II aminoacyl tRNA synthetases**. Violin plots for the AARS tRNA turnover rates are shown for each of the class II enzymes, with hisRS shown in Figure 5 of the main text. Each of the individual proteomics data points used to construct the violin plot are shown (black dots) along with the value used for optimization (red line).

**Fig. H Average tRNA usage rates for the three proline tRNA isoacceptors**. Theoretical average tRNA usage rates for tRNA^*pro*1^ (black line), tRNA^*pro*2^ (red line) and tRNA^*pro*3^ (green line) are shown using a fixed ratio of tRNA^*pro*2^ to tRNA^*pro*3^ of y = 1.25 in ternary complex. The graph is plotted as a function of the ratio of tRNA^*pro*1^ to tRNA^*pro*3^ in ternary complex.

**Fig. I Theoretical and Experimental Estimates of tRNA Charging Fractions in Minimal Media**. The total fraction of tRNAs charged in exponentially growing *E. coli* cells. Black lines with error bars indicate the range of experimental measurement from Avcilar-Kucukgoze *et al*. [36], while green triangles give the results of the stochastic translational model with aminoacyl tRNA synthetase kinetics following the reaction scheme in Figures C and D at a growth rate of *µ* = 0.41 h^*−*1^.

**Table A Classification and Properties of Aminoacyl tRNA Synthetases from *E. coli***. For each of the 23 genes encoding an aminoacyl tRNA synthetase, the class, number of subunits which make up the functional enzyme, and if the enzyme has been observed to have burst kinetics or editing, is shown.

**Table B Kinetic parameters for Class I aminoacyl tRNA synthetase models**. Labels of the reactions correspond to those shown in Supplementary Figure C.

**Table C Kinetic parameters for Class II aminoacyl tRNA synthetase models**. Part 1 of 2. Labels of the reactions correspond to those shown in Supplementary Figure D.

**Table D Kinetic parameters for Class II aminoacyl tRNA synthetase models**. Part 2 of 2. Labels of the reactions correspond to those shown in Supplementary Figure D.

**Table E Estimates of AARS Activity *in vivo***. For each of the AARS enzymes, the activity (i.e. tRNA turnover rate *r*) of a single enzyme is estimated in *E. coli* cells growing at *µ* = 0.69 h^*−*1^ based on amino acid usage and the number of AARS. Amino acid usage for the Mori *et al*. data [31] was calculated computationally using a model of *in vivo* translation [24] and the fractional amount of tRNAs in ternary complex needed for optimal translation, while the amino acid usage for Jakubowski was taken from their experimental amino acid radio-labelling measurements [41].

**Table F Average Number of Proteins, tRNA, and Ribosomes in *E. coli* Cells at Different Growth Rates**. Protein numbers are taken from proteome measurements from several groups [26, 30, 31] and reported here as number per cell. The specific data for each growth rate are; *µ* = 0.41 h^*−*1^ Valgepea *et al*. 2013 [30], *µ* = 0.69 h^*−*1^ and *µ* = 1.04 h^*−*1^ Mori *et al*. 2021 [31], *µ* = 1.98 h^*−*1^ Li *et al*. 2014 [26]. The number of tRNA and Ribosomes (per cell) are taken from Dennis and Bremer [27]. The doubling time in minutes is computed from the growth rate using *τ* = 60 ln(2)*/µ*.

**Table G Average number of tRNAs per cell and their codon recognition**.

Data for the number of tRNAs that are in free ternary complex at a growth rate of *µ* = 0.41 h^*−*1^ (*τ* = 100 min). The numbers that give the optimal translation rate have been computed from Eq. 3 and the procedure outlined in section 4, and the total tRNA numbers are taken from Dong *et al*. [28]. The values used in the model are the result of the fitting procedure in Section 7.

**Table H Concentrations of Amino Acids in *E. coli***. Experimental measurements of intracellular amino acid concentrations are shown for Bennett *et al*. [20], and minimal media (MM) and minimal media plus amino acids (MM+AA) from Avcilar-Kucukgoze [36]. The values used in the model for the MM+AA scenario is shown in the final column. (n.d. = not determined)

**Table I Optimized numbers of Aminoacyl tRNA Synthetases in Exponentially Growing *E. coli* and optimized** *k*_*cat*_ **and** *K*_*m*_ **values**. The optimized average number of AARS enzymes are shown for different growth rates, *µ* = 0.41 h^*−*1^ (*τ* = 100 min), *µ* = 0.69 h^*−*1^ (*τ* = 60 min), *µ* = 1.04 h^*−*1^ (*τ* = 40 min), and *µ* = 1.73 h^*−*1^ (*τ* = 24 min). Corresponding optimized *k*_*cat*_ and Michaelis–Menten parameters for tRNA 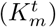 and amino acid 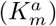 are given in the right hand columns.

**Table J Optimized numbers of tRNAs per cell and their codon recognition**. The optimized total tRNA numbers are shown for the Michaelis–Menten model and the full kinetic model where AARS enzymes are modelled as having reaction scheme according to Figure C for Class I enzymes or Figure D for class II enzymes.

**Table K Model predictions of AARS Activity *in vivo***. For each of the AARS enzymes, the activity (i.e. tRNA turnover rate) of a single enzyme *r* per second is estimated in *E. coli* cells growing at *µ* = 1.04 h^*−*1^ (*τ* = 40 min) based on amino acid usage and the number of AARS. Amino acid usage for the data in this work was calculated computationally using a model of *in vivo* translation [24], while the amino acid usage and AARS numbers for Choi and Covert were taken from supplementary material Tables 8 and S3 [10].

