## Supplementary material for "An empirical model of aminoacylation kinetics for *E. coli* class I and II aminoacyl tRNA synthetases": S1 Text

### 1. Variation of Kinetic Parameters for Class I and II Enzyme Models

In this section, a systematic variation of the kinetic parameters is performed in order to identify the subset which have the largest effect on $k_{cat}$ and $K_m$ values. This is done for both pyrophosphate exchange kinetics and for the overall aminoacylation reaction. The effects of kinetic parameters on the overall chemistry rate ($k_{chem}$) and the single turnover rate for aminoacyl transfer ($k_{tran}$) are also examined. This allows for the identification of the critical parameters which effect all the features of aminoacyl tRNA synthetase kinetics that have been experimentally observed.

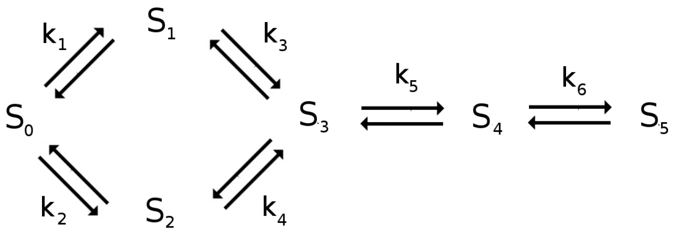


**Figure A. Kinetic reaction scheme for the pyrophosphate exchange reaction.** State $S_0$ represents the AARS enzyme, while $S_1$ and $S_2$ are the amino acid and ATP bound enzymes, respectively. State $S_3$ is the enzyme with both amino acid and ATP bound, $S_4$ is the adenylate and pyrophosphate bound state, while state $S_5$ is the adenylate bound enzyme state.

**Effect of Varying Kinetic Parameters on Pyrophosphate Exchange.** Figure A gives the reversible reaction network that is used to simulate pyrophosphate exchange. Although there are a total of 12 parameters that can be varied, the 12 dimensional search space can be greatly simplified to a smaller region which reproduces the experimentally observed $k_{cat}$ and $K_m$ values. Specifically, I will reduce the search space to a 4-dim space encompassing the dissociation constants for amino acid and ATP, $K_{d1}$, $K_{d2}$, and the amino acid activation rates $k_{5f}$ and $k_{5b}$. Although there are probably other regions of the parameter space which will reproduce the experimentally observed $k_{cat}$ and $K_m$'s, this region of parameter space is consistent with kinetic rate constants that have been explicitly measured for some tRNA synthetases (see supplementary S2 spreadsheet and citations within). Thus, this smaller region of parameter space should be sufficient for the construction of empirical kinetic aminoacyl tRNA synthetase models.

Examining Figure A, it is clear that potential parameters that will likely effect the $K_m$ values associated with pyrophosphate exchange are the dissociation constants for amino acid binding ($K_{d1} = k_{1b}/k_{1f}$ and $K_{d4}$) and ATP binding ($K_{d2} = k_{2b}/k_{2f}$ and $K_{d3}$). Similarly, $k_5$ and $k_6$ are likely to effect $k_{cat}$ values. As a first examination of the parameter space, I set $K_{d1} = K_{d4}$ and $K_{d2} = K_{d3}$ while varying $K_{d1}$ and $K_{d2}$ between 10 and 1000 $\mu$M. In addition I fix $k_{5f} = 180s^{-1}$ and $k_{5b} = 400s^{-1}$ with $k_{6f} = 5\ \mu M^{-1}s^{-1}$ and $k_{6b} = 3000s^{-1}$. The results of this variation are shown as contour plots in Supplementary Figure B (a-c). As expected, varying the $K_d$ for ATP or amino acid altered the corresponding $K_m$ value and suggests that for pyrophosphate exchange, $K_d \approx K_m$. Varying of the forward rates ($k_{1f}$ and/or $k_{2f}$) had negligible effect on either $k_{cat}$ or $K_m$ establishing that the predominate effect on the $K_m$ values in the pyrophosphate exchange reaction is due to the overall $K_d$ of amino acid or ATP.

Next, I fix $K_{d1} = K_{d4} = 54\ \mu$M and $K_{d2} = K_{d3} = 600\ \mu$M along with $k_{6f} = 5\ \mu\ M^{-1}s^{-1}$ and $k_{6b} = 3000s^{-1}$, and systematically vary $k_{5f}$ and $k_{5b}$ between values of 10 and 400 $s^{-1}$. The contour plots in Supplementary Figure B (d-f) show the effects on $k_{cat}$ and $K_m$ values in the pyrophosphate exchange reaction. As can be seen, $K_m$ increases toward the $K_d$ value of the substrate as $k_{5f}$ becomes substantially less then $k_{5b}$. In contrast, the overall $k_{cat}$ value increases as $k_{5f}$ increases.

Finally, I examine varying $k_{6f}$ from 0.1 to 200 $\mu$M$^{-1}s^{-1}$ and $k_{6b}$ between 10 and 2000 $s^{-1}$. The values of $K_{d1} = K_{d4} = 54$ $\mu$M and $K_{d2} = K_{d3} = 600\ \mu$M are fixed along with $k_{5f} = 180s^{-1}$ and $k_{5b} = 400s^{-1}$. As can been seen from the contour plots in Supplementary Figure B (g-i), there is little effect on $K_m$ values for ATP or the amino acid, while $k_{cat}$ values start to plateau when $k_{6b} > 1000s^{-1}$. Moreover, there is limited effect on $k_{cat}$ via variation of $k_{6f}$. Thus, $k_{6f}$ can be chosen to match the measured pyrophosphate dissociation constant which is in the range of $320 - 500\mu$M (1, 2). All together this demonstrates that, by altering the $K_d$'s for the amino acid and ATP, along with $k_{5f}$ and $k_{5b}$, one can obtain $k_{cat}$ values for pyrophosphate exchange in the range of $20 - 100\ s^{-1}$ and $K_m$s between 0 and the $K_d$ of the substrate.

**Effect of Varying Kinetic Parameters on Aminoacylation.** Figure C gives the reversible reaction network that is used to simulate the overall aminoacylation reaction of an aminoacyl tRNA synthetase for class I enzymes. Here there are a total of 44 parameters that can be varied to potentially effect overall aminoacylation kinetics. However, some of these rates, in particular the binding of tRNA and aa-tRNA to the enzyme (rates $k_0^x$ with $x = a, b, c, d, e$ and $k_{11}$, respectively) and the transfer rate ($k_{8f}$), have usually been experimentally measured. Moreover, if one first adjusts the 12 parameters $k_1$ to $k_6$ to fit pyrophosphate exchange and assumes that these rates in the presence of tRNA are the same, i.e. $k_1 = k_1^t$ etc., then this leaves just $k_7$, $k_9$ and $k_{10}$ to vary.

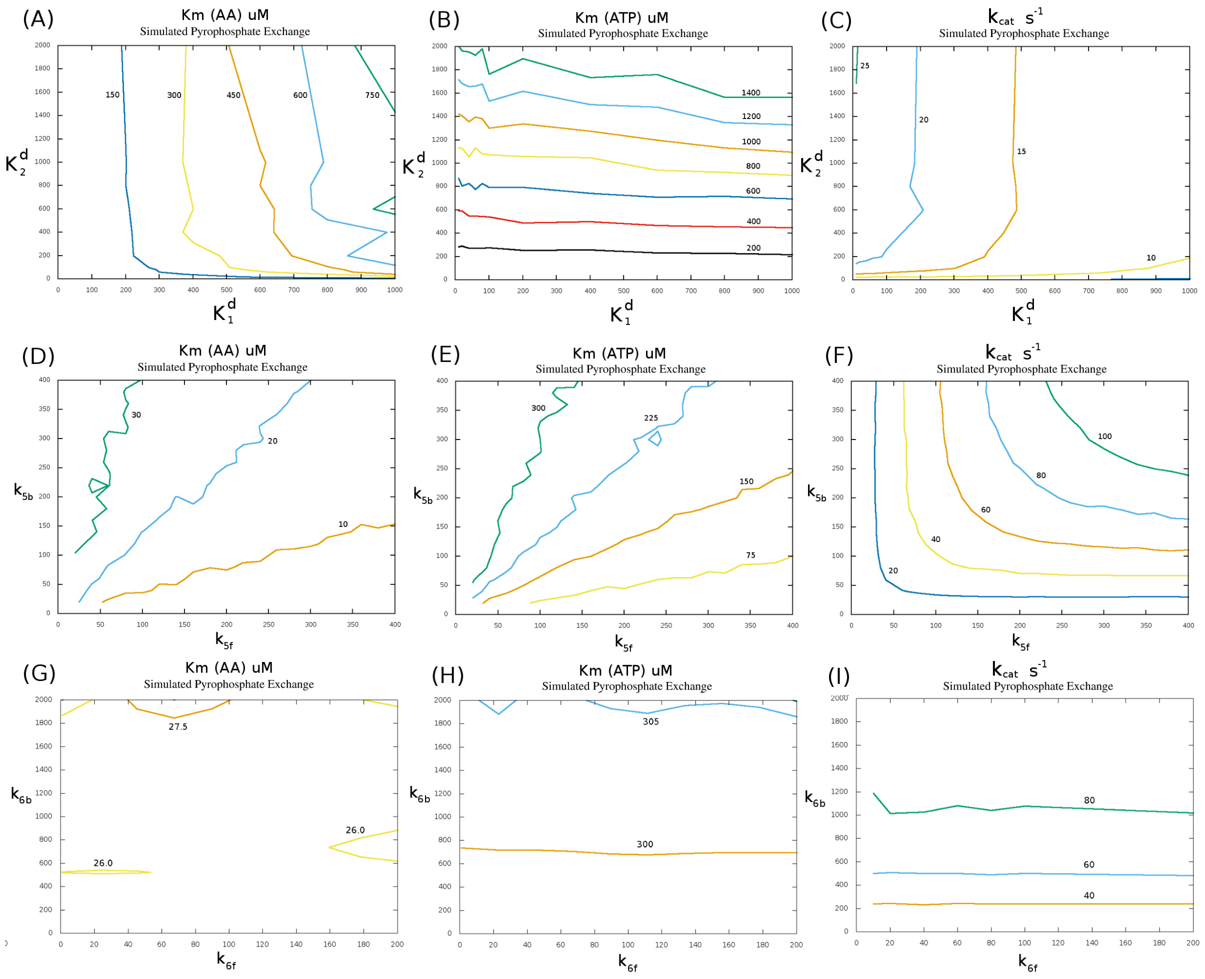


**Figure B. Effect of parameter variation on pyrophosphate exchange kinetics.** (A-C) Effect on Michaelis–Menten constant $K_m$ and pyrophosphate $k_{cat}$ as the amino acid and ATP dissociation constant is varied. (D-F) Effect on the Michaelis–Menten constant $K_m$ and pyrophosphate $k_{cat}$ as the amino acid activation rates $k_{5f}$ and $k_{5b}$ are varied. (G-I) Effect on the Michaelis–Menten constant $K_m$ and pyrophosphate $k_{cat}$ as the pyrophosphate release rates $k_{6f}$ and $k_{6b}$ are varied.

The rate $k_{10}$ governs the reversible binding of AMP to the enzyme after amino acid transfer to the tRNA. This dissociation rate has been estimated based on experimental observations to have a range of $K_d = k_{10b}/k_{10f} \approx 1200 - 2400\mu$M (2, 3). Variation of this parameter has negligible effect on $K_m$ or $k_{cat}$ (obviously as long as AMP release is greater than the overall catalysis rate - $k_{10b} > k_{cat}$) but alters the overall steady-state percentage of charged tRNAs in the presence of different concentrations of AMP. Values of this parameter will be important for insuring tRNAs charge in the presence of cellular concentrations of free AMP. Total AMP concentrations in *E. coli* have been estimated at 180-250 $\mu$M (4), with free AMP concentrations lower then this. Since this parameter has negligible effect on $K_m$ and $k_{cat}$, I set $k_{10f} = 1\ \mu M^{-1}s^{-1}$ and $K_{10b} = 2400\ s^{-1}$ to insure AMP dissociation does not inhibit the overall aminoacylation rate.

This leaves just $k_7$ and $k_9$ to vary in order to attempt to fit the $k_{cat}$ and $K_m$ values for aminoacylation. I relax this condition and additionally allow $k_6^t$, i.e. pyrophosphate release in the presence of uncharged tRNA, to also vary. Thus, the 38 dimensional space has been potentially limited to a much smaller 6 dimensional space for fitting to experimental $k_{cat}$ and $K_m$ values. The two reaction routes through aminoacylation of tRNA thus comprise, (1) a route where $PP_i$ is first released prior to the transfer of amino acid to the tRNA (governed by rate $k_6^t$) and (2) a route where $PP_i$ is not released until after transfer of the amino acid to the tRNA (governed by rates $k_7$ and $k_9$). I have left both of these routes in for two reasons. First, Kern and Lapointe (5) noted that, for some tRNA synthetases, presence of $PP_i$ may result in conformational changes to the enzyme which bring tRNA closer to the reaction center where the adenylate is present. Second, when I set $k_{7f} = k_{7b} = 0$, forcing $PP_i$ release prior to amino acid transfer, I have found that it is very difficult to reproduce both $k_{chem}$ and $K_m$ values that have been measured experimentally for some tRNA synthetase enzymes. Essentially, a slow release of $PP_i$ is (in general) required to move $K_m$ value closer to the $K_d$ value for the substrate. This has the added problem that it forces $PP_i$ release to become the rate limiting step. For enzymes such as cysRS which display burst kinetics, this will result in the disappearance of the burst kinetics in the model. Pope *et al.* noticed a similar issue when they attempted to theoretically simulate ileRS aminoacylation for *S. aureus* based on an extensive set of experimentally measured kinetic rates for the enzyme (6). In this work they noticed that, unless $PP_i$ was modelled as being released *after* transfer of the amino acid to the tRNA, $K_m$ values for ATP and isoleucine where far below what was measured experimentally. Thus, for all these reasons, I have included a reaction path which allows amino acid transfer in the presence of pyrophosphate.

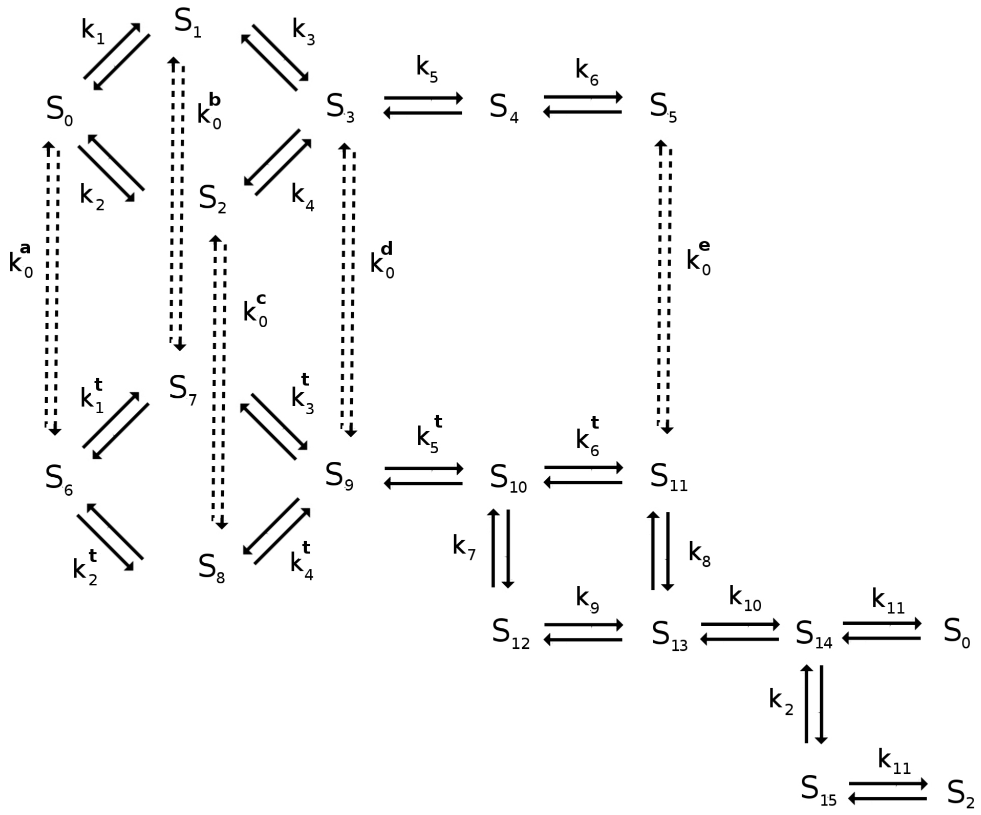


**Figure C. Kinetic reaction scheme for Class I aminoacyl tRNA synthetases.** Reaction scheme diagram labelling the individual kinetic reactions in the aminoacylation reaction for the class I aminoacyl tRNA synthetase enzymes. Individual kinetic rates for each reaction can be found in Supplementary Table B.

For the class II reaction scheme shown in Figure D, the binding of substrates can occur at two different sites. I model the tRNA charging kinetics based on the flip-flop mechanism detailed by Guth et al. (7). This mechanism is based on experimental observations that tRNA charging occurs first in site 1, followed by activation of the amino acid and formation of the adenylate before the charged tRNA in site 1 is released. I model the charged tRNA as being able to leave site 1 as soon as the adenylate is formed in site 2. The mechanism and rates follow similar fitting strategy to that of class I enzymes, as one first uses any experimental data on single turnover kinetics and pyrophosphate exchange to fit the kinetic parameters for site 1 of the enzyme. The remaining kinetic rates for site 2 can be varied straightforwardly following a similar scheme to that of pyrophosphate exchange to fit the model to the experimental data for $K_m$ and $k_{cat}$ of aminoacylation.

### 2. Kinetic Parameters for Aminoacyl tRNA Synthetases

Following the procedure outlined in the previous section, I have identified a set of kinetic parameters which reproduce the experimentally observed $k_{cat}$ and $K_m$ values for each of the 20 aminoacyl tRNA synthetases. For most of the enzymes, there are multiple measurements of both $k_{cat}$ and $K_m$ for the pyrophosphate exchange and aminoacylation reactions from several different experimental groups (see S2 spreadsheet). In these cases, after removing outliers, I have used the average of the $k_{cat}$ and $K_m$ data to fit the empirical models to. It is important to note, that some of the fitted $k_{cat}$ values for the aminoacylation reaction are above the average of the measurements, and occasionally outside the range of values found experimentally. This is to insure that the overall turnover rate supported by the AARS enzyme is sufficiently fast to support the turnover rate needed for exponential growth in *E. coli.* Table B and Tables C/D give the kinetic rates for class I and class II aminoacyl tRNA synthetases, respectively, which fit the experimental measurements reported in supplementary S2 spreadsheet. Information on the values of $k_{cat}$ and $K_m$ that the empirical model produces for each of the enzymes can also be found in supplementary Table I.

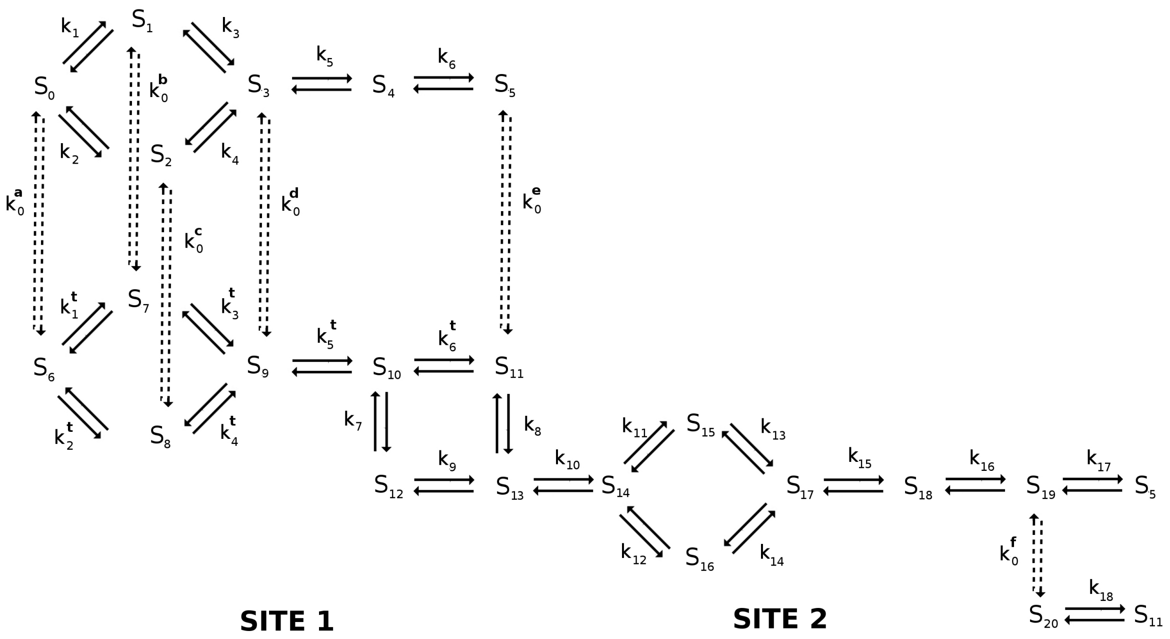


**Figure D. Kinetic reaction scheme for Class II aminoacyl tRNA synthetases.** Reaction scheme diagram labelling the individual kinetic reactions in the aminoacylation reaction for the class II aminoacyl tRNA synthetase enzymes. Individual kinetic rates for each reaction can be found in Supplementary Tables C and D.

### 3. Calculation of *in vivo* Amino Acid Usage and tRNA Turnover Rates

In this section I discuss how the amino acid usage and tRNA turnover rates are computed from computational simulations of ribosome translation that would occur *in vivo*. Additionally, recent proteomics data of AARS numbers in *E. coli* cells growing at different rates are compared with previous 2-D electrophoresis data from Neidhardt *et al.* (8).

**Estimation of AARS Numbers from Proteomics Data.** Jakubowski and Goldman (9) previously measured amino acid incorporation into protein using $^3$H labelled amino acids and used measurements of AARS numbers from Neidhardt *et al.* (8) to estimate average tRNA charging rates. Based on the measurements of AARS numbers using 2-D gel electrophoresis data from Neidhardt *et al.* and the estimated tRNA charging rates computed in (9), it has been suggested that the $k_{cat}$ values measured experimentally *in vitro* for most AARSs are too small to support the rates of tRNA charging that are estimated for exponentially growing *E. coli* (10). However, it is important to note that Neidhardt *et al.* stated in their discussion that their results would only reflect the actual numbers of AARS in cell if one accepted that they achieved near 100% efficiency in extraction of AARS enzymes from cell lysates. Thus, it is reasonably possible that the Neidhardt measurements are underestimated along with several of the amino acid incorporation rates measured by Jakubowski and Goldman's (9) being overestimated. This would result in an overall overestimate of the expected tRNA charging rate *in vivo*.

Indeed, examining the data in Table E reveals the amount of AARS enzyme reported in Jakubowski and Goldman (9) is substantially lower then what was measured more recently by Mori *et al.* (11). Moreover, while the Jakubowski and Goldman measurements of amino acid usage for several amino acids (e.g. valine, isoleucine, leucine, tryptophan, proline, histidine and lysine) are very close to theoretical estimates based on the number of translating ribosomes, there are several which are substantially different (glutamine, asparagine). In fact, the Jakubowski and Goldman data suggest that both cystine and tryptophan are more frequently incorporated into *E. coli* proteins then glutamine or asparagine, which is contradictory to the observation that cystine and tryptophan are the most rare amino acids in *E. coli* proteins. This suggest that some of these measurements may not be completely reliable. Fortunately over the past decade, several experimental groups have measured absolute concentrations of proteins in exponential growing *E. coli* cells in various growth conditions using either liquid chromatography mass spectroscopy or ribosome profiling methods (11–15). These more recent proteomics data provide additional data that can be compared with Neidhardt *et al.* and give the average numbers of AARSs in exponentially growing *E. coli* cells covering four different growth rates. Table F shows a selection of the data used in this study while the complete proteomics data set of 12 measurements can be found in Tab 2 of the supporting S2 spreadsheet.

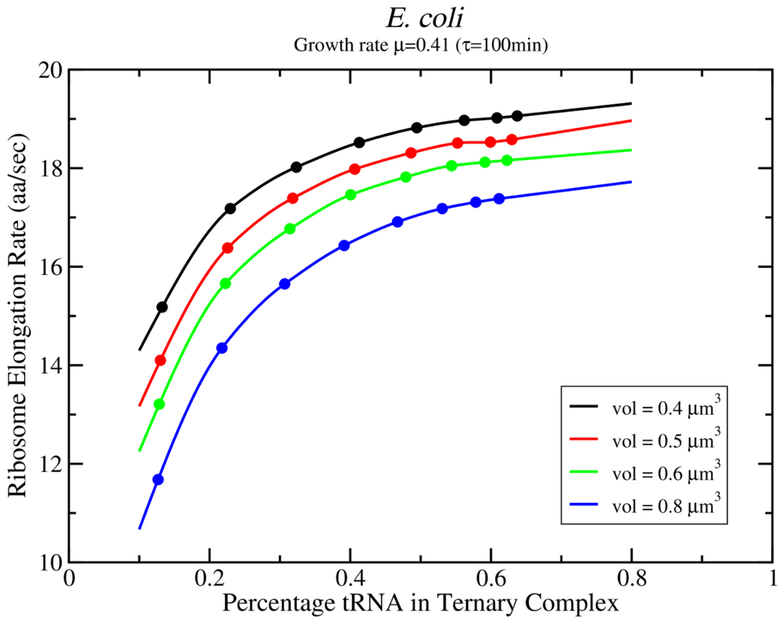


**Figure E. The dependence of the translation elongation rate on total ternary complex concentration.** Stochastic simulations of translation *in vivo* are used to calculate the average elongation rate of ribosomes in *E. coli* for different cellular volumes verses the percentage of the total amount of tRNA in ternary complex. The fractions of each tRNA isoacceptor in ternary complex set to the optimal values listed in Table G (see section 4 for further details).

**Estimation of Amino Acid Usage from Simulations of *in vivo* Translation.** One way to reasonably estimate the amino acid usage rate is by taking the total number of elongating ribosomes multiplied by the average elongation rate and the average frequency of an amino acid in *E. coli* proteins. Following standard predictions of Dennis and Bremer (16), the total number of elongating ribosomes can be assumed to be 80% of the total number of ribosomes in the cell which allows for a 5% fraction to be terminating or initiating with the remaining 15% free. Regarding the average elongation rate of ribosomes, this has been estimated by Dennis and Bremer (16) to be 15, 18, and 22 amino acids per second for growth rates of $\mu$ =0.41, 0.69, and 1.98 $h^{-1}$, respectively. However, Dai *et al.* (17) have measured the elongation rates to be slightly slower at 12, 15, and 18 amino acids per second based on translation of LacZ mRNAs. Thus for a given *E. coli* growth rate (*e.g.* 0.41 $h^{-1}$), one could take the average estimated elongation rate (12 aa/sec) and multiply by the active fraction of ribosomes and the average frequency of amino acids to obtain the amino acid usage rate $a_i$.

Despite the simplicity of the calculation, there are two issues with this approach. First, the elongation rate and usage rate of individual tRNA isoacceptors is dependent on the tRNA bias in free ternary complex in the cell, as well as the relation of this tRNA bias to the overall codon frequency in the transcriptome (see discussion in section 4 on tRNA bias below). Second, the elongation rate at specific growth rates is further complicated by experiments which observed an increased protein production rate following a nutrient up-shift (18). In these experiments, initially slow growing *E. coli* cells (initial doubling time $\tau_i = 90$ min) had their medium supplemented with glucose and amino acids resulting in an up-shift in protein production and growth rate (final doubling time $\tau_f = 28$ min). Interestingly, the up-shift in protein production occurred within a few minutes (18), faster then would be needed to synthesise more ribosomes, mRNA, tRNA, or AARS enzymes. This suggests that nutrient up-shift causes an increase in the elongation rate and active ribosome fraction to the levels that would be observed in the fastest growing cells, resulting in an overall increase in the protein production rate. These experiments suggest that the number of AARS enzymes in the cell need to be capable of supporting a tRNA charging rate to enable the maximum elongation rate that would occur if a nutrient up-shift took place.

However, what is the maximum elongation rate that is achievable? To answer this question, I have simulated the translation that would occur *in vivo* in *E. coli* cells during exponential growth using my stochastic model of translation (19, 20). This model simulates all known kinetic steps involved in initiation, elongation, and termination in *E. coli* on the full transcriptome of mRNAs that would be present in the cell. For instance, at a growth rate of $\mu = 0.41$, my model simulates the movement of $\approx$ 8000 ribosomes on a transcriptome of $\approx 10^6$ nucleotides. Moreover my stochastic model does not enforce an exact elongation rate but allows the elongation rate to emerge due to tRNA concentration and competition effects between isoacceptors. Using the optimal tRNA bias (see section 4 below) along with a transcriptome based on mRNA seq data from Li (13), I have calculated the elongation rate as a function of the percentage of the total tRNA that is in ternary complex as shown in Figure E. As can be seen, the elongation rate peaks when roughly 50-60% of the available tRNA is in free ternary complex. At concentrations of ternary complex higher then this, the elongation rate is limited by the time to accommodate the amino acid and translocate the ribosome.

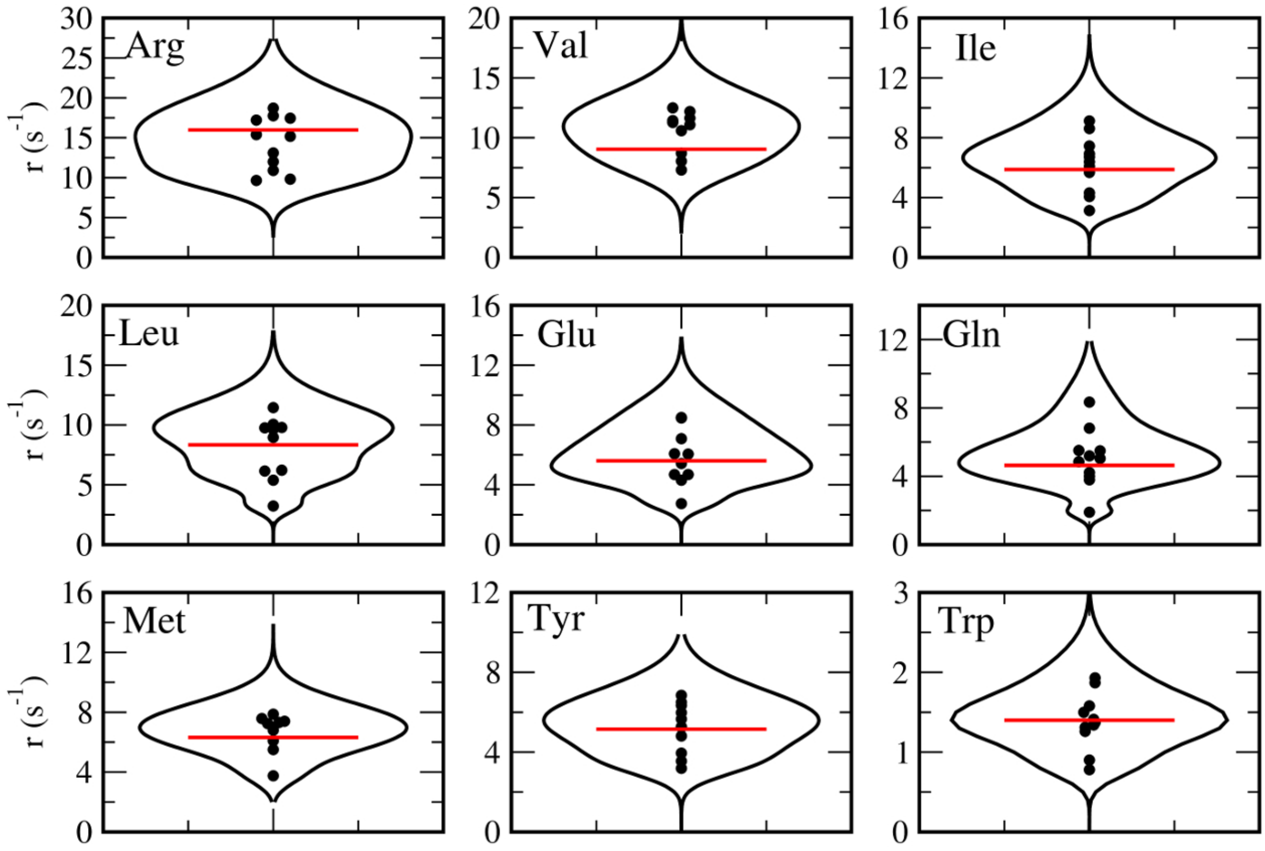


**Figure F. Estimated average tRNA turnover rates for class I aminoacyl tRNA synthetases.** Violin plots for the AARS tRNA turnover rates are shown for each of the class I enzymes, with cysRS shown in Figure 5 of the main text. Each of the individual proteomics data points used to construct the violin plot are shown (black dots) along with the value used for optimization (red line).

In summary, my stochastic ribosome model along with the following three settings are used to estimate both the usage of amino acids and tRNA isoacceptors; (1) the optimal tRNA fractions are used for the amount of each tRNA in free ternary complex, (2) estimates of the average cellular volumes of $v = 0.45, 0.65, 0.90, 1.95$ $\mu$m$^3$ are used for growth rates of $\mu = 0.41, 0.69, 1.04, 1.73$ (see discussion on cellular volume estimates below) and, (3) half of the available tRNA is forced to reside in free ternary complex. For example, at a growth rate of $\mu = 0.41$ h$^{-1}$, 37000 tRNAs out of a total of 74000 are forced to reside in free ternary complex (cf. Table F). Amino acid and tRNA isoacceptor usage rates are then calculated based on averages over 30 minutes of translation in a typical *E. coli* cell containing the appropriate number of ribosomes and mRNAs for a given growth rate.

**Estimation of AARS Activity *in vivo*.** Using the amino acid usage rate estimates from the computer simulations and the number of AARS from proteomics data, I have estimated the tRNA turnover rates ($r_i$) *in vivo* of the $i = [1, 20]$ aminoacyl tRNA synthetase enzymes from *E. coli*. For each proteomics data point $k$, the $r_i(k)$ is calculated from

$$r_i(k) = \frac{a_i}{n_i(k)} \, ,$$

where $n_i(k)$ is the number of enzymes and $a_i$ is the amino acid usage calculated from the theoretical translation simulations on the full transcriptome. Supporting Table E gives an example of the results for the proteomics data from Mori *et al.* for the *E. coli* growth rate of $\mu = 0.69$ h$^{-1}$ (11).

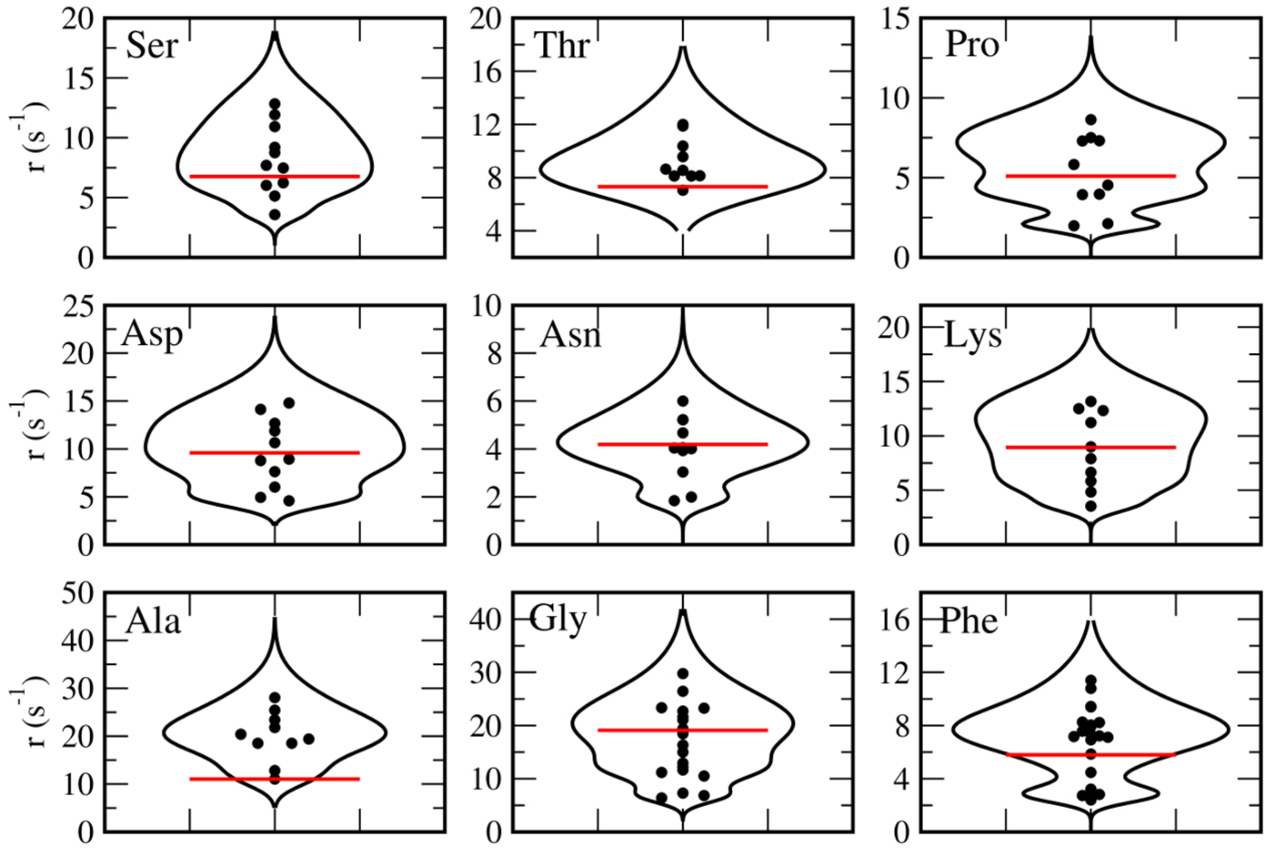


**Figure G. Estimated average tRNA turnover rates for class II aminoacyl tRNA synthetases.** Violin plots for the AARS tRNA turnover rates are shown for each of the class II enzymes, with hisRS shown in Figure 5 of the main text. Each of the individual proteomics data points used to construct the violin plot are shown (black dots) along with the value used for optimization (red line).

The full set of proteomics data covering 12 measurements was used to construct violin plots shown in Supporting Figures F and G along with those in Figure 5 in the main text for cysRS and hisRS. As can be seen in the figures, consensus turnover rates emerge for each of the aminoacyl-tRNA synthetase enzymes, represented by peaks in the violin plots, with a few AARS enzymes (*e.g.* proRS) having more then one potential turnover rate. Red lines in the figures represent the optimised turnover rate that have been used to calculate the average number of enzymes in an *E. coli* cell. Overall there is a good agreement with the consensus turnover rate with the optimised rate that has been used in the simulations with the exception of alaRS. Here alaRS turnover rate was lowered to approximately 10 s$^{-1}$ as single turnover experiments determined $k_{tran} = 16.4$ s$^{-1}$ for *E. coli* alaRS (21) which places an upper limit on $k_{cat}$ since it is expected that $k_{cat} < k_{tran}$.

### 4. tRNA Bias in Ternary Complex Required For Maximal Ribosome Elongation Rate

In total, *E. coli* has 42 different tRNA isoacceptors which decode the 64 codons of the genetic code. During translation, tRNAs in ternary complex must bind to the ribosome and decode the codon in the A-site of the ribosome. This requires the ribosome to successfully identify the correct tRNA isoacceptor that can decode the current codon in the ribosome A-site from the set of all tRNAs that are present in the cell. Some codons such as CCG, can be decoded by more then one isoacceptor, in this case by either tRNA$^{pro1}$ or tRNA$^{pro3}$, while other codons have a unique isoacceptor which decodes it. Thus, the time for the correct codon to enter the A-site and decode the codon is directly related to (a) the probability of correctly identifying a tRNA from those present in the cell that is capable of decoding it and (b) the overall concentration of all combined tRNAs. The relative concentrations of the 42 different tRNAs in ternary complex is sometimes referred to as the tRNA bias, and adjusting these relative concentrations will impact on the average decoding time of a given codon due to alteration of the probability that the correct tRNA enters the A-site. Let $p_c$ denote the probability of picking out a correct tRNA from all tRNAs in ternary complex that can decode codon $c$, while $q_c = (1 - p_c)$ is the probability of picking out an incorrect tRNA. For example, codon CCG can be decoded by either tRNA$^{pro1}$ or tRNA$^{pro3}$. Thus, $p_{ccg}$ can be calculated as

$$p_{ccg} = \frac{1}{N}(N_{pro1} + N_{pro3}) = n_{pro1} + n_{pro3}\ ,$$

where $N$ is the total number of tRNA in ternary complex, $N_{pro1}$ is the total number of tRNA$^{pro1}$ in ternary complex. It is important to note that even though $p_c < 1$ for all codons, the sum

$$\sum_c p_c > 1\,,$$

since tRNA isoacceptors can decode more then one codon causing $n_{pro1}$ (as an example) to appear more then once in the sum. During decoding, selection of the correct tRNA will follow a Bernoulli probability distribution, and each codon will have its own Bernoulli probability $p_c$. Specifically, there will be $k$ attempts of the *wrong* tRNA binding to the A-site, followed by the $k+1$ th attempt in which the correct tRNA is bound to the ribosome to decode the codon. The total time $\tau(k)$ to identify and bind the correct tRNA where $k$ failures was made is given by

$$\tau(k) = (\tau_{on} + \tau_{off})k + \tau_{on}\,,$$

where $\tau_{on}$ is the average time for a tRNA to bind the ribosome and transfer to the A-site for decoding and $\tau_{off}$ is the average time for tRNA to unbind from the A-site and be released from the ribosome. Note that, in a first approximation, these times can be considered independent of the codon and tRNA isoacceptor. The average time to identify the correct tRNA to decode codon $c$ (denoted as $\tau_c$) is given as the expectation value of $\tau(k)$ over all possible number of $k$ failures, *i.e.*

$$\begin{aligned}
\tau_c = \langle \tau(k) \rangle_c &= \sum_{k=0}^{\infty} q_c^k p_c \tau(k) \\
&= (\tau_{on} + \tau_{off}) \sum_{k=0}^{\infty} k q_c^k p_c + \tau_{on} \sum_{k=0}^{\infty} q_c^k p_c\,.
\end{aligned}$$

Using the relations

$$\begin{aligned}
\sum_{k=0}^{\infty} q_c^k p_c &= \frac{p_c}{1-q_c} = 1\,, \\
\sum_{k=0}^{\infty} k q_c^k p_c &= \frac{1}{p_c} - 1\,,
\end{aligned}$$

one finds

$$\tau_c = \frac{(\tau_{on} + \tau_{off})}{p_c} - \tau_{off}\,.$$

The average *total* time to decode codon $c$ is the average time to bind the correct tRNA isoacceptor in the A-site ($\tau_c$) plus the average time to accommodate the amino acid and translocate the ribosome to the next codon ($\tau_a$). Thus, one can calculate the average elongation rate in codons per second as $\langle E \rangle = \frac{1}{T}$, where $T$ is the average time to decode any codon, i.e.

$$\begin{aligned}
T &= \sum_c \tau_c f_c + \tau_a \\
&= \tau_a - \tau_{off} + (\tau_{on} + \tau_{off}) \sum_c \frac{f_c}{p_c}\,,
\end{aligned}$$

where $f_c$ is the frequency of codon $c$ in the current transcriptome of mRNAs present in the cell. First I note that $\tau_a$ is a fixed value which depends only on rates of translocation and accommodation of the amino acid into the nascent peptide chain. Thus, it is clearly independent of the tRNA concentration and tRNA bias in ternary complex. Similarly, $\tau_{off}$ is also independent of tRNA concentration and can be effectively considered as a constant. Hence, altering the value of $T$ can only be done by either (a) affecting the time to bind any tRNA into the A-site of the ribosome ($\tau_{on}$) which is inversely proportional to the *total* tRNA concentration, or (b) by altering the tRNA bias in the ternary complex which will change the values of $p_c$ for all of the codons.

The goal is to minimize $T$ by identifying the optimal tRNA bias, i.e. the optimal fractional numbers (e.g. $n_{pro1} = N_{pro1}/N$) of tRNA in ternary complex. With respect to the fractional number of 42 tRNAs $n_i$, the function that should be minimised is

$$g(n_i, \lambda) = \sum_c \frac{f_c}{p_c} + \lambda \left( \sum_i n_i - 1 \right)\,,$$

with $\lambda$ the Lagrange multiplier. However, this function has no extrema due to the recognition of some codons by multiple tRNAs causing $g$ to be over constrained. To illustrate this problem, consider the derivative of $g$ with respect to the fractional number of tRNA pro1, pro2, and pro3 in ternary complex (denote these as $n_1, n_2, n_3$) which each decode 1,2, and 3 codons respectively. The $p_c$ terms for the four proline codons are

$$\begin{aligned} p_{cca} &= n_3 \\ p_{ccc} &= n_2 \\ p_{ccg} &= n_1 + n_3 \\ p_{ccu} &= n_2 + n_3 \, , \end{aligned} \qquad [1]$$

which gives the following partial derivatives of $g$

$$\begin{aligned} \frac{\partial g}{\partial n_1} &= -\frac{f_{ccg}}{p_{ccg}^2} + \lambda \\ \frac{\partial g}{\partial n_2} &= -\frac{f_{ccc}}{p_{ccc}^2} - \frac{f_{ccu}}{p_{ccu}^2} + \lambda \\ \frac{\partial g}{\partial n_3} &= -\frac{f_{cca}}{p_{cca}^2} - \frac{f_{ccg}}{p_{ccg}^2} - \frac{f_{ccu}}{p_{ccu}^2} + \lambda \, . \end{aligned} \qquad [2]$$

Setting these 3 derivatives to zero to solve for the minimum results in 3 coupled quadratic equations

$$\begin{aligned} \lambda &= \frac{f_{ccg}}{(n_1 + n_3)^2} \\ \lambda &= \frac{f_{ccu}}{(n_2 + n_3)^2} + \frac{f_{ccc}}{n_2^2} \\ 0 &= \frac{f_{ccu}}{(n_2 + n_3)^2} + \frac{f_{cca}}{n_3^2} \end{aligned}$$

Letting $x = n_3/n_2$, the last quadratic equation can be re-arranged to

$$1 + 2x + \left(1 + \frac{f_{ccu}}{f_{cca}}\right) x^2 = 0 \, .$$

Since $f_c \geq 0$ for all codons, then one can easily see that this quadratic equation has only imaginary solutions for $x = n_3/n_2$, implying that the three constraint conditions cannot be satisfied simultaneously. In general, this problem arises for any codon which is not decoded by a single tRNA, as is the case for the CCG and CCU codons in the proline example. If we instead had the situation where each sense codon is decoded by a unique tRNA, then $p_c = n_c$ for all codons and

$$\frac{\partial g}{\partial p_i} = -\frac{f_i}{n_i^2} + \lambda = 0 \, ,$$

which must be true for all $n_i$ when $g$ is at an extrema. Solving for $n_i$

$$n_i^2 = \frac{f_i}{\lambda} \, ,$$

and substituting into the constraint equation, one finds

$$\sum_c \sqrt{f_c} = \sqrt{\lambda} \, .$$

Since $n_i = p_i > 0$ for all codons, this implies that there is only one extrema for $g$ which is given by

$$n_i = \frac{\sqrt{f_i}}{\sum_c \sqrt{f_c}} \, . \qquad [3]$$

A similar formula where the optimal total concentration of tRNAs decoding codon $c$ was found to be proportional to the square root of $f_c$ was also derived using a different theoretical approach by Ehrenburg and Kurland (22) as well as Berg and Kurland (23).

One solution to the constraint problem is to enforce that the ratio between tRNA isoacceptors in ternary complex decoding a particular codon is a constant. As an example, consider the three tRNA isoacceptors (pro1, pro2, and pro3) which decode the four proline codons. Again, let the fractional numbers of each be denoted by $n_1, n_2, n_3$. Then one can construct the ratios

$$r_i = \frac{n_i}{n_1 + n_2 + n_3} = \frac{N_i}{N_1 + N_2 + N_3} \, ,$$

for each isoacceptor. Defining $n_{pro} = n_1 + n_2 + n_3$ allows one to write the fractional number of each tRNA isoacceptor as the ratio times the total fractional number of tRNAs that decode proline, $n_i = r_i n_{pro}$. Assuming that the ratios $r_i$ are fixed constants and that only $n_{pro}$ is variable, the probabilities in Eq. 1 become

$$\begin{aligned} p_{cca} &= r_3 n_{pro} \\ p_{ccc} &= r_2 n_{pro} \\ p_{ccg} &= (r_1 + r_3) n_{pro} \\ p_{ccu} &= (r_2 + r_3) n_{pro} \,. \end{aligned} \quad [4]$$

This means that only one partial derivative with respect to $n_{pro}$ needs to be taken, as opposed to three in Eq. 2, i.e.

$$\frac{\partial g}{\partial n_{pro}} = -\frac{f_{cca}}{p_{cca}^2} r_3 - \frac{f_{ccc}}{p_{ccc}^2} r_2 - \frac{f_{ccg}}{p_{ccg}^2}(r_1 + r_3) - \frac{f_{ccu}}{p_{ccu}^2}(r_2 + r_3) + \lambda \,.$$

Since $p_{cca}$ etc. are all proportional to $n_{pro}$, one can directly solve the constraint equation for $n_{pro}$ in terms of the constants $r_i$ and $\lambda$ and a exact solution for all of the tRNA fractions can be obtained.

Using this approach, I have calculated the tRNA fractions that would yield the optimal translation rate based on the codon usage (or codon bias) that has been measured in both Li *et al.* and Dong *et al.* (13, 24). Briefly, Li *et al.* used mRNA-seq to determine the average mRNA levels (in units of RPKM) that are present in exponentially growing *E. coli* cells. This information can be used to back construct a snapshot of the average transcriptome in the cell (see Methods in main text). Once a transcriptome is constructed, the codon usage can be compared with the codon usage which was computed separately by Dong *et al.* (24). Tab 3 in S2 spreadsheet shows that the codon usage as measured by Dong *et al.* is in excellent agreement with the codon usage computed from the transcriptome that I have constructed using mRNA-seq data from Li *et al.* (13).

The codon bias was then used to determine the codon frequencies $f_c$ and the tRNA fractions (cf. Table G) that would yield the optimal elongation rate. Tab 4 in supplementary S2 spreadsheet gives the amino acid usage rates per second when the tRNA concentrations in ternary complex are fixed to those required for optimal translations. As an example, supplementary Table G gives the number of tRNAs in ternary complex for a growth rate of $\mu = 0.41$ h$^{-1}$. The numbers of tRNAs in ternary complex used by the model and adjusted to take into account tRNA abundances measured by Dong *et al* (24) are also given in Table G while Tab 5 in supplementary S2 spreadsheet give the corresponding amino acid usage rates per second when tRNAs in ternary complex are fixed at these numbers.

### 5. Effect of tRNA Bias in Ternary Complex on tRNA Usage Rates

Given a codon (e.g. CCG) which is decoded by more then one tRNA isoacceptor (tRNA$^{pro1}$ and tRNA$^{pro3}$), one might ask what would be the relative usage rates of each tRNA isoacceptor in decoding the CCG codon. More generally, since tRNA$^{pro3}$ decodes three codons (CCA,CCG, and CCU), it would be useful to obtain the average usage rate of tRNA$^{pro3}$ overall. To calculate this, denote the probability of decoding codon CCG by tRNA$^{pro3}$ as $P_{ccg}^3$. Since CCG can be decoded by two tRNAs, this probability is simply

$$P_{ccg}^3 = \frac{N_3}{N_1 + N_3} \,,$$

where $N_1$ is the number of tRNA$^{pro1}$ which are in ternary complex. The usage rate of tRNA$^{pro3}$ in decoding the CCG codon is then

$$U_{ccg}^3 = \alpha P_{ccg}^3 f_{ccg} \,,$$

where $f_{ccg}$ is the codon frequency of CCG in the transcriptome and $\alpha$ is a factor relating to the average translational speed of the ribosome so that $U_{ccg}^3$ has units of s$^{-1}$. Repeating the procedure for the other proline codons (CCA, CCC, and CCU), we obtain the total average usage rates of all three tRNAs as

$$\begin{aligned} U^1 &= \alpha \frac{N_1}{N_1 + N_3} f_{ccg} \\ U^2 &= \alpha \left[ \frac{N_2}{N_2 + N_3} f_{ccu} + f_{ccc} \right] \\ U^3 &= \alpha \left[ \frac{N_3}{N_1 + N_3} f_{ccg} + \frac{N_3}{N_2 + N_3} f_{ccu} + f_{cca} \right] . \end{aligned}$$

Since the CCA codon can only be decoded by tRNA$^{pro3}$, $N_3$ must be non-zero and we must enforce that $N_3 > 0$, with a similar argument for $N_2$. Let $x = N_1/N_3$ and $y = N_2/N_3 > 0$, then the usage rates can be rewritten as

$$\begin{aligned} U^1 &= \alpha \frac{x}{x+1} f_{ccg} \\ U^2 &= \alpha \left[ \frac{y}{y+1} f_{ccu} + f_{ccc} \right] \\ U^3 &= \alpha \left[ \frac{1}{x+1} f_{ccg} + \frac{1}{y+1} f_{ccu} + f_{cca} \right] . \end{aligned}$$

Finally, if we define $\beta$ as

$$\beta = \alpha(f_{ccg} + f_{cca} + f_{ccc} + f_{ccu}) \, ,$$

with

$$r_{ccx} = \frac{f_{ccx}}{(f_{ccg} + f_{cca} + f_{ccc} + f_{ccu})} \, ,$$

then $U^1 + U^2 + U^3 = \beta$. Codon usage rates (per million) for the CCA, CCC, CCG, and CCU codons are 5085, 1011, 25949, and 5558, respectively (see supplementary S2 spreadsheet, T3).

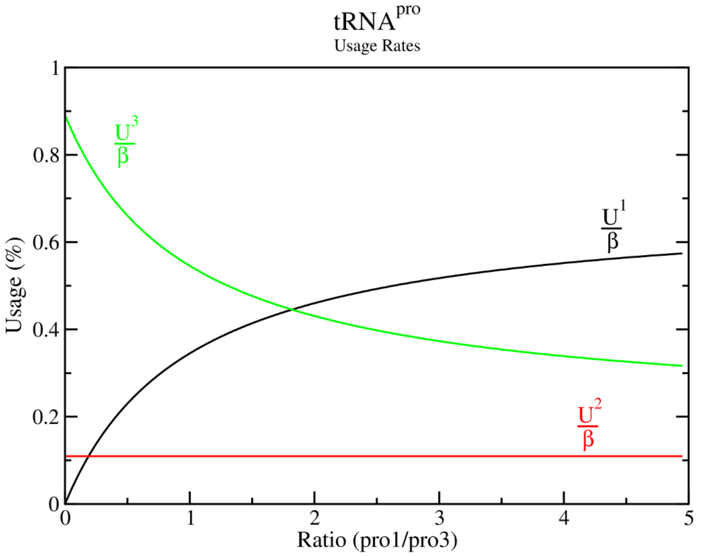


**Figure H. Average tRNA usage rates for the three proline tRNA isoacceptors.** Theoretical average tRNA usage rates for tRNA$^{pro1}$ (black line), tRNA$^{pro2}$ (red line) and tRNA$^{pro3}$ (green line) are shown using a fixed ratio of tRNA$^{pro2}$ to tRNA$^{pro3}$ of y = 1.25 in ternary complex. The graph is plotted as a function of the ratio of tRNA$^{pro1}$ to tRNA$^{pro3}$ in ternary complex.

Figure H shows how the relative usage of tRNA$^{pro1}$ vs tRNA$^{pro3}$ changes as the amount of tRNA$^{pro1}$ increases, assuming a fixed ratio of tRNA$^{pro2}$ to tRNA$^{pro3}$ of y = 1.25 in ternary complex. When the amount of tRNA$^{pro1}$ in ternary complex is zero, then roughly 90% of proline codons are decoded by tRNA$^{pro3}$ and 10% by tRNA$^{pro2}$. As the relative amount of tRNA$^{pro1}$ increases relative to tRNA$^{pro3}$, the usage rate of tRNA$^{pro1}$ steadily increases until it becomes the dominant tRNA that is used by the ribosome to decode proline codons. This effect is taken into account when fitting total tRNA numbers to the experimental measurements of Dong *et al.* (24) and for estimating the ratios for tRNA bias calculations in Eq. 4.

### 6. Estimation of Average E. Coli Cell Volumes at Different Growth Rates

Since I work with numbers of tRNAs and AARS enzymes, I need to know the volume of an average *E. coli* cell at various growth rates in order to fit to $K_m$ values which are typically measured in units of $\mu$M. Several experimental groups have measured the dimensions of *E. coli* cells (25–28), which as noted by Bremer and Dennis, are typically done by either a particle counter (Coulter counter) or electron microscopy (16). Bremer and Dennis report that several experimental measurements done with the particle counter method determined average cell sizes of 0.83 and 1.12$\mu$m$^3$ for cells with doubling times of $\tau = 46$ and 28 min, respectively. They also reported electron microscopy experiments which gave average cell volumes of 0.63 and 1.20$\mu$m$^3$ for cells with doubling times of $\tau = 60$ and 24 min, respectively. The results of their analysis suggested that cell volume increases exponentially with growth rate, with values of 0.50, 0.63 and 1.20 $\mu$m$^3$ for cells with doubling times of 100, 60, and 24 min respectively.

In this work I have taken the results from Bremer and Dennis along with several experimental groups (25–28) and obtained a consensus average volume measurement for exponentially growing *E. coli* cells. The consensus values that I have used in this work are 0.45, 0.65, 0.90, and 1.95 $\mu$m$^3$ for cells with doubling times of 100, 60, 40, and 24 min respectively. These values are similar to those plotted in supplementary figure 7 of Belliveau *et al.* (29) which also obtains consensus average volumes for exponentially growing *E. coli* cells from an analysis of multiple experimental measurements.

### 7. Optimization Procedure for kcat, Km, and tRNA Numbers

When optimizing $k_{cat}$ and $K_m$ values of AARS enzymes for computer simulations we need to ideally choose optimal values that result in the simulations of tRNA charging and ribosome elongation also being consistent with a variety of additional experimental observations. These include; (1) experimental measurements of the average total tRNA numbers in the cell, (2) proteomics data on the number of enzymes, (3) the percentages of charged tRNA in the cell, and (4) experimental measurements of average amino acid concentrations. Although there are 20 AARS enzymes, we are fortunate that each can be optimized separately as they are independent of each other. Although each AARS enzyme can be optimised independently, this optimization problem is still extraordinarily complex since we are attempting to match multiple experimental data points, each of which will effect the choice of $k_{cat}$ and $K_m$ values as well as have their own experimental errors. Moreover, as can be seen from the data on $k_{cat}$ measurements from a variety of experimental groups (main text, Figure 6), there is a substantial range of potential values making it difficult to pinpoint a precise one to choose.

As a starting point, let us assume that ATP and amino acid are saturating on the AARS enzyme. Then we essentially wish to fit

$$r = \frac{k_{cat} T_u}{K_m^t + T_u} \,,$$

where $r$ is the turnover rate for the AARS enzyme and $T_u$ is the quantity of *free* uncharged tRNA in the cell. Note that the turnover rate per enzyme $r$ depends on the amino acid usage rate and the number of enzymes, which is experimentally measured *in vivo*. Similarly, $k_{cat}$ and $K_m$ are measured by *in vitro* kinetic assays. Finally, the free amount of uncharged tRNA in the cell will determine the charging ratio and be dependent on the total number of tRNAs in the cell. Thus, each variable in the equation essentially needs to be chosen in some way which minimizes the error to these experimental data points. To simplify things, I fix $r$ for each enzyme using the consensus value from proteomics data (red lines in Figures F and G), except for alaRS, proRS, and pheRS, which are lowered to get a better consensus fit with single-turnover transfer rates ($k_{tran}$) measured for the enzymes.

If one now supposes that we know the quantity of *free* uncharged tRNA in the cell $T_u$, then one can re-arrange the above equation to the straight line equation

$$k_{cat} = a K_m^t + r \,, \tag{5}$$

where the slope of the line is given by $a = r/T_u$. Thus, given the amount of free uncharged tRNA in the cell and the turnover rate $r$, one can identify a pair of $k_{cat}$ and $K_m^t$ values on the line which minimize the error to experimental measurements. Hence, we can see that the critical (and most difficult) part of the fitting is the determination of the amount of free uncharged tRNA present in the cell.

Ideally, one would want to systematically vary the amount of free uncharged tRNA ($T_u$) as well as the amount of each tRNA in ternary complex ($T_{tc}$) for all 42 isoacceptors, looking for solutions which match the total amount of tRNA measured by Dong (24). The amount of uncharged free tRNA allows us to identify a best fit pair of $k_{cat}$ and $K_m^t$ which minimises their error with respect to experiment, while the amount of tRNA in ternary complex will set the amino acid usage rate and expected turnover rate. Then for each variation, the tRNA consumption rate can be calculated and an AARS enzyme model can be parameterised to reproduce $k_{cat}$ and $K_m$. Then finally the parameterised model can be tested that it reproduces the expected kinetics. In practice however, the fitting procedure can only be partially automated since the computational complexity of systematically varying the free uncharged tRNA concentration and re-building and testing a parameterised AARS model each time is too high. I have set out the three stage procedure that I have used below.

#### Stage 1: Initial parameterisation of AARS enzymes

Step 1: Use the optimal tRNA numbers of each isoacceptor in ternary complex (Table G) to estimate the upper bound on; (1) the average rates of tRNA usage and, (2) the average number of tRNA bound to ribosomes ($T_r$). This is done by fixing ternary complex concentration in my ribosome translation model (19), then averaging the resulting tRNA usages over 30 minutes of steady-state translation. Tab 4 in supporting spreadsheet S2 gives the computed upper bound amino acid usage rates. For different growth rates, an assumption that 50% of the total tRNA is in ternary complex is used so that the elongation rate is nearly maximized following Figure E.

Step 2: Use the turnover rates calculated from step 1 to estimate the number of enzymes $N$ using the consensus turnover rates from Figures F and G (red lines). As a first approximation, assume that the amount of free charged tRNA ($T_c$) is zero and estimate the number of tRNA bound to AARS as $T_b = 0.75N$, which tends to be a very good starting estimate. The amount of free uncharged tRNA is then estimated using $T_u = T_t - T_{tc} - T_r - T_b$), where $T_t$ is the total tRNA measured in Dong *et al.* (24), $T_{tc}$ is the total tRNA in ternary complex, and $T_r$ is the total bound to ribosome computed in step 1. Use the average experimental value for $K_m^t$ and Eq. 5 to obtain a $k_{cat}$ estimate. As an example calculation, at $\mu = 0.41$ h$^{-1}$, cysRS has (from Tab 8 in S2 spreadsheet) $T_t = 1625$, $T_{tc} = 451$, and $T_r = 96$, with a consumption rate of 731.5 per sec (from Tab 4 in S2 spreadsheet). Using the consensus turnover rate of 1.6 for cysRS (Figure 5 main text), the number of cysRS enzymes is $N = 457$ and $T_b = 343$. Thus, one expects that the number of free uncharged tRNA$^{cys}$ is approximately $T_u = 735$, or 2.71 $\mu$M using a cell volume of 0.45 $\mu$m$^3$. Applying Eq. 5 with the average $K_m^t = 1.06$ for cysRS (see cysRS tab in supplementary spreadsheet S2) gives $k_{cat} = 2.2$ s$^{-1}$.

Step 3: Fit parameters for the AARS model using the procedure in section 1 of supplementary S1 Text assuming $K_m^a$ and $K_m^{atp}$ are the experimental average. Test that the model can support the turnover rate computed in Step 1 assuming free concentrations of AMP and pyrophosphate of $[AMP] = 250\mu$M and $[PP] = 500\mu$M (4, 30, 31). Note: $k_{cat}$ may require a small increase at this stage to accommodate inhibition by AMP and pyrophosphate.

**Stage 2: Refinement of tRNA numbers in ternary complex**

Step 1: Adjust the number of tRNA in ternary complex by simulating the kinetics of all 20 enzymes undergoing constant tRNA consumption at the optimal rate. This returns target values for the amount of tRNA in ternary complex which will be the model fit (see Table G). Note this adjustment needs to be done since there are several highly expressed tRNAs (e.g. arg2 and leu1) that are have substantially more tRNA in ternary complex then is optimal.

Step 2: Repeat STAGE 1 procedure using the adjusted ternary complex numbers.

**Stage 3: Refinement of AARS models and total tRNA numbers**

Step 1: With models now constructed for each AARS based on adjusted ternary complex numbers, we can obtain a better estimate for the true amount of free uncharged tRNA. Using kinetic simulations of AARS kinetics with fixed tRNA consumption rates, fine adjustments are made to the total tRNA numbers in order to match Dong *et al.* and the overall % of tRNA that is charged.

Step 2: Construct a Michaelis–Menten model by using the $k_{cat}$, $K_m$ values and enzyme numbers in Table I. The total tRNA numbers are adjusted computationally in the ribosome translation model (19) until the average amount of tRNA in ternary complex matches the target (see supporting spreadsheet S2, Tab 8 for target tRNA numbers in ternary complex).

The results of the fitting procedure for $k_{cat}$, $K_m$, number of enzymes, and tRNA numbers are shown in Tables I and J. A direct comparison with measured tRNA numbers from Dong *et al.* (24) at a growth rate of $\mu = 0.41$ h$^{-1}$ can be found in Table G, or in supplementary spreadsheet S2, Tab 8 for other growth rates. While most of the tRNAs deviate by less then 20% from the Dong measurements, there are a few outliers with $\approx 60\%$ deviation. In particular tRNAs lys, asn, arg4, arg5, metM, pro3 and phe. Interestingly, as reported by Mohanty and Kushner (32), *E. coli* MG1655 expresses several tRNA as polycystronic transcripts which are then post-processed. For instance, ser3 (serV) and four copies of arg4 (argQ, argV, argY, argZ) are all expressed via a single polycistronic transcript. Since tRNA is a highly stable RNA in the cell (33), it is reasonable to expect that a ratio of arg2 / ser3 = 4 is maintained. Indeed, measurements by Dong *et al.* report values very close to this ratio. However, one copy of leu3 with two copies of metM, gln1, and gln2 are present on another polycistronic transcript suggesting a ratio of metM/leu3 $\approx 2$. This ratio is not seen in the Dong measurements. However, I find that enforcement of this ratio gives much better results. This accounts for the high differences between the model fit and measurements seen for pro3, metM, his, gln1, and gln2 tRNAs. For tRNAs lys, asn, and phe, I observed that the total quantities of tRNA needed to increase in order to account for the large proportion of tRNAs that are in complex with ribosomes and AARS enzymes.

### 8. Theoretical tRNA Charging Fractions

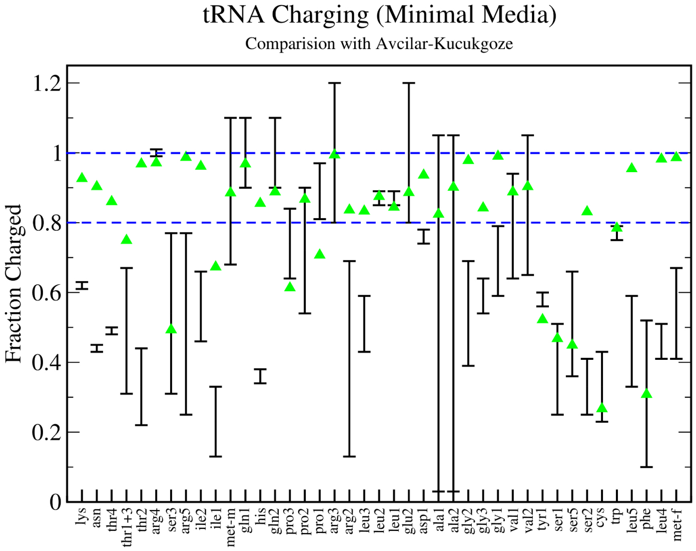


Figure I.**Theoretical and Experimental Estimates of tRNA Charging Fractions in Minimal Media.** The total fraction of tRNAs charged in exponentially growing *E. coli* cells. Black lines with error bars indicate the range of experimental measurement from Avcilar-Kucukgoze *et al.* (34), while green triangles give the results of the stochastic translational model with aminoacyl tRNA synthetase kinetics following the reaction scheme in Figures C and D at a growth rate of $\mu = 0.41$ h$^{-1}$.

**Table A. Classification and Properties of Aminoacyl tRNA Synthetases from *E. coli*. For each of the 23 genes encoding an aminoacyl tRNA synthetase, the class, number of subunits which make up the functional enzyme, and if the enzyme has been observed to have burst kinetics or editing, is shown.**

| AARS | Gene | Class | Subunits | Burst | tRNA Dep. | Editing |
|---|---|---|---|---|---|---|
| cysRS | cysS | I | 1 | Y | | |
| argRS | argS | I | 1 | Y | Y | |
| valRS | valS | I | 1 | Y | | Y |
| ileRS | ileS | I | 1 | | | Y |
| leuRS | leuS | I | 1 | Y | | Y |
| gluRS | gltX | I | 1 | Y | Y | |
| glnRS | glnS | I | 1 | Y | Y | |
| metRS | metG | I | 2 | Y | | Y |
| tyrRS | tyrS | I | 2 | Y | | |
| trpRS | trpS | I | 2 | Y | | |
| serRS | serS | II | 2 | | | Y |
| thrRS | thrS | II | 2 | | | Y |
| proRS | proS | II | 2 | | | Y |
| hisRS | hisS | II | 2 | | | |
| aspRS | aspS | II | 2 | | | |
| asnRS | asnS | II | 2 | | | |
| lysRS | lysS | II | 2 | | | Y |
| alaRS | alaS | II | 2 | | | Y |
| lysRS* | lysU | II | 2 | | | Y |
| glyRS$\alpha$ | glyS | II | 2 | | | |
| glyRS$\beta$ | glyQ | II | 2 | | | |
| pheRS$\alpha$ | pheS | II | 2 | | | Y |
| pheRS$\beta$ | pheT | II | 2 | | | Y |

**Table B. Kinetic parameters for Class I aminoacyl tRNA synthetase models. Labels of the reactions correspond to those shown in Supplementary Figure C.**

| Enzyme | $k_{1f}$ $1/\mu$Ms | $k_{1b}$ $1/s$ | $k_{2f}$ $1/\mu$Ms | $k_{2b}$ $1/s$ | $k_{3f}$ $1/\mu$Ms | $k_{3b}$ $1/s$ | $k_{4f}$ $1/\mu$Ms | $k_{4b}$ $1/s$ | $k_{5f}$ $1/s$ | $k_{5b}$ $1/s$ | $k_{6f}$ $1/\mu$Ms | $k_{6b}$ $1/s$ | $k^a_{0f}$ $1/\mu$Ms | $k^a_{0b}$ $1/s$ | $k^b_{0f}$ $1/\mu$Ms | $k^b_{0b}$ $1/s$ |
|---|---|---|---|---|---|---|---|---|---|---|---|---|---|---|---|---|
| cysRS | 0.24 | 14.4 | 6.00 | 3480 | 6.00 | 2320 | 0.24 | 9.6 | 160 | 400 | 6.00 | 3000 | 46.0 | 138 | 46.0 | 138 |
| argRS | 2.40 | 180 | 6.00 | 720 | 6.00 | 720 | 2.40 | 180 | 0 | 0 | 0 | 0 | 40.0 | 184 | 40.0 | 36.8 |
| valRS | 1.00 | 138 | 6.00 | 3000 | 6.00 | 2000 | 1.00 | 92 | 140 | 400 | 6.00 | 3000 | 80.0 | 30 | 80.0 | 30 |
| ileRS | 2.20 | 15 | 6.00 | 2400 | 6.00 | 2400 | 2.20 | 15 | 150 | 400 | 6.00 | 3000 | 120 | 138 | 120 | 138 |
| leuRS | 1.20 | 90 | 6.00 | 3300 | 6.00 | 3300 | 1.20 | 90 | 135 | 350 | 6.00 | 3000 | 40.0 | 168 | 40.0 | 56 |
| gluRS | 0.15 | 27 | 3.40 | 3740 | 3.40 | 3740 | 0.15 | 27 | 0 | 0 | 0 | 0 | 80.0 | 64 | 80.0 | 64 |
| glnRS | 0.20 | 216 | 5.00 | 3400 | 5.00 | 3400 | 0.20 | 216 | 0 | 0 | 0 | 0 | 120 | 42 | 120 | 42 |
| tyrRS | 0.60 | 15 | 4.20 | 3360 | 4.20 | 2688 | 0.60 | 12 | 160 | 320 | 6.00 | 3000 | 42.0 | 48 | 42.0 | 48 |
| trpRS | 0.30 | 13.5 | 8.00 | 2500 | 8.00 | 2500 | 0.30 | 13.5 | 320 | 400 | 6.00 | 3000 | 45.0 | 117 | 45.0 | 117 |

| Enzyme | $k^t_{1f}$ $1/\mu$Ms | $k^t_{1b}$ $1/s$ | $k^t_{2f}$ $1/\mu$Ms | $k^t_{2b}$ $1/s$ | $k^t_{3f}$ $1/\mu$Ms | $k^t_{3b}$ $1/s$ | $k^t_{4f}$ $1/\mu$Ms | $k^t_{4b}$ $1/s$ | $k^t_{5f}$ $1/s$ | $k^t_{5b}$ $1/s$ | $k^t_{6f}$ $1/\mu$Ms | $k^t_{6b}$ $1/s$ | $k^c_{0f}$ $1/\mu$Ms | $k^c_{0b}$ $1/s$ | $k^d_{0f}$ $1/\mu$Ms | $k^d_{0b}$ $1/s$ |
|---|---|---|---|---|---|---|---|---|---|---|---|---|---|---|---|---|
| cysRS | 0.24 | 14.4 | 6.00 | 3480 | 6.00 | 3480 | 0.24 | 14.4 | 160 | 400 | 0.01 | 5.0 | 46.0 | 138 | 30.0 | 135 |
| argRS | 2.40 | 36 | 6.00 | 3600 | 6.00 | 3600 | 2.40 | 36 | 270 | 400 | 0.01 | 5.0 | 15.0 | 345 | 40.0 | 184 |
| valRS | 1.00 | 138 | 6.00 | 3000 | 6.00 | 3000 | 1.00 | 138 | 140 | 400 | 0.01 | 5.0 | 80.0 | 30 | 80.0 | 45 |
| ileRS | 2.20 | 15 | 6.00 | 2400 | 6.00 | 2400 | 2.20 | 15 | 150 | 400 | 0.01 | 5.0 | 120 | 138 | 120 | 138 |
| leuRS | 1.20 | 30 | 2.20 | 3630 | 2.20 | 3630 | 1.20 | 30 | 135 | 350 | 0.01 | 5.0 | 15.0 | 189 | 40.0 | 168 |
| gluRS | 0.15 | 27 | 3.40 | 3740 | 3.40 | 3740 | 0.15 | 27 | 140 | 400 | 0.01 | 5.0 | 80.0 | 64 | 80.0 | 64 |
| glnRS | 0.20 | 216 | 5.00 | 3400 | 5.00 | 3400 | 0.20 | 216 | 140 | 400 | 0.01 | 5.0 | 120 | 42 | 120 | 42 |
| tyrRS | 0.60 | 15 | 4.20 | 3630 | 4.20 | 2688 | 0.60 | 12 | 160 | 320 | 0.01 | 5.0 | 42.0 | 48 | 42.0 | 48 |
| trpRS | 0.30 | 13.5 | 8.00 | 2500 | 8.00 | 2500 | 0.30 | 13.5 | 320 | 400 | 0.01 | 5.0 | 45.0 | 117 | 45.0 | 117 |

| Enzyme | $k_{7f}$ $1/s$ | $k_{7b}$ $1/s$ | $k_{8f}$ $1/s$ | $k_{8b}$ $1/s$ | $k_{9f}$ $1/\mu$Ms | $k_{9b}$ $1/s$ | $k_{10f}$ $1/\mu$Ms | $k_{10b}$ $1/s$ | $k_{11f}$ $1/\mu$Ms | $k_{11b}$ $1/s$ | $k^e_{0f}$ $1/\mu$Ms | $k^e_{0b}$ $1/s$ |
|---|---|---|---|---|---|---|---|---|---|---|---|---|
| cysRS | 114 | 400 | 17.3 | 60.7 | 6.00 | 3000 | 1.00 | 2400 | 12.3 | 3.70 | 30.0 | 135 |
| argRS | 340 | 400 | 74.8 | 88 | 6.00 | 3000 | 1.00 | 2400 | 40.0 | 29.0 | 40.0 | 184 |
| valRS | 240 | 400 | 30.7 | 51.2 | 6.00 | 3000 | 1.00 | 2400 | 20.0 | 16.0 | 80.0 | 45 |
| ileRS | 38 | 114 | 7.8 | 23.4 | 6.00 | 3000 | 1.00 | 2400 | 100 | 65.0 | 120 | 138 |
| leuRS | 289 | 340 | 62.9 | 74 | 6.00 | 3000 | 1.00 | 2400 | 10.0 | 15.7 | 40.0 | 168 |
| gluRS | 220 | 400 | 28.0 | 51 | 6.00 | 3000 | 1.00 | 2400 | 30.0 | 8.1 | 80.0 | 64 |
| glnRS | 220 | 400 | 28.0 | 51 | 6.00 | 3000 | 1.00 | 2400 | 12.0 | 7.4 | 120 | 42 |
| tyrRS | 140 | 280 | 32.0 | 64 | 6.00 | 3000 | 1.00 | 2400 | 16.0 | 8.7 | 42.0 | 48 |
| trpRS | 175 | 350 | 44.0 | 88 | 6.00 | 3000 | 1.00 | 2400 | 10.0 | 3.3 | 45.0 | 117 |

**Table C. Kinetic parameters for Class II aminoacyl tRNA synthetase models. Part 1 of 2. Labels of the reactions correspond to those shown in Supplementary Figure D.**

| Enzyme | $k_{1f}$ $1/\mu$Ms | $k_{1b}$ $1/$s | $k_{2f}$ $1/\mu$Ms | $k_{2b}$ $1/$s | $k_{3f}$ $1/\mu$Ms | $k_{3b}$ $1/$s | $k_{4f}$ $1/\mu$Ms | $k_{4b}$ $1/$s | $k_{5f}$ $1/$s | $k_{5b}$ $1/$s | $k_{6f}$ $1/\mu$Ms | $k_{6b}$ $1/$s | $k_{0f}^{a}$ $1/\mu$Ms | $k_{0b}^{a}$ $1/$s | $k_{0f}^{b}$ $1/\mu$Ms | $k_{0b}^{b}$ $1/$s |
|---|---|---|---|---|---|---|---|---|---|---|---|---|---|---|---|---|
| metRS | 1.40 | 70.0 | 4.00 | 3600 | 4.00 | 2880 | 1.40 | 56.0 | 170 | 320 | 6.00 | 3000 | 120 | 84 | 120 | 84 |
| serRS | 0.30 | 18.0 | 6.00 | 3600 | 6.00 | 2880 | 0.30 | 14.4 | 160 | 400 | 6.00 | 3000 | 50.0 | 87.5 | 50.0 | 87.5 |
| thrRS | 0.24 | 60.0 | 6.00 | 3600 | 6.00 | 2250 | 0.24 | 37.5 | 180 | 400 | 6.00 | 3000 | 50.0 | 130 | 50.0 | 130 |
| proRS | 0.25 | 100 | 8.00 | 1600 | 8.00 | 1600 | 0.25 | 100 | 160 | 400 | 6.00 | 3000 | 42.0 | 84 | 42.0 | 84 |
| hisRS | 2.00 | 150 | 3.40 | 3600 | 3.40 | 2880 | 2.00 | 120 | 300 | 500 | 6.00 | 3000 | 90.0 | 150 | 90.0 | 150 |
| aspRS | 0.20 | 174 | 4.60 | 3450 | 4.60 | 3450 | 0.20 | 174 | 60 | 150 | 6.00 | 3000 | 80.0 | 140 | 80.0 | 140 |
| asnRS | 0.24 | 7.2 | 4.60 | 3496 | 4.60 | 3496 | 0.24 | 7.2 | 160 | 360 | 6.00 | 3000 | 40.0 | 48 | 40.0 | 48 |
| lysRS | 2.00 | 180 | 6.00 | 3600 | 6.00 | 2400 | 2.00 | 120 | 100 | 250 | 6.00 | 3000 | 58.0 | 125 | 58.0 | 125 |
| alaRS | 0.70 | 280 | 8.00 | 1600 | 8.00 | 1600 | 0.70 | 280 | 180 | 400 | 6.00 | 3000 | 92.0 | 184 | 92.0 | 184 |
| glyRS | 0.32 | 56.0 | 6.00 | 3600 | 6.00 | 3600 | 0.32 | 56.0 | 160 | 400 | 6.00 | 3000 | 80.0 | 100 | 80.0 | 100 |
| pheRS | 2.00 | 270 | 2.80 | 3600 | 2.80 | 2400 | 2.00 | 180 | 300 | 500 | 6.00 | 3000 | 37.0 | 44 | 37.0 | 44 |

| Enzyme | $k_{1f}^{t}$ $1/\mu$Ms | $k_{1b}^{t}$ $1/$s | $k_{2f}^{t}$ $1/\mu$Ms | $k_{2b}^{t}$ $1/$s | $k_{3f}^{t}$ $1/\mu$Ms | $k_{3b}^{t}$ $1/$s | $k_{4f}^{t}$ $1/\mu$Ms | $k_{4b}^{t}$ $1/$s | $k_{5f}^{t}$ $1/$s | $k_{5b}^{t}$ $1/$s | $k_{6f}^{t}$ $1/\mu$Ms | $k_{6b}^{t}$ $1/$s | $k_{0f}^{c}$ $1/\mu$Ms | $k_{0b}^{c}$ $1/$s | $k_{0f}^{d}$ $1/\mu$Ms | $k_{0b}^{d}$ $1/$s |
|---|---|---|---|---|---|---|---|---|---|---|---|---|---|---|---|---|
| metRS | 1.40 | 70.0 | 4.00 | 3600 | 4.00 | 2880 | 1.40 | 56.0 | 170 | 320 | 6.00 | 3000 | 120 | 84 | 120 | 84 |
| serRS | 0.30 | 18.0 | 6.00 | 3600 | 6.00 | 2880 | 0.30 | 14.4 | 25.0 | 12.5 | 0.01 | 5.0 | 50.0 | 87.5 | 50.0 | 87.5 |
| thrRS | 0.24 | 60.0 | 6.00 | 3600 | 6.00 | 2250 | 0.24 | 37.5 | 29.0 | 19.0 | 6.00 | 3000 | 50.0 | 130 | 50.0 | 130 |
| proRS | 0.25 | 100 | 8.00 | 1600 | 8.00 | 1600 | 0.25 | 100 | 16.0 | 12.0 | 6.00 | 3000 | 42.0 | 84 | 42.0 | 84 |
| hisRS | 2.00 | 150 | 3.40 | 3600 | 3.40 | 2880 | 2.00 | 120 | 19.0 | 9.60 | 6.00 | 3000 | 90.0 | 150 | 90.0 | 150 |
| aspRS | 0.20 | 174 | 4.60 | 3450 | 4.60 | 3450 | 0.20 | 174 | 40.0 | 20.0 | 6.00 | 3000 | 80.0 | 140 | 80.0 | 140 |
| asnRS | 0.24 | 7.2 | 4.60 | 3496 | 4.60 | 3496 | 0.24 | 7.2 | 16.0 | 12.0 | 6.00 | 3000 | 40.0 | 48 | 40.0 | 48 |
| lysRS | 2.00 | 180 | 6.00 | 3600 | 6.00 | 2400 | 2.00 | 120 | 32.0 | 22.4 | 6.00 | 3000 | 58.0 | 125 | 58.0 | 125 |
| alaRS | 0.70 | 280 | 8.00 | 1600 | 8.00 | 1600 | 0.70 | 280 | 22.5 | 22.5 | 6.00 | 3000 | 92.0 | 184 | 92.0 | 184 |
| glyRS | 0.32 | 56.0 | 6.00 | 3600 | 6.00 | 3600 | 0.32 | 56.0 | 70.0 | 35.0 | 6.00 | 3000 | 80.0 | 100 | 80.0 | 100 |
| pheRS | 2.00 | 270 | 2.80 | 3600 | 2.80 | 2400 | 2.00 | 180 | 18.0 | 3.0 | 6.00 | 3000 | 37.0 | 44 | 37.0 | 44 |

**Table D. Kinetic parameters for Class II aminoacyl tRNA synthetase models. Part 2 of 2. Labels of the reactions correspond to those shown in Supplementary Figure D.**

| Enzyme | $k_{7f}$ 1/s | $k_{7b}$ 1/s | $k_{8f}$ 1/s | $k_{8b}$ 1/s | $k_{9f}$ 1/$\mu$Ms | $k_{9b}$ 1/s | $k_{10f}$ 1/$\mu$Ms | $k_{10b}$ 1/s | $k_{11f}$ 1/$\mu$Ms | $k_{11b}$ 1/s | $k_{12f}$ 1/$\mu$Ms | $k_{12b}$ 1/s | $k^e_{0f}$ 1/$\mu$Ms | $k^e_{0b}$ 1/s | $k^f_{0f}$ 1/$\mu$Ms | $k^f_{0b}$ 1/s |
|---|---|---|---|---|---|---|---|---|---|---|---|---|---|---|---|---|
| metRS | 135 | 243 | 42.0 | 75.6 | 6.00 | 3000 | 1.00 | 2400 | 2.30 | 50.6 | 6.00 | 1800 | 120 | 84 | 120 | 84 |
| serRS | 180 | 360 | 26.0 | 52.0 | 6.00 | 3000 | 1.00 | 2400 | 0.30 | 18.0 | 6.00 | 360 | 60.0 | 21 | 50.0 | 87.5 |
| thrRS | 180 | 306 | 17.6 | 30.0 | 6.00 | 3000 | 1.00 | 2400 | 2.40 | 144 | 6.00 | 2100 | 50.0 | 38 | 50.0 | 38 |
| proRS | 80 | 240 | 15.0 | 45.0 | 6.00 | 3000 | 1.00 | 2400 | 0.25 | 70.0 | 8.00 | 1600 | 42.0 | 25.2 | 42.0 | 25.2 |
| hisRS | 115 | 383 | 20.0 | 67.0 | 6.00 | 3000 | 1.00 | 2400 | 4.00 | 70.0 | 6.00 | 2400 | 90.0 | 45 | 90.0 | 45 |
| aspRS | 180 | 360 | 25.0 | 50.0 | 6.00 | 3000 | 1.00 | 2400 | 0.80 | 51.2 | 4.60 | 1470 | 80.0 | 28 | 80.0 | 28 |
| asnRS | 120 | 360 | 15.5 | 46.5 | 6.00 | 3000 | 1.00 | 2400 | 0.24 | 7.2 | 6.00 | 1500 | 40.0 | 16 | 40.0 | 16 |
| lysRS | 160 | 320 | 30.0 | 60.0 | 6.00 | 3000 | 1.00 | 2400 | 4.00 | 64.0 | 6.00 | 240 | 58.0 | 35 | 58.0 | 35 |
| alaRS | 140 | 350 | 22.3 | 55.8 | 6.00 | 3000 | 1.00 | 2400 | 0.70 | 84.0 | 8.00 | 1600 | 92.0 | 83 | 92.0 | 83 |
| glyRS | 180 | 360 | 60.0 | 120 | 6.00 | 3000 | 1.00 | 2400 | 0.32 | 56.0 | 6.00 | 600 | 80.0 | 20 | 80.0 | 20 |
| pheRS | 120 | 360 | 18.0 | 54.0 | 6.00 | 3000 | 1.00 | 2400 | 3.00 | 39.0 | 6.00 | 1800 | 37.0 | 4.4 | 37.0 | 4.4 |

| Enzyme | $k_{13f}$ 1/$\mu$Ms | $k_{13b}$ 1/s | $k_{14f}$ 1/$\mu$Ms | $k_{14b}$ 1/s | $k_{15f}$ 1/s | $k_{15b}$ 1/s | $k_{16f}$ 1/$\mu$Ms | $k_{16b}$ 1/s | $k_{17f}$ 1/$\mu$Ms | $k_{17b}$ 1/s | $k_{18f}$ 1/$\mu$Ms | $k_{18b}$ 1/s |
|---|---|---|---|---|---|---|---|---|---|---|---|---|
| metRS | 6.00 | 1800 | 2.30 | 50.6 | 67.0 | 10.0 | 6.00 | 3000 | 12.0 | 20 | 12.0 | 20 |
| serRS | 6.00 | 360 | 0.30 | 18.0 | 25.0 | 12.5 | 6.00 | 3000 | 50.0 | 50 | 50.0 | 50 |
| thrRS | 6.00 | 2100 | 2.40 | 144 | 39.0 | 19.0 | 6.00 | 3000 | 50.0 | 70 | 50.0 | 70 |
| proRS | 8.00 | 1600 | 0.25 | 70.0 | 16.0 | 12.0 | 6.00 | 3000 | 42.0 | 84 | 42.0 | 84 |
| hisRS | 6.00 | 2400 | 4.00 | 70.0 | 30.0 | 15.0 | 6.00 | 3000 | 50.0 | 60 | 50.0 | 60 |
| aspRS | 4.60 | 1470 | 0.80 | 51.2 | 40.0 | 20.0 | 6.00 | 3000 | 80.0 | 140 | 80.0 | 140 |
| asnRS | 6.00 | 1500 | 0.24 | 7.2 | 16.0 | 12.0 | 6.00 | 3000 | 40.0 | 40 | 40.0 | 40 |
| lysRS | 6.00 | 240 | 4.00 | 64.0 | 64.0 | 16.0 | 6.00 | 3000 | 58.0 | 125 | 58.0 | 125 |
| alaRS | 8.00 | 1600 | 0.70 | 84.0 | 60.0 | 22.5 | 6.00 | 3000 | 92.0 | 230 | 92.0 | 230 |
| glyRS | 6.00 | 600 | 0.32 | 56.0 | 70.0 | 35.0 | 6.00 | 3000 | 120 | 240 | 120 | 240 |
| pheRS | 6.00 | 1800 | 3.00 | 39.0 | 18.0 | 3.0 | 6.00 | 3000 | 40.0 | 80 | 40.0 | 80 |

**Table E. Estimates of AARS Activity *in vivo*. For each of the AARS enzymes, the activity (i.e. tRNA turnover rate $r$) of a single enzyme is estimated in *E. coli* cells growing at $\mu = 0.69$ h$^{-1}$ based on amino acid usage and the number of AARS. Amino acid usage for the Mori *et al.* data (11) was calculated computationally using a model of *in vivo* translation (19) and the fractional amount of tRNAs in ternary complex needed for optimal translation, while the amino acid usage for Jakubowski was taken from their experimental amino acid radio-labelling measurements (9).**

| | | Mori et al.(11) | | | Jakubowski et al. (9) | | |
|---|---|---|---|---|---|---|---|
| Protein | Gene | N. Pro. | AA Use (s$^{-1}$) | $r$ (s$^{-1}$) | N. Enz. | AA Use (s$^{-1}$) | $r$ (s$^{-1}$) |
| cysRS | cysS | 826 | 1606 | 1.94 | | 4600 | |
| argRS | argS | 916 | 14491 | 17.75 | 691 | 6770 | 9.79 |
| valRS | valS | 1553 | 19407 | 12.49 | 536 | 14050 | 26.21 |
| ileRS | ileS | 2061 | 13844 | 6.71 | 1226 | 15270 | 13.45 |
| leuRS | leuS | 1903 | 18583 | 9.76 | 875 | 20250 | 23.14 |
| gluRS | gltX | 2194 | 15534 | 7.08 | 873 | 1730 | 1.98 |
| glnRS | glnS | 1518 | 8337 | 5.49 | 1013 | 4980 | 4.91 |
| metRS | metG | 1732 | 6342 | 7.32 | | 7630 | |
| tyrRS | tyrS | 1686 | 5486 | 6.50 | | 1920 | |
| trpRS | trpS | 2475 | 1704 | 1.37 | | 3150 | |
| serRS | serS | 1725 | 11066 | 12.83 | | 51090 | |
| thrRS | thrS | 2807 | 12120 | 8.63 | 527 | 25210 | 47.84 |
| proRS | proS | 1969 | 8501 | 8.63 | | 10030 | |
| hisRS | hisS | 934 | 4324 | 9.25 | | 4030 | |
| aspRS | aspS | 1636 | 12104 | 14.79 | | 15930 | |
| asnRS | asnS | 3604 | 8424 | 4.67 | | 2100 | |
| lysRS | lysS | 2435 | 16038 | 13.17 | 425 | 11850 | 27.88 |
| alaRS | alaS | 1627 | 22820 | 28.05 | | 13390 | |
| lysRS* | lysU | 278 | | | | | |
| glyRS$\alpha$ | glyS | 1744 | 18848 | 21.61 | 682 | 10210 | 14.97 |
| glyRS$\beta$ | glyQ | 1614 | 18848 | 23.35 | | | |
| pheRS$\alpha$ | pheS | 1319 | 7519 | 11.40 | 992 | 9950 | 10.03 |
| pheRS$\beta$ | pheT | 1819 | 7519 | 8.26 | | | |

**Table F. Average Number of Proteins, tRNA, and Ribosomes in *E. coli* Cells at Different Growth Rates. Protein numbers are taken from proteome measurements from several groups (11–13) and reported here as number per cell. The specific data for each growth rate are; $\mu = 0.41$ h$^{-1}$ Valgepea *et al.* 2013 (12), $\mu = 0.69$ h$^{-1}$ and $\mu = 1.04$ h$^{-1}$ Mori *et al.* 2021 (11), $\mu = 1.98$ h$^{-1}$ Li *et al.* 2014 (13). The number of tRNA and Ribosomes (per cell) are taken from Dennis and Bremer (16). The doubling time in minutes is computed from the growth rate using $\tau = 60\ln(2)/\mu$.**

| Protein | Gene | $\mu = 0.41$ h$^{-1}$ $\tau = 100$ | $\mu = 0.69$ h$^{-1}$ $\tau = 60$ | $\mu = 1.04$ h$^{-1}$ $\tau = 40$ | $\mu = 1.98$ h$^{-1}$ $\tau = 21$ |
|---|---|---|---|---|---|
| Ribo | | 8000 | 15000 | 26000 | 73000 |
| tRNA | | 74000 | 139000 | 241000 | 680000 |
| cysRS | cysS | 609 | 826 | 1409 | 4958 |
| argRS | argS | 733 | 916 | 1457 | 4901 |
| valRS | valS | 1179 | 1553 | 2502 | 8921 |
| ileRS | ileS | 2135 | 2061 | 2832 | 6143 |
| leuRS | leuS | 2706 | 1903 | 2534 | 8069 |
| gluRS | gltX | 2751 | 2194 | 2980 | 13429 |
| glnRS | glnS | 2076 | 1518 | 2113 | 8204 |
| metRS | metG | 1618 | 1732 | 2415 | 6555 |
| tyrRS | tyrS | 1447 | 1686 | 3661 | 7986 |
| trpRS | trpS | 1975 | 2475 | 3764 | 7805 |
| serRS | serS | 2948 | 1725 | 3241 | 11805 |
| thrRS | thrS | 1222 | 2807 | 4091 | 12096 |
| proRS | proS | 4060 | 1969 | 2415 | 12156 |
| hisRS | hisS | 1438 | 934 | 1956 | 3774 |
| aspRS | aspS | 2513 | 1636 | 2993 | 8307 |
| asnRS | asnS | 4035 | 3604 | 5615 | 17436 |
| lysRS | lysS | 4498 | 2435 | 3204 | 15890 |
| alaRS | alaS | 2000 | 1627 | 2962 | 8493 |
| lysRS* | lysU | 117 | 278 | 372 | 1644 |
| glyRS$\alpha$ | glyS | 2871 | 1744 | 2499 | 12974 |
| glyRS$\beta$ | glyQ | 2313 | 1614 | 2067 | 12365 |
| pheRS$\alpha$ | pheS | | 1319 | 2443 | 8935 |
| pheRS$\beta$ | pheT | 1579 | 1819 | 3324 | 8573 |
| EfTu | tufA | 39582 | 51042 | 68208 | 289537 |
| EfTu | tufB | 39581 | 51105 | 68292 | 289537 |
| EfTs | tfs | 16181 | 15894 | 26017 | 95122 |
| EfG | fusA | 18213 | 15065 | 25810 | 84151 |
| Inf1 | infA | 4843 | 5694 | 9220 | 30461 |
| Inf2 | infB | 2856 | 2493 | 4082 | 13806 |
| Inf3 | infC | 4876 | 4765 | 7890 | 22624 |
| RRF | frr | 5420 | 5237 | 7112 | 23316 |
| RF1 | prfA | | 184 | 341 | 1059 |
| RF2 | prfB | | 108 | 110 | 10031 |
| RF3 | prfC | 614 | 874 | 1088 | 7808 |

**Table G. Average number of tRNAs per cell and their codon recognition. Data for the number of tRNAs that are in free ternary complex at a growth rate of $\mu = 0.41$ h$^{-1}$ ($\tau = 100$ min). The numbers that give the optimal translation rate have been computed from Eq. 3 and the procedure outlined in section 4, and the total tRNA numbers are taken from Dong *et al.* (24). The values used in the model are the result of the fitting procedure in section 7.**

| | | Ternary Complex | | | Total tRNA | | |
|---|---|---|---|---|---|---|---|
| tRNA | Codon Rec. | Opt. Trans. | Model | Error | Dong | Model | Error |
| Lys | AAA,AAG | 1424 | 1429 | 0.00 | 2264 | 3210 | 0.41 |
| Asn | AAC,AAU | 1028 | 914 | 0.12 | 1285 | 2370 | 0.84 |
| Thr4 | ACA,ACU,ACG | 629 | 675 | 0.07 | 1055 | 1216 | 0.15 |
| Thr1+3 | ACC,ACU | 728 | 541 | 0.34 | 1378 | 1552 | 0.12 |
| Thr2 | ACG | 353 | 449 | 0.27 | 666 | 541 | 0.18 |
| Arg4 | AGA | 143 | 246 | 0.72 | 782 | 293 | 0.62 |
| Ser3 | AGC,AGU | 641 | 641 | 0.00 | 1462 | 1224 | 0.16 |
| Arg5 | AGG | 72 | 147 | 1.04 | 522 | 165 | 0.68 |
| Ile2 | AUA | 139 | 271 | 0.94 | 224 | 320 | 0.42 |
| Ile1 | AUC,AUU | 1319 | 1337 | 0.01 | 3722 | 3919 | 0.05 |
| MetM | AUG | 813 | 795 | 0.02 | 862 | 1540 | 0.78 |
| Gln1 | CAA | 471 | 1156 | 1.45 | 905 | 1540 | 0.70 |
| His | CAC,CAU | 739 | 536 | 0.37 | 729 | 1166 | 0.60 |
| Gln2 | CAG | 904 | 660 | 0.37 | 1025 | 1720 | 0.67 |
| Pro3 | CCA,CCU,CCG | 393 | 420 | 0.07 | 629 | 1166 | 0.85 |
| Pro2 | CCC,CCU | 495 | 509 | 0.03 | 836 | 708 | 0.15 |
| Pro1 | CCG | 524 | 560 | 0.07 | 812 | 1165 | 0.43 |
| Arg3 | CGG | 138 | 1067 | 6.73 | 869 | 1166 | 0.34 |
| Arg2 | CGU,CGC,CGA | 1338 | 2150 | 0.61 | 4842 | 4896 | 0.01 |
| Leu3 | CUA,CUG | 179 | 320 | 0.79 | 829 | 680 | 0.17 |
| Leu2 | CUC,CUU | 545 | 550 | 0.01 | 1155 | 980 | 0.15 |
| Leu1 | CUG | 1206 | 2080 | 0.72 | 4966 | 4322 | 0.13 |
| Glu2 | GAA,GAG | 1403 | 1488 | 0.06 | 5189 | 4206 | 0.18 |
| Asp1 | GAC,GAU | 1236 | 1223 | 0.01 | 2707 | 2586 | 0.04 |
| Ala1B | GCU,GCA,GCG | 1586 | 1607 | 0.01 | 3906 | 3919 | 0.01 |
| Ala2 | GCC | 597 | 490 | 0.22 | 706 | 810 | 0.15 |
| Gly2 | GGA,GGG | 279 | 1008 | 2.61 | 1055 | 1216 | 0.15 |
| Gly3 | GGC,GGU | 1493 | 1540 | 0.03 | 5066 | 3981 | 0.21 |
| Gly1a | GGG | 163 | 705 | 3.32 | 1336 | 800 | 0.40 |
| Val1 | GUA,GUG,GUU | 1364 | 1510 | 0.11 | 4020 | 3404 | 0.15 |
| Val2ab | GUC,GUU | 524 | 574 | 0.09 | 1462 | 1165 | 0.20 |
| Tyr1+2 | UAC,UAU | 834 | 834 | 0.00 | 2087 | 2110 | 0.01 |
| Ser1 | UCA,UCU,UCG | 818 | 763 | 0.07 | 1851 | 1420 | 0.23 |
| Ser5 | UCC,UCU | 416 | 394 | 0.06 | 865 | 1112 | 0.28 |
| Ser2 | UCG | 160 | 314 | 0.96 | 346 | 367 | 0.06 |
| Cys | UGC,UGU | 451 | 498 | 0.10 | 1625 | 1315 | 0.19 |
| Trp | UGG | 463 | 496 | 0.07 | 925 | 943 | 0.02 |
| Leu5 | UUA,UUG | 336 | 1033 | 2.07 | 1155 | 1315 | 0.14 |
| Phe | UUC,UUU | 976 | 976 | 0.00 | 1199 | 2134 | 0.78 |
| Leu4 | UUG | 650 | 1512 | 1.32 | 2108 | 1744 | 0.17 |

**Table H. Concentrations of Amino Acids in *E. coli*. Experimental measurements of intracellular amino acid concentrations are shown for Bennett *et al.* (4), and minimal media (MM) and minimal media plus amino acids (MM+AA) from Avcilar-Kucukgoze (34). The values used in the model for the MM and MM+AA scenarios are shown in the final two columns. *(Combined concentration of ile+leu was measured to be 300 $\mu$M). (nd = not determined)**

| | Bennett | Avcilar-Kucukgoze | | Model | |
|---|---|---|---|---|---|
| AA | | MM | MM+AA | MM | MM+AA |
| cys | nd | nd | nd | 30 | 300 |
| arg | 570 | nd | nd | 570 | 570 |
| val | 4200 | 185 | 3000 | 4200 | 4200 |
| ile | 80* | 9 | 290 | 80 | 300 |
| leu | 220* | nd | nd | 220 | 360 |
| glu | 96000 | nd | nd | 96000 | 96000 |
| gln | 3800 | nd | 210 | 3800 | 3800 |
| met | 150 | nd | nd | 150 | 160 |
| tyr | 29 | 11 | 210 | 29 | 270 |
| trp | 12 | 65 | 770 | 20 | 220 |
| ser | 68 | 57 | 4000 | 68 | 4000 |
| thr | 180 | 8 | 340 | 180 | 450 |
| pro | 390 | nd | nd | 390 | 2850 |
| his | 68 | 8 | 130 | 68 | 160 |
| asp | 4200 | nd | nd | 4200 | 4200 |
| asn | 510 | 2 | 230 | 510 | 510 |
| lys | 410 | 24 | 250 | 410 | 410 |
| ala | 2600 | 27 | 1210 | 2600 | 2600 |
| gly | 800 | 62 | 360 | 800 | 800 |
| phe | 18 | 12 | 270 | 20 | 270 |

**Table I. Optimized numbers of Aminoacyl tRNA Synthetases in Exponentially Growing *E. coli* and optimized $k_{cat}$ and $K_m$ values. The optimized average number of AARS enzymes are shown for different growth rates, $\mu = 0.41$ h$^{-1}$ ($\tau = 100$ min), $\mu = 0.69$ h$^{-1}$ ($\tau = 60$ min), $\mu = 1.04$ h$^{-1}$ ($\tau = 40$ min), and $\mu = 1.73$ h$^{-1}$ ($\tau = 24$ min). Corresponding optimized $k_{cat}$ and Michaelis–Menten parameters for tRNA ($K_m^t$) and amino acid ($K_m^a$) are given in the right hand columns.**

| | Optimized Numbers | | | | Optimized Kcat/Km | | |
|---|---|---|---|---|---|---|---|
| AARS | $\mu = 0.41$ | $\mu = 0.69$ | $\mu = 1.04$ | $\mu = 1.73$ | $k_{cat}$ (s$^{-1}$) | $K_m^t (\mu$M) | $K_m^a (\mu$M) |
| cysRS | 440 | 895 | 1610 | 3800 | 2.9 | 1.05 | 16.1 |
| argRS | 450 | 857 | 1540 | 3400 | 20.2 | 1.76 | 15.1 |
| valRS | 1050 | 2100 | 3675 | 8140 | 10.6 | 0.41 | 42.0 |
| ileRS | 1140 | 2240 | 3945 | 9000 | 7.2 | 1.18 | 5.8 |
| leuRS | 1050 | 2115 | 3870 | 8900 | 11.2 | 1.17 | 15.6 |
| gluRS | 1350 | 2750 | 4860 | 10800 | 6.2 | 0.27 | 72.3 |
| glnRS | 850 | 1800 | 3265 | 7520 | 5.8 | 0.16 | 188 |
| metRS | 480 | 1020 | 1860 | 4180 | 10.7 | 0.75 | 8.1 |
| tyrRS | 500 | 1030 | 1840 | 4265 | 6.6 | 0.46 | 14.2 |
| trpRS | 550 | 1210 | 2260 | 5380 | 3.1 | 0.29 | 11.4 |
| serRS | 780 | 1640 | 3030 | 6800 | 8.7 | 0.39 | 56.0 |
| thrRS | 800 | 1600 | 2890 | 6510 | 9.6 | 0.72 | 24.1 |
| proRS | 790 | 1530 | 2750 | 6400 | 6.7 | 0.48 | 149 |
| hisRS | 400 | 800 | 1450 | 3330 | 9.2 | 0.41 | 7.9 |
| aspRS | 600 | 1200 | 2050 | 4600 | 12.5 | 0.47 | 38.8 |
| asnRS | 960 | 2015 | 3660 | 8260 | 5.8 | 0.52 | 34.6 |
| lysRS | 890 | 1450 | 2575 | 5690 | 16.3 | 0.94 | 8.8 |
| alaRS | 1000 | 1800 | 3260 | 7300 | 13.2 | 0.91 | 47.1 |
| glyRS | 480 | 930 | 1660 | 3650 | 24.7 | 0.68 | 140 |
| pheRS | 610 | 1250 | 2280 | 5185 | 7.7 | 0.35 | 9.1 |

**Table J. Optimized numbers of tRNAs per cell and their codon recognition. The optimized total tRNA numbers are shown for the Michaelis–Menten model and the full kinetic model where AARS enzymes are modelled as having reaction scheme according to Figure C for Class I enzymes or Figure D for class II enzymes.**

| | | Michaelis–Menten | | | | Full Model | | | |
|---|---|---|---|---|---|---|---|---|---|
| tRNA | Codon Rec. | $\mu = 0.41$ | $\mu = 0.69$ | $\mu = 1.04$ | $\mu = 1.73$ | $\mu = 0.41$ | $\mu = 0.69$ | $\mu = 1.04$ | $\mu = 1.73$ |
| Lys | AAA,AAG | 2759 | 5575 | 10627 | 23872 | 3210 | 6322 | 10980 | 23119 |
| Asn | AAC,AAU | 1805 | 3384 | 6509 | 14766 | 2370 | 4675 | 8324 | 18084 |
| Thr4 | ACA,ACU,ACG | 1113 | 2200 | 4046 | 9199 | 1216 | 2399 | 4187 | 9108 |
| Thr1+3 | ACC,ACU | 1348 | 2538 | 4532 | 10212 | 1552 | 2988 | 4979 | 10548 |
| Thr2 | ACG | 533 | 1077 | 2076 | 4757 | 541 | 1107 | 2016 | 4539 |
| Arg4 | AGA | 288 | 564 | 1190 | 2730 | 293 | 578 | 1153 | 2604 |
| Ser3 | AGC,AGU | 961 | 1846 | 3309 | 7541 | 1224 | 2414 | 4213 | 9327 |
| Arg5 | AGG | 165 | 323 | 689 | 1580 | 165 | 325 | 660 | 1496 |
| Ile2 | AUA | 301 | 620 | 1142 | 2708 | 320 | 661 | 1154 | 2679 |
| Ile1 | AUC,AUU | 2297 | 4565 | 9050 | 21003 | 3919 | 7731 | 13310 | 28748 |
| MetF | AUG | 2364 | 5132 | 7809 | 21870 | 2364 | 5132 | 7809 | 21870 |
| MetM | AUG | 1366 | 2618 | 4953 | 11230 | 1540 | 3078 | 5372 | 11729 |
| Gln1 | CAA | 1377 | 2764 | 5089 | 11410 | 1540 | 3078 | 5372 | 11729 |
| His | CAC,CAU | 936 | 1829 | 3556 | 8237 | 1166 | 2300 | 4014 | 8881 |
| Gln2 | CAG | 1155 | 2254 | 4448 | 10142 | 1720 | 3433 | 5992 | 13100 |
| Pro3 | CCA,CCU,CCG | 862 | 1753 | 3317 | 7643 | 1166 | 2300 | 4014 | 8881 |
| Pro2 | CCC,CCU | 648 | 1342 | 2644 | 6038 | 708 | 1450 | 2702 | 6042 |
| Pro1 | CCG | 935 | 1880 | 3608 | 8337 | 1165 | 2316 | 4131 | 9173 |
| Arg3 | CGG | 1186 | 2280 | 4167 | 9398 | 1166 | 2300 | 4014 | 8881 |
| Arg2 | CGU,CGC,CGA | 4475 | 8880 | 16261 | 37500 | 4896 | 9656 | 16852 | 37308 |
| Leu3 | CUA,CUG | 544 | 1078 | 2083 | 4665 | 680 | 1361 | 2375 | 5179 |
| Leu2 | CUC,CUU | 824 | 1671 | 3277 | 7402 | 980 | 2012 | 3544 | 7838 |
| Leu1 | CUG | 3441 | 6740 | 12992 | 28963 | 4322 | 8506 | 14780 | 31841 |
| Glu2 | GAA,GAG | 2829 | 5483 | 10707 | 24536 | 4206 | 8291 | 14471 | 32464 |
| Asp1 | GAC,GAU | 2294 | 4500 | 8982 | 20966 | 2586 | 5104 | 8908 | 19696 |
| Ala1B | GCU,GCA,GCG | 3689 | 8508 | 15144 | 35209 | 3919 | 7731 | 13310 | 28748 |
| Ala2 | GCC | 787 | 1775 | 3429 | 8008 | 810 | 1689 | 3016 | 6747 |
| Gly2 | GGA,GGG | 1181 | 2354 | 4312 | 9632 | 1216 | 2399 | 4187 | 9108 |
| Gly3 | GGC,GGU | 3456 | 7184 | 13180 | 30997 | 3981 | 7853 | 13706 | 30817 |
| Gly1a | GGG | 790 | 1565 | 2841 | 6329 | 800 | 1578 | 2754 | 5983 |
| Val1 | GUA,GUG,GUU | 2780 | 5387 | 10578 | 24202 | 3404 | 6700 | 11694 | 25926 |
| Val2ab | GUC,GUU | 969 | 1931 | 3767 | 8627 | 1165 | 2324 | 4056 | 8873 |
| Tyr1+2 | UAC,UAU | 1586 | 3060 | 5942 | 13225 | 2110 | 4154 | 7250 | 15620 |
| Ser1 | UCA,UCU,UCG | 1109 | 2187 | 3915 | 8900 | 1420 | 2808 | 4850 | 10711 |
| Ser5 | UCC,UCU | 747 | 1417 | 2483 | 5637 | 1112 | 2113 | 3641 | 7884 |
| Ser2 | UCG | 349 | 732 | 1297 | 2970 | 367 | 760 | 1307 | 2952 |
| Cys | UGC,UGU | 991 | 1903 | 3499 | 7801 | 1315 | 2594 | 4528 | 9996 |
| Trp | UGG | 689 | 1366 | 2694 | 6200 | 943 | 1920 | 3536 | 7977 |
| Leu5 | UUA,UUG | 1231 | 2386 | 4470 | 10048 | 1315 | 2594 | 4528 | 9996 |
| Phe | UUC,UUU | 1693 | 3352 | 6450 | 14677 | 2134 | 4210 | 7320 | 15926 |
| Leu4 | UUG | 1695 | 3335 | 6113 | 13796 | 1744 | 3440 | 6004 | 13283 |

**Table K. Model predictions of AARS Activity *in vivo*. For each of the AARS enzymes, the activity (i.e. tRNA turnover rate) of a single enzyme $r$ per second is estimated in *E. coli* cells growing at $\mu = 1.04$ h$^{-1}$ ($\tau = 40$ min) based on amino acid usage and the number of AARS. Amino acid usage for the data in this work was calculated computationally using a model of *in vivo* translation (19), while the amino acid usage and AARS numbers for Choi and Covert were taken from supplementary material Tables 8 and S3 (10).**

| | This work | | | Choi and Covert (10) | | |
|---|---|---|---|---|---|---|
| AARS | N. Enz. | AA Use (s$^{-1}$) | $r$ (s$^{-1}$) | N. Enz. | AA Use (s$^{-1}$) | $r$ (s$^{-1}$) |
| cysRS | 1610 | 2714.8 | 1.68 | 606 | 2696.6 | 4.44 |
| argRS | 1540 | 24209.7 | 15.72 | 625 | 20752.4 | 33.20 |
| valRS | 3675 | 32156.5 | 8.75 | 1625 | 28900.7 | 17.78 |
| ileRS | 3945 | 23078.4 | 5.85 | 1698 | 20723.5 | 12.20 |
| leuRS | 3870 | 31360.8 | 8.10 | 1278 | 30571.2 | 23.92 |
| gluRS | 4860 | 25845.7 | 5.31 | 2864 | 24502.9 | 8.55 |
| glnRS | 3265 | 13962.0 | 4.27 | 1310 | 14129.4 | 10.78 |
| metRS | 1860 | 10548.6 | 5.67 | 843 | 9434.0 | 11.19 |
| tyrRS | 1840 | 9173.5 | 4.98 | 994 | 9792.9 | 9.85 |
| trpRS | 2260 | 2939.7 | 1.30 | 879 | 3383.7 | 3.84 |
| serRS | 3030 | 18529.5 | 6.11 | 1438 | 18586.3 | 12.92 |
| thrRS | 2890 | 20132.9 | 6.96 | 1824 | 20426.6 | 11.19 |
| proRS | 2750 | 14275.0 | 5.19 | 862 | 13473.6 | 15.63 |
| hisRS | 1450 | 7219.3 | 4.97 | 427 | 6981.9 | 16.35 |
| aspRS | 2050 | 20082.0 | 9.79 | 567 | 20693.3 | 36.49 |
| asnRS | 3660 | 14055.5 | 3.84 | 2140 | 15273.5 | 7.13 |
| lysRS | 2575 | 26206.4 | 10.17 | 1410 | 24330.0 | 17.25 |
| alaRS | 3260 | 37879.9 | 11.61 | 689 | 36929.9 | 53.59 |
| glyRS | 1660 | 31119.9 | 18.74 | 804 | 29038.0 | 36.11 |
| pheRS | 2280 | 12634.3 | 5.54 | 773 | 12144.5 | 15.71 |
